# Afternoon to early evening bright light exposure reduces later melatonin production in adolescents

**DOI:** 10.1101/2024.10.02.616112

**Authors:** Rafael Lazar, Fatemeh Fazlali, Marine Dourte, Christian Epple, Oliver Stefani, Manuel Spitschan, Christian Cajochen

**Affiliations:** Centre for Chronobiology, Psychiatric University Clinics (UPK) Basel, Basel, Switzerland; Research Cluster Molecular and Cognitive Neurosciences, University of Basel, Basel, Switzerland; Sleep and Chronobiology Laboratory, GIGA-CRC Human Imaging Unit, University of Liège, Liège, Belgium; UR2NF, Neuropsychology and Functional Neuroimaging Research Unit, Center for Research in Cognition and Neurosciences, Neurosciences Institute, Université Libre de Bruxelles (ULB), Brussels, Belgium; Lucerne University of Applied Sciences and Arts, Horw, Switzerland; Translational Sensory & Circadian Neuroscience, Max Planck Institute for Biological Cybernetics, Tübingen, Germany; Chronobiology & Health, TUM School of Medicine and Health (TUM MH), Technical University of Munich, Munich, Germany

## Abstract

Whether light exposure during the day reduces non-visual light effects later in the evening has not been studied in adolescents. We investigate whether afternoon-early evening (AEE) light interventions (130 lx, 2500 lx, 4.5 hours, compared to 6.5 lx) would increase melatonin levels during later evening light exposure (130 lx) in a counterbalanced crossover study with 22 adolescents (14-17 years, 11 female). Contrary to our hypothesis, evening melatonin levels decreased after AEE bright light exposure, while sleepiness and vigilance were unaffected, and skin temperature showed no clear changes. The AEE light had acute alerting effects and bright light exposure in the 32 hours before laboratory entry was associated with higher evening melatonin and sleepiness. These findings suggest that bright AEE light increases alertness but may delay melatonin production by interfering with circadian rhythms. The study highlights the complex effects of light timing and its implications for managing adolescents’ light exposure.

## Introduction

Adolescents are highly susceptible to chronic sleep loss on school days^1^, with empirical data suggesting that between 30 and 70% of them suffer from inadequate sleep^1,2^. Extensive research has revealed numerous ramifications associated with adolescents’ sleep deprivation, including mental health issues^3^, increased risk behaviour^4^, and decreased cognitive functioning^5^. While adolescents’ sleep need remains stable across adolescence^6^ (Ø ∼ 9 hours^7,8^), data from several studies indicate that adolescents’ weekday sleep duration decreases as they age (between 10–18 years)^9–11^. This sleep loss is likely associated with a combination of early school/work start times and later bedtimes, often referred to as “social jetlag”^12^, triggered by a delay-shift of the “inner clock” during puberty (see reviews^13–15)^. Physiologically, this delay has been linked to a slower build-up of sleep pressure in late compared to early puberty^16,17^, enabling older adolescents to stay up longer^18^. The physiological “ability” to stay up longer tends to converge with greater bedtime autonomy in older adolescents^19^ facilitating nighttime activities which can have a multifaceted impact on sleep^20,21^. First, engaging content may prolong bedtime and sleep onset latency by increasing arousal^22^. Second, nighttime activities are likely to be combined with light exposure^13^ which can suppress adolescents’ melatonin release, increase alertness, and delay the circadian rhythm significantly^23–26^.

### Light and sleep-wake-regulation

Light is the essential synchroniser for humans’ circadian timing system, entraining the inner clock to the environment and playing a significant role in immunological, cognitive, emotional, and sleep-wake regulation (see reviews^27,28^). Light exposure can delay or advance circadian rhythms profoundly, the direction depending on the individuals’ “biological time” (e.g. time relative to their melatonin onset) at exposure^25,29^. The relationship between the timing of a given light exposure and the resulting direction and magnitude of the phase shift, at different points in the circadian cycle, is typically characterised by the phase response curve (PRC)^29^. Broadly speaking, early morning light advances the circadian phase, and late evening light delays it. The size of the light effect depends on the intensity and wavelength profile of the light^30,31^ and the individuals’ light sensitivity^32^ which in turn can be influenced by prior light history^33^. Brighter light and a higher short-wavelength proportion increase the strength of lights’ impact on the circadian system, particularly in individuals with a higher circadian sensitivity and a lower prior light exposure history. Additionally, light exposure has acute alerting physiological and behavioural effects: bright light can enhance alertness^34,35^, elevate heart rate^31,36^, increase core body temperature (CBT) while decreasing peripheral vasodilation^37,38^, and decrease melatonin release naturally occurring at night^23,39–41^. Converging evidence suggests that these non-visual effects of light in mammals are primarily driven by the intrinsically photosensitive retinal ganglion cells (ipRGCs), expressing the short-wavelength-sensitive photopigment melanopsin^30,40,42–44^. The ipRGCs relay the light information to the “circadian master clock” located in the suprachiasmatic nuclei (SCN) of the hypothalamus via the retinohypothalamic tract (RHT)^45^.

### Light exposure interventions in adolescents

Though triggered naturally during puberty, adolescents’ late sleep-wake rhythms are likely exacerbated by an increasing opportunity for ill-timed light exposure behaviour in older adolescents^13,46^, making light a viable target for circadian interventions. Firstly, a lack of morning light exposure is associated with a later circadian timing in adolescents^47,48^, likely due to reducing phase-advancing parts of light exposure. Consistent with these findings, bright light interventions in the morning have shown potential for circadian phase advancing in adolescents^49–52^, with the limitation that they often have to reduce morning sleep to apply light in the most sensitive phase-advancing hours^50^ or use special brief light flash devices during sleep^49^. Furthermore, several studies have demonstrated that night light exposure can acutely attenuate melatonin release, and have phase delaying effects in adolescents^20,23,25,53,54^ while electronic device use has been associated with decreased sleep quality^55–57^. However, evening restrictions for smartphone use were shown to be challenging to negotiate with adolescents^57^ and excessively bright light in the home environment remains a widespread problem^58,59^. Thus, other ways of reducing the alerting and physiological effects of evening light exposure in adolescents should be explored.

### Prior evidence for circadian photosensitivity adaptation

There is a growing body of literature indicating that the circadian amplitude and circadian photosensitivity can adapt depending on the prior light history. Studies in animal models ^60,61^ in vitro ipRGCs ^62,63^ and adults ^33,64–79^ suggest that increasing prior daytime light exposure may increase the circadian amplitude and decrease the effects of subsequent light exposure at night. This has been demonstrated for the effects of night light exposure on phase-shifting^64,72,73,75^, melatonin secretion^33,65,67,74,78^, and alertness^71^ in adults. However, it is less clear what role the timing, duration and intensity of the prior light exposure play in modulating circadian responses, as many of these studies have applied high-contrast long-duration light conditions over multiple days^33,64–66,68,71,73^. Some studies have demonstrated circadian photosensitivity adaptation after morning light exposure^75,76^, after prior light in the evening^79^ and via light exposure at night^67,77^. However, there seems to be a lack of experimental studies investigating prior light exposure in the afternoon and early evening.

### Afternoon light exposure in adolescents

In general, only few studies have investigated the effects of afternoon light exposure or lack thereof on the circadian rhythms and sleep of adolescents^25,47,48,80,81^. Crowley and Eastman^25^ constructed a PRC based on N=44 adolescents aged 14-17 years, which showed that two hours of bright light exposure (∼5000 lx illuminance) in the afternoon had phase-advancing effects when it lasted up to ∼9 hours before midsleep time (i.e. up to ∼6 pm for an adolescent who habitually goes to bed at ∼10:30 pm and wakes up at ∼7:30 am). Furthermore, data from field studies involving adolescents have shown an association between increased afternoon bright light exposure and earlier sleep onset^81^ or earlier sleep offset the next day^48^.

Although light history has been suggested to influence subsequent circadian photosensitivity^65^, no study to date has systematically characterised how afternoon and early evening light of different intensities might modulate physiological responses to subsequent late evening light exposure in adolescents. The aim of this study was to investigate whether increasing after-school light exposure could be a viable approach to mitigating the alerting effects of artificial light around bedtime in adolescents. We used a counterbalanced crossover study design to test how adolescents’ physiological responses to late evening light exposure depend on prior afternoon to early evening (AEE) light exposure across 3 different light levels. The primary endpoint of this study was salivary melatonin concentration (AUC) during late evening light exposure, and we tested the following hypotheses:

#### A-priori

Evening salivary melatonin levels (AUC), subjective sleepiness (KSS), and distal-to-proximal skin temperature gradient (DPG) are elevated and response speed (PVT) is reduced when preceded by the “moderate” or “bright” AEE light intervention, compared to “dim” light.

#### Post-hoc

Subjective sleepiness (KSS) and the distal-to-proximal skin temperature gradient (DPG) are reduced, and response speed (PVT) is elevated during the “moderate” or “bright” AEE light intervention, compared to “dim” light.

## Results

22 healthy German-speaking adolescents (14-17 years, mean=15.91±1.15 years, 50% female) completed the counterbalanced crossover protocol (see Figure 1), which was adapted to each individual’s habitual bedtime (HBT; mean=22:15, SD=31 min). Table 1 summarises the characteristics of the participants stratified by sex, and Suppl. Table S1 shows the applied inclusion and exclusion criteria. The AEE light intervention stimuli (see Fig. 2A and C) applied between 7.5 and 3 hours before each individual’s HBT had approximately equivalent spectral distributions and differed by ∼1.3 log_10_ units in terms of light level (see Fig. 2B and D). The later evening light exposure stimulus, applied in all sessions between 3 hours before HBT and 1.5 hours after HBT, was equivalent to the “moderate” light intervention. The average time courses of the outcome measures per light condition are shown in Figure 3, while Figure 4 shows the average light history during the day prior to each experiment, expressed as the percentage above threshold per hour at different illuminance thresholds. For thresholds ≥500 lx illuminance, the curves show small peaks throughout the day, likely corresponding to increased access to daylight.

**Figure 1.**
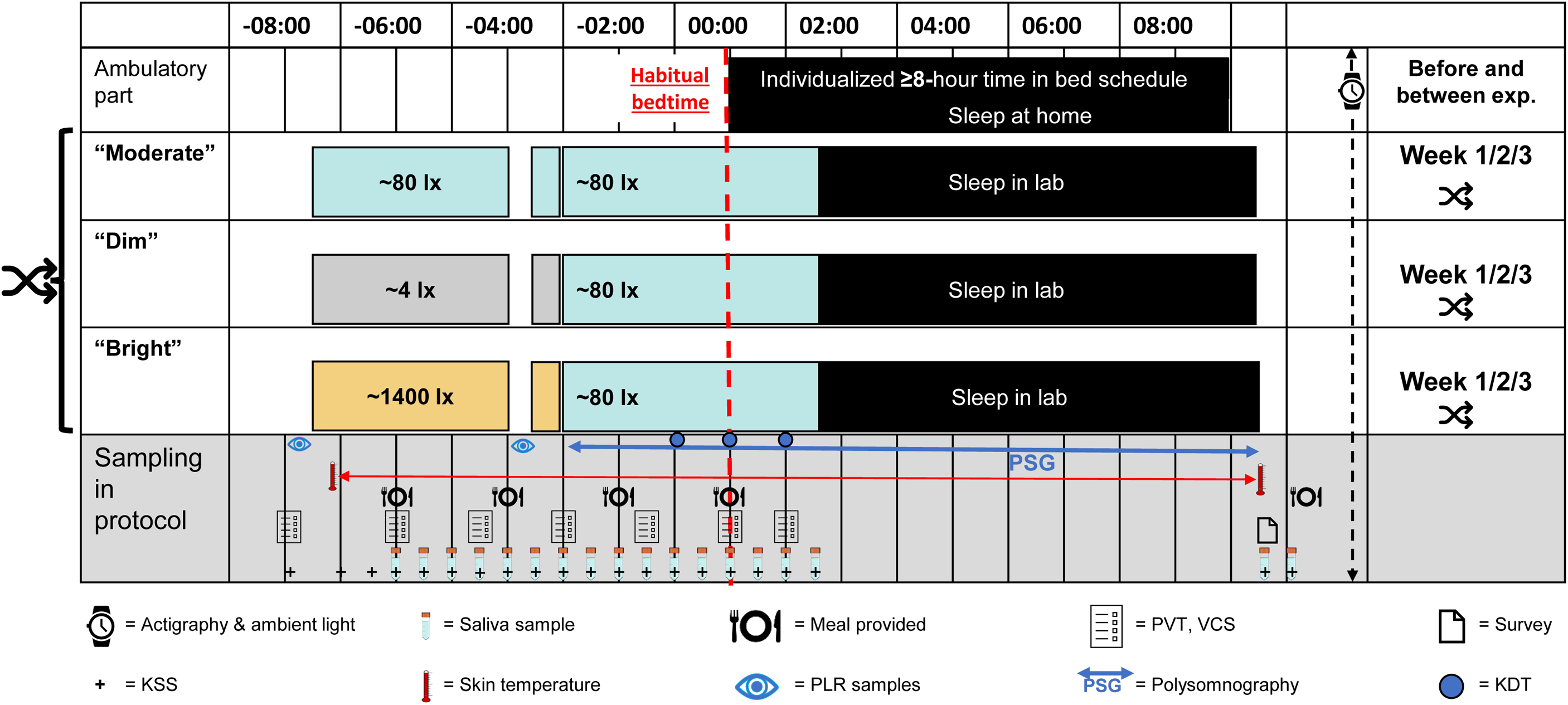
Experimental crossover protocol for assessing the AEE light interventions. Adolescent volunteers participated in a 21-day study schedule that included ≥5 days of sleep-wake stabilisation with sleep and light monitoring before each of the 3 experimental visits, which were individually scheduled to begin 8 h before HBT and last until 10.5 h after HBT. On average, HBT (0:00 in the protocol) corresponded to 22:15 (SD=31 min). AEE light interventions lasted from 7.5 h to 3 h before HBT and varied between “dim”, “moderate” and “bright” light exposure in a counterbalanced order (∼4 lx, ∼80 lx, ∼1400 lx mEDI, respectively), followed by later evening light exposure from 3 h before to 1.5 h after HBT (∼80 lx mEDI). While awake, toilet breaks were scheduled every 1.5 h, and 4 equicaloric meals were provided every 2 h. We obtained half-hourly samples of subjective sleepiness (from 7 h before HBT) and saliva (from 6 h before HBT) until 1 h after HBT, plus a final evening sample taken 1 h and 20 min after HBT. 2 morning samples were taken 5 and 35 min after wake-up. PVT and VCS were administered on arrival as baseline, then 5 times during the protocol at 90-min intervals and the last time 1 h after HBT. Starting 7 h before HBT, skin temperatures were continuously monitored at 60-s intervals. PLR measurements, polysomnography (including KDTs), skin temperature after bedtime and sleep-related questionnaires taken in the morning were not included in this study. <u>Abbreviations:</u> AEE = afternoon to early evening, Exp. = Experimental sessions, HBT = Habitual bedtime (for home sleep), KDT = Karolinska Drowsiness Test, KSS = Karolinska Sleepiness Scale, mEDI = melanopic Equivalent Daylight Illuminance, PLR = Pupillary light response, PVT = Psychomotor Vigilance Task, VCS = Visual comfort and well-being Scales.

**Table 1:**
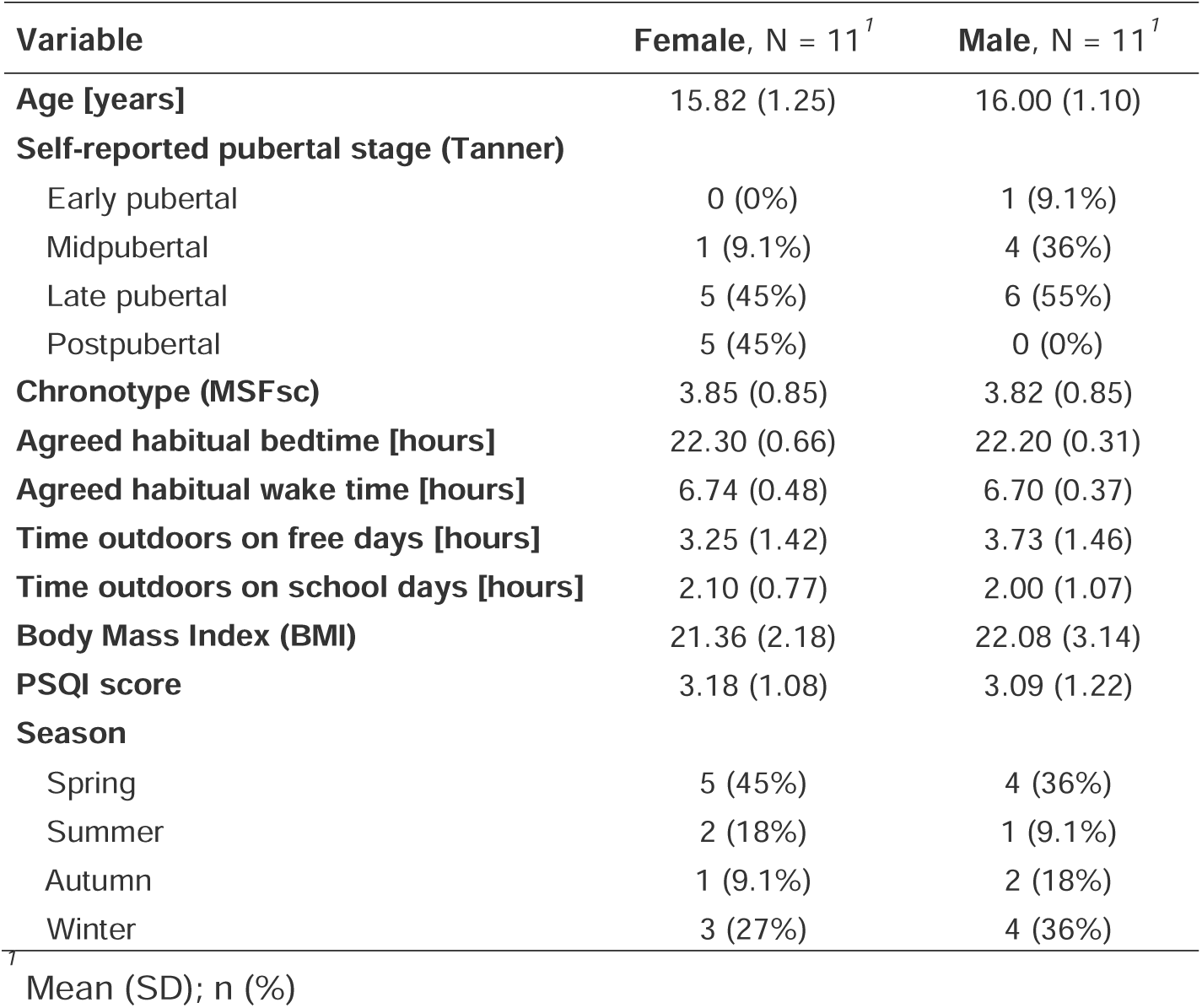
Participant characteristics stratified by sex. Summary of main participant characteristics, stratified by sex. Age, pubertal stage, chronotype (via the short MCTQ^147^), time spent outdoors and PSQI^161^ score were based on self-report measures collected at the screening and as part of the sleep diary. Habitual bedtimes and wake-up times were agreed during the in-person screening. Season (metrological) was based on participants’ dates of participation, while body mass index was based on in-person measurements of height and weight. Numerical variables are presented as “mean (SD)”, while categorical variables are presented as “n” (%). <u>Abbreviations</u>: AEE = afternoon to early evening, BMI = body mass index, MCTQ = Munich Chronotype Questionnaire, MSF_SC_ = Midpoint of sleep on free days corrected for “oversleep”, PSQI = Pittsburgh Sleep Quality Index.

| Variable | Female, N = 11<sup>†</sup> | Male, N = 11<sup>†</sup> |
| --- | --- | --- |
| **Age [years]** | 15.82 (1.25) | 16.00 (1.10) |
| **Self-reported pubertal stage (Tanner)** |  |  |
| Early pubertal | 0 (0%) | 1 (9.1%) |
| Midpubertal | 1 (9.1%) | 4 (36%) |
| Late pubertal | 5 (45%) | 6 (55%) |
| Postpubertal | 5 (45%) | 0 (0%) |
| **Chronotype (MSFsc)** | 3.85 (0.85) | 3.82 (0.85) |
| **Agreed habitual bedtime [hours]** | 22.30 (0.66) | 22.20 (0.31) |
| **Agreed habitual wake time [hours]** | 6.74 (0.48) | 6.70 (0.37) |
| **Time outdoors on free days [hours]** | 3.25 (1.42) | 3.73 (1.46) |
| **Time outdoors on school days [hours]** | 2.10 (0.77) | 2.00 (1.07) |
| **Body Mass Index (BMI)** | 21.36 (2.18) | 22.08 (3.14) |
| **PSQI score** | 3.18 (1.08) | 3.09 (1.22) |
| **Season** |  |  |
| Spring | 5 (45%) | 4 (36%) |
| Summer | 2 (18%) | 1 (9.1%) |
| Autumn | 1 (9.1%) | 2 (18%) |
| Winter | 3 (27%) | 4 (36%) |

<sup>†</sup> Mean (SD); n (%)

**Figure 2.**
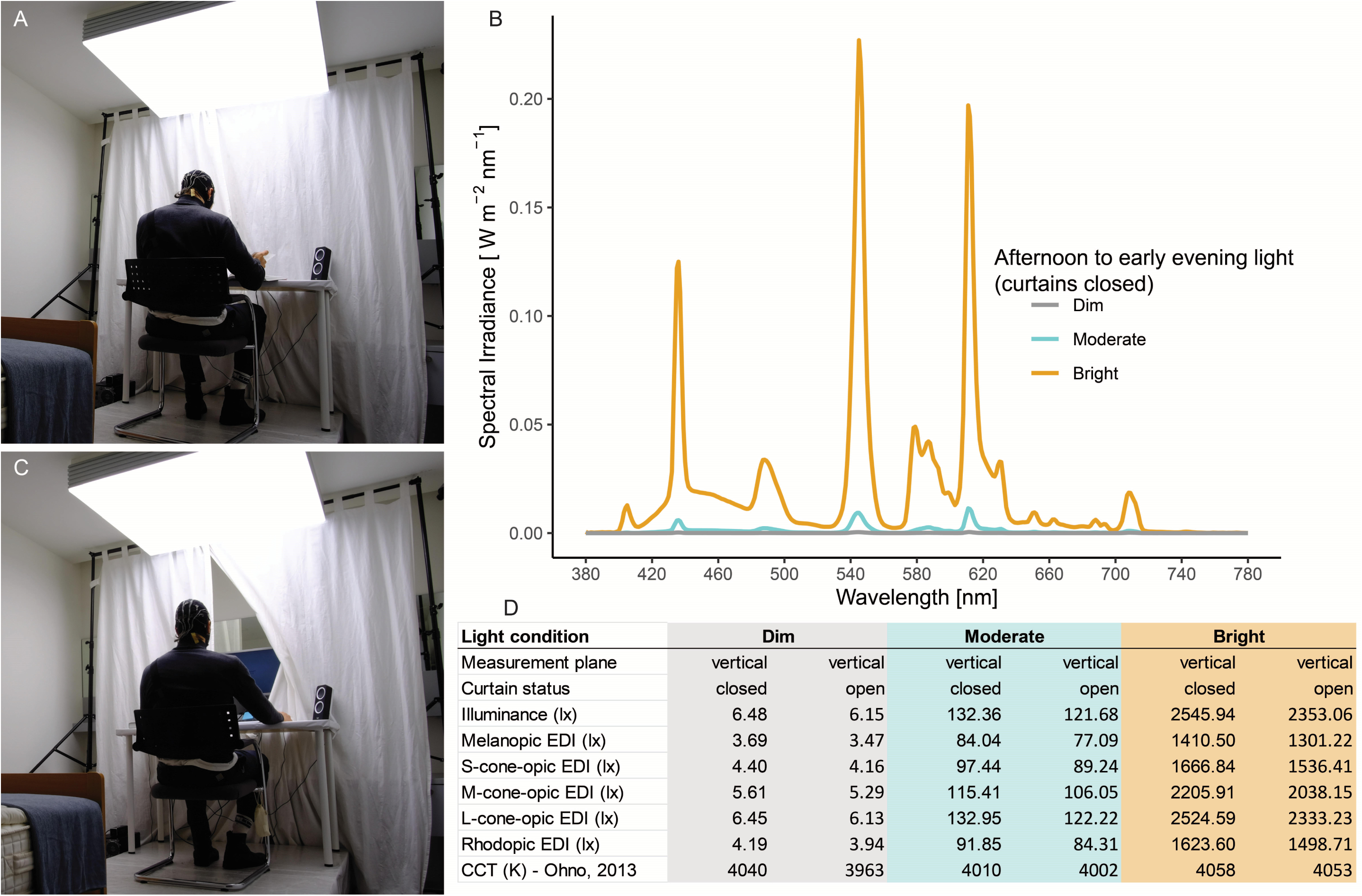
Laboratory setup and stimulus specification. The individual depicted is author R.L. **A,** Laboratory setup with the fluorescent ceiling lamp panel during task-free periods (curtains closed). **B**, Spectral irradiance distribution of the 3 AEE light exposure interventions during task-free periods (curtains closed). The vertical light measurements were taken with a spectroradiometer placed at the eye level of the seated participants (115 cm from the floor, 85 cm from the overhead light source, 80 cm from the white curtain, Spectraval 1501, JETI Technische Instrumente GmbH, Jena, Germany, last calibration: 07.03.2023). **C**, Laboratory setup during task performace using an LCD screen with black background and red lettering (curtains open). **D**, Tabulated stimulus specifications per light intervention during the experiment, derived from vertical spectral irradiance measurements on the participants’ eye level with curtains closed and open, including alpha-opic Equivalent Daylight Illuminances (alpha-opic EDIs) and Correlated Colour Temperature (CCT).

**Figure 3.**
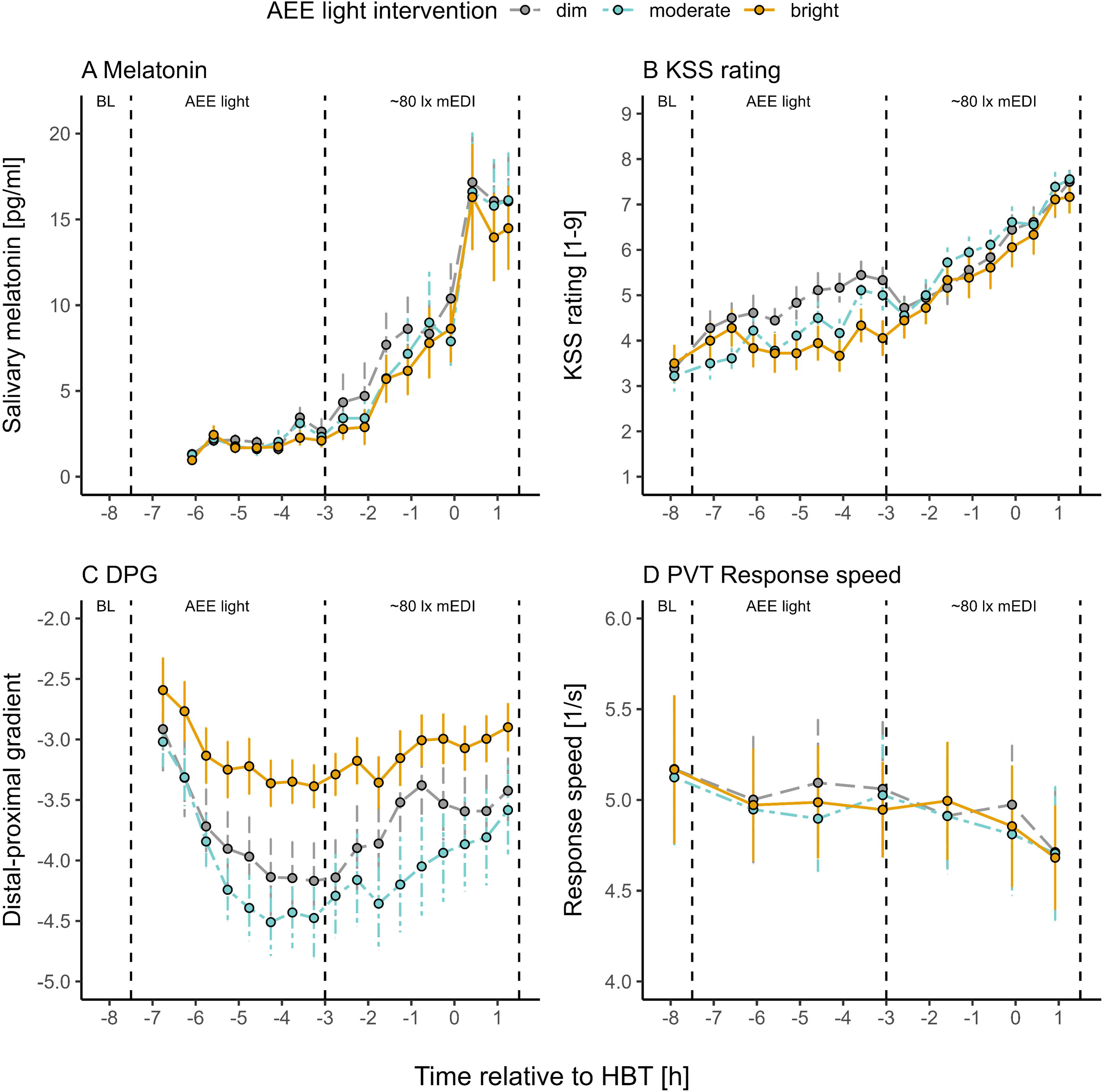
Time course of main outcomes across the protocol per light intervention condition relative to habitual bedtime. **A**, Salivary melatonin concentrations throughout the experimental evening based on data from 20 participants (n=2 excluded). HBT (0 on the x-axis) on average corresponded to 22:15 (SD=31 min). Coloured dots show the mean per condition (AEE light intervention), with error bars indicating the standard error. 2 single missing melatonin samples were estimated with linear interpolation (1 sample no. 9 and 13 after the dim and moderate light intervention, respectively). **B**, Subjective sleepiness ratings throughout the experiment based on data from 18 participants (4 excluded). Coloured dots show the mean per condition, with error bars indicating the standard error. **C**, Distal-proximal skin temperature gradient (DPG) throughout the experiment based on data from 21 participants (1 excluded). Coloured dots show the mean value in 30-min bins per condition, with error bars indicating the standard error. The first mean time bin (transition sample not used in LMM) is based on 19 single datapoints during the moderate light intervention,18 single datapoints for the “bright” light intervention and 17 single datapoints for dim light. **D**, PVT response speed (mean 1/RT) throughout the experiment based on data from 22 participants (0 excluded). Coloured dots show the mean value per condition, with error bars indicating the 95% confidence interval. Values at baseline (BL), not included in the LMM analysis, are based on 18 participants. <u>Abbreviations:</u> AEE = afternoon to early evening, BL = Baseline before light intervention, DPG = Distal-proximal skin temperature gradient, mEDI = melanopic Equivalent Daylight Illuminance, HBT = Habitual bedtime (for home sleep), KSS = Karolinska Sleepiness Scale, PVT = Psychomotor Vigilance Task, RT = Reaction Time.

**Figure 4.**
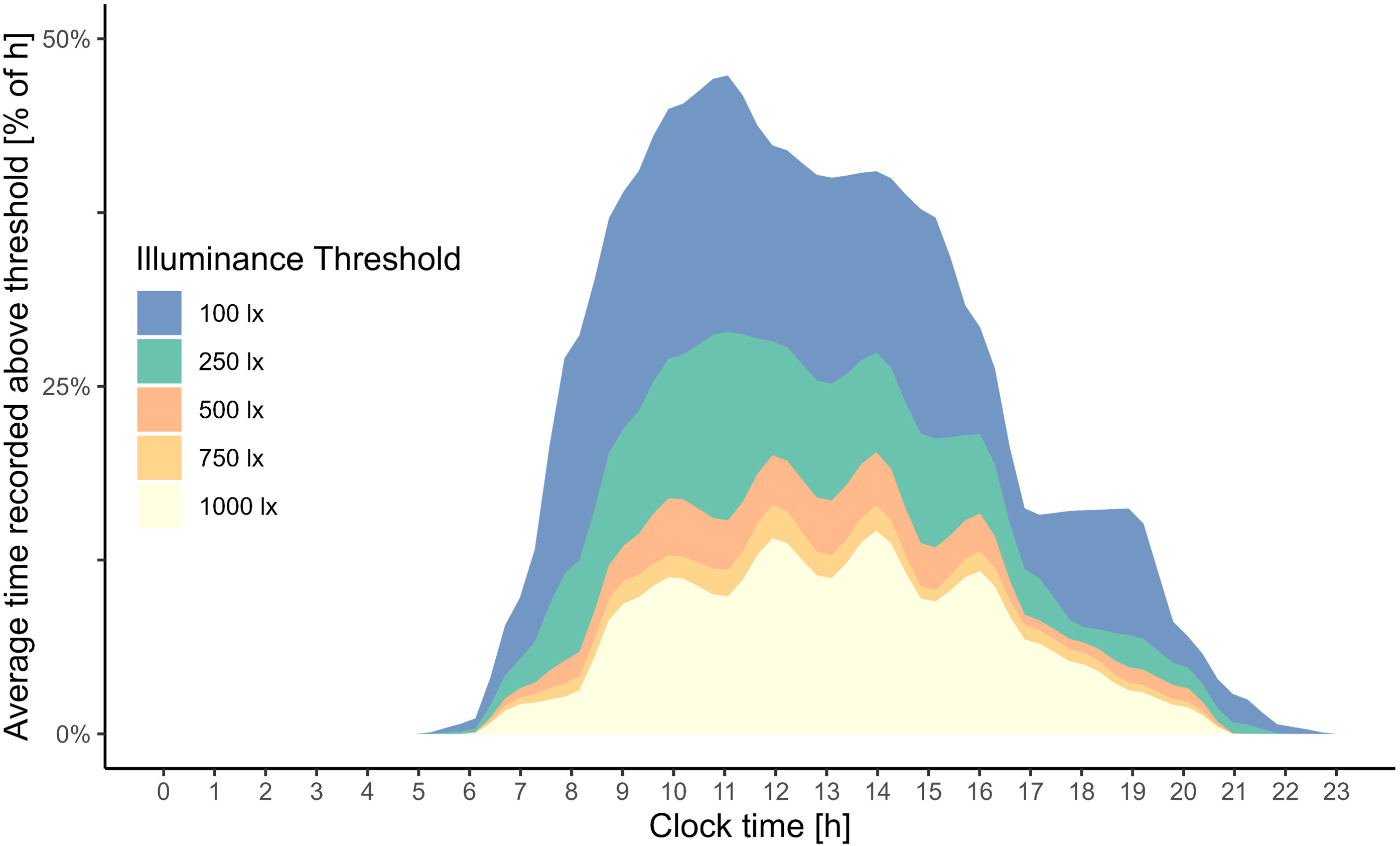
Average light exposure history on the day before experimental sessions. Distribution of light exposure history throughout the day before the experiment, averaged across all participants (n=22). The y-axis shows the wrist-recorded time above threshold (TAT) in % as a function of the clock time in hours on the x-axis. The different colours code the different wrist-recorded illuminance thresholds. For thresholds ≥500 lx, several midday peaks of increased TAT are visible, likely representing school breaks with opportunity for daylight exposure.

### Salivary melatonin

For our primary outcome, melatonin AUC during the later evening light exposure, n=20 participants were included in the linear mixed model (LMM) analysis. Figure 5 A shows the median and mean melatonin AUC values as a function of the preceding light intervention, together with the interquartile range (IQR) and individual data points, while Figure 3 A illustrates the average melatonin sample time course per light intervention relative to HBT. A full summary of the melatonin LMM estimates is given in Table 2 (sparse models in Suppl. Table S2).

**Figure 5.**
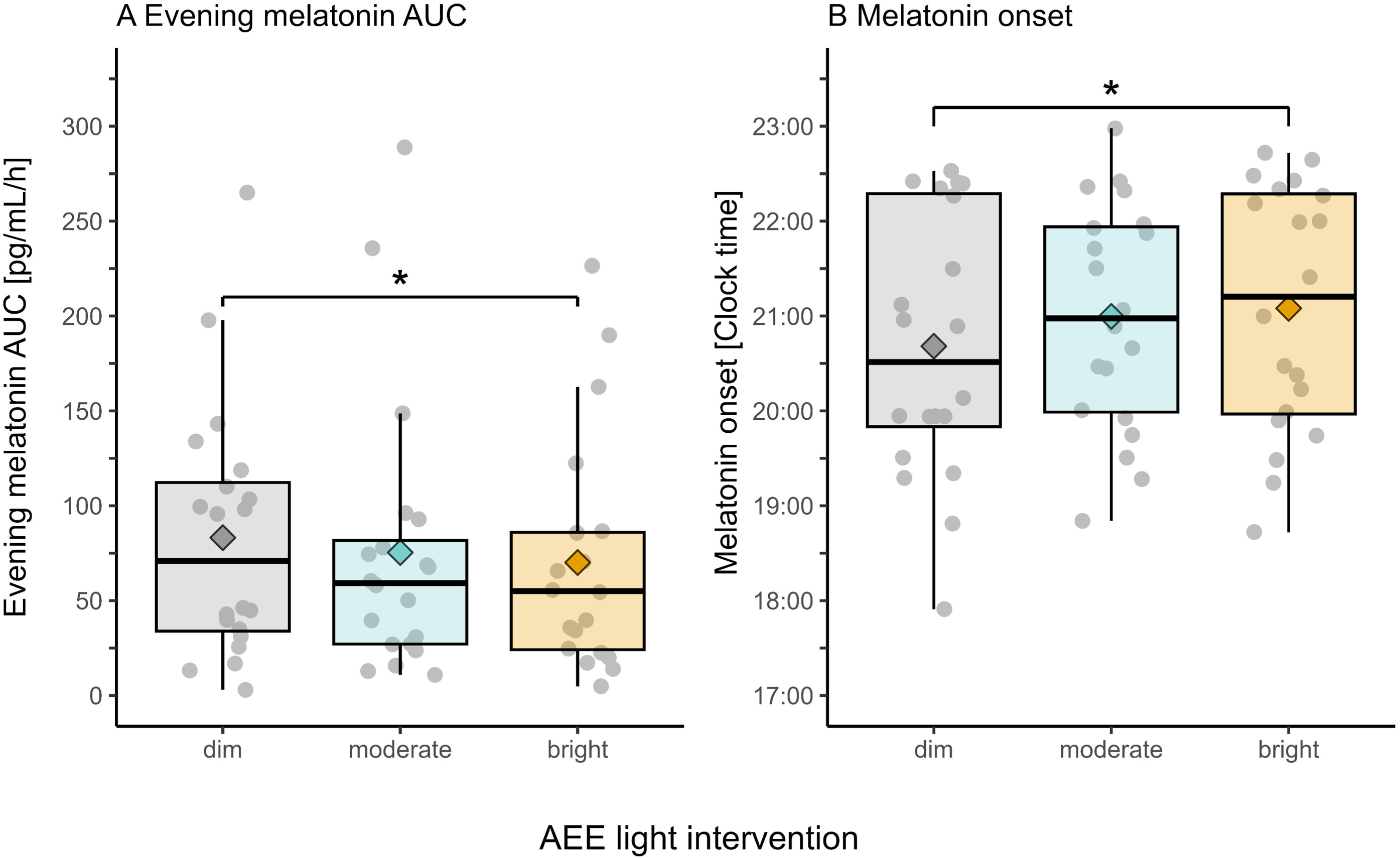
Salivary melatonin levels and onsets per light intervention condition. **A,** Evening melatonin AUC in pg/ml/h, across the 3 AEE light intervention conditions, based on data from 20 participants (n=2 excluded). Single missing evening melatonin samples (n=2) were estimated by linear interpolation. Coloured diamonds represent the mean melatonin AUC per condition. Coloured box plots show the interquartile range (IQR, box borders), the median (central horizontal line) and the range of minimum and maximum values within 1.5 times the IQR (whiskers). The grey dots mark the melatonin AUC of the individual participants in each condition. The significant difference (p<0.05) in melatonin AUC between the bright light intervention and dim light in the linear mixed model analysis is indicated by the black asterisk. **B,** Melatonin onset given in clock time, across the 3 AEE light intervention conditions, based on data from 20 participants (n=2 excluded). Single missing evening melatonin samples (n=2) were estimated by linear interpolation. Coloured diamonds represent the mean melatonin onset per condition. Coloured box plots show the interquartile range (IQR, box borders), the median (central horizontal line) and the range of minimum and maximum values within 1.5 times the IQR (whiskers). The grey dots mark the melatonin onset of the individual participants in each condition. The significant difference (p<0.05) in melatonin onset between the bright light intervention and dim light in the linear mixed model analysis is indicated by the black asterisk. <u>Abbreviations:</u> AEE = afternoon to early evening, AUC = Area under the curve, IQR = Interquartile range, MO = Melatonin onset.

**Table 2:** LMM analysis for melatonin AUC and melatonin onset (covariate model). Summary report of linear mixed model (LMM) analyses for melatonin AUC (primary outcome) and melatonin onset (post-hoc analysis), applying the model with theoretically relevant covariates (covariate model). Estimates of the AEE light intervention conditions are given relative to the dim light condition. Estimates of pubertal stage are given relative to “post pubertal stage”. P-Values of (p<0.05) were considered as significant (in bold). The covariates marked with an asterisk (*) were used as passive stratification factors to control for additional variance. Accordingly, these LMM estimates were not interpreted to avoid a “table 2 fallacy”^162^. <u>Abbreviations:</u> AEE = afternoon to early evening, AIC = Akaike Information Criterion, AUC = Area under the curve, CI = Confidence Interval (95%), df = degrees of freedom, ICC = Intraclass Correlation Coefficient, MSF_SC_ = Midpoint of sleep on free days corrected for oversleep, N_subjects_ = Number of subjects in the analysis, σ² = Variance of the residuals of the model, τ_00 subjects_ = Variance in the random intercepts at the subject level, std. Error = standard error of the estimates, TAT 1000 lx = Time above threshold (>1000 lx illuminance, wrist-recorded).

| Predictors | Estimates |  | CI (95%) |  | Std. Error |  | df |  | t value |  | p |  |
| --- | --- | --- | --- | --- | --- | --- | --- | --- | --- | --- | --- | --- |
|  | Melatonin AUC | Melatonin onset time | Melatonin AUC | Melatonin onset time | Melatonin AUC | Melatonin onset time | Melatonin AUC | Melatonin onset time | Melatonin AUC | Melatonin onset time | Melatonin AUC | Melatonin onset time |
| Intercept | 179.05 | 18.45 | 21.45, 336.88 | 15.57, 21.33 | 88.92 | 1.63 | 15 | 15 | - | - | - | - |
| Condition [moderate] | -7.37 | 0.30 | -16.24, 1.43 | -0.01, 0.61 | 4.56 | 0.16 | 37 | 37 | -1.62 | 1.88 | 0.114 | 0.068 |
| Condition [bright] | -11.29 | 0.35 | -20.34, -2.49 | 0.04, 0.67 | 4.60 | 0.16 | 37 | 37 | -2.45 | 2.17 | **0.019** | **0.037** |
| Bright light history [TAT 1000 lx] | 5.65 | -0.16 | 1.19, 9.37 | -0.28, -0.02 | 2.07 | 0.07 | 43 | 52 | 2.73 | -2.41 | **0.009** | **0.020** |
| *Chronotype [MSFsc] | -30.62 | 0.70 | -66.81, 5.56 | -0.04, 1.35 | 20.41 | 0.37 | 14 | 14 | -1.50 | 1.88 | 0.155 | 0.081 |
| *Pubertal stage [late pubertal] | 38.75 | -0.35 | -29.26, 106.71 | -1.58, 0.88 | 38.34 | 0.70 | 15 | 14 | 1.01 | -0.50 | 0.329 | 0.624 |
| *Pubertal Stage [midpubertal] | -21.37 | 0.06 | -98.36, 55.61 | -1.34, 1.45 | 43.41 | 0.79 | 14 | 14 | -0.49 | 0.07 | 0.630 | 0.945 |
| *Pubertal Stage [early pubertal] | 20.80 | -1.07 | -115.71, 157.17 | -3.55, 1.41 | 76.94 | 1.41 | 15 | 14 | 0.27 | -0.76 | 0.791 | 0.459 |
| **Random effects & Model fit criteria** |  |  | **$\sigma^2$** | **T<sub>00</sub> subjects** | **ICC** | **N<sub>subjects</sub>** | **Observations** | **Marginal R<sup>2</sup> / Conditional R<sup>2</sup>** |  | **AIC** |  |  |
| Evening melatonin AUC |  |  | 207.54 | 4623.46 | 0.96 | 20 | 60 | 0.221 / 0.967 |  | 530.364 |  |  |
| Melatonin onset time |  |  | 0.26 | 1.47 | 0.85 | 20 | 60 | 0.244 / 0.888 |  | 162.679 |  |  |

There was no significant effect on evening melatonin AUC following the moderate AEE light intervention compared to following dim light (β=-7.37, *p*=0.114). However, contrary to our hypothesis, there was a significant negative effect following the bright light intervention compared to following dim light (β=-11.29, *p*=0.019), i.e. previous bright light until 3 hours before HBT was associated with lower later evening melatonin levels (−11.29 pg/ml/h) compared to previous dim light.

The covariate “bright light history” (time above 1000 lx threshold, TAT 1000 lx) was a significant predictor in the model (β=5.65, *p*=0.009), indicating that a prolonged exposure to bright light (>1000 lx) from the previous day to laboratory entry was associated with an increase in evening melatonin AUC, i.e. 5.65 pg/ml/h per hour spent in above 1000 lx illuminance.

Following these findings, we performed an additional post-hoc analysis of the melatonin onset times (MO), which were determined with the hockey-stick method^82^, using the same LMM model approach without pre-specified hypotheses. A visualisation of the median and mean melatonin onset with individual data points and IQR, expressed in decimal hours (clock time), is shown in Figure 5 B. There was no significant effect on melatonin onset after the moderate AEE light intervention compared to after dim light (β=0.30, *p*=0.068). However, there was a significant positive effect after the bright light intervention compared to after dim light (β=0.35, *p*=0.037). The bright light intervention between 7.5 and 3 hours before HBT was associated with a 21 min later melatonin onset on that evening compared to the previous dim light (summary of the LMM estimates, see Table 2).

The covariate “bright light history” was a significant predictor in the model (β=-0.16, *p*=0.020), indicating that a prolonged exposure to bright light (>1000 lx) from the previous day to laboratory entry (∼32 h timespan) was associated with an earlier melatonin onset on the experimental evening, i.e. ∼10 min (0.16 h) earlier for each hour recorded in above 1000 lx illuminance. These results for melatonin onset are in close alignment with those observed for melatonin AUC.

For completeness, the 2 melatonin values from the next morning were analysed with the same LMM structure and are summarised in the Supplementary Information (see Supplementary Table S3).

### Subjective sleepiness

For subjective sleepiness during late evening light exposure, no significant effect was present following the moderate (β=0.14, *p*=0.265) or bright light intervention (β=-0.13, *p*=0.320) compared to following dim light. The factor “time” (median centred) showed a significant positive effect, with later time (1-hour increments) being associated with higher reported sleepiness (β=0.73, *p*<0.001). Neither the interaction “moderate light intervention * time” (β=0.01, p=0.958) nor the interaction “bright light intervention * time” (β=-0.02, p=0.830) showed a significant effect.

Bright light history was a significant covariate in the model (β=0.24, *p*<0.001), indicating that a prolonged exposure to bright light (>1000 lx) from the previous day to laboratory entry was associated with an increase in reported evening sleepiness (0.24 KSS units for every 60 minutes recorded in above 1000 lx illuminance).

As a post-hoc analysis, we further tested the acute light effects on sleepiness during the AEE light intervention, applying the same LMM structure. Significant negative effects were present both during the moderate (β=-0.63, *p*<0.001) and bright intervention (β=-0.92, *p*<0.001) compared to during the dim light. Factor “time” showed a significant positive effect, with later time (1-hour increments) being associated with a higher sleepiness level (β=0.29, *p*<0.001). The “moderate light intervention * time” interaction was not significant (β=0.08, *p*=0.477) compared to the interaction “dim light * time”. However, the interaction “bright light intervention * time” yielded a significant negative effect (β=-0.29, *p*=0.008), showing that subjective sleepiness increased less over time during the bright light intervention than during dim light (cf. Figure 3 B).

There was no significant effect of the bright light history covariate on sleepiness during the AEE light condition (β=-0.04, *p*=0.483).

A full summary of the subjective sleepiness LMM estimates is given in Suppl. Table S4 (sparse models in Suppl. Table S5).

### Vigilant attention/response speed

For PVT response speed during the later evening light exposure, there was no significant effect following the moderate AEE light intervention (β=-0.06, *p*=0.290) or the bright light intervention (β=-0.04, *p*=0.461) compared to after dim light. Factor “time” (median centred) did not show a significant effect (β=-0.07, *p*=0.070). Neither the interaction “moderate light intervention * time” (β=-0.01, p=0.846) nor the interaction “bright light intervention * time” (β=-0.05, p=0.341) showed a significant effect compared to the interaction “dim light * time”.

There was no significant effect of the bright light history covariate on response speed during the later evening light exposure (β=-0.04, *p*=0.126).

As a post-hoc analysis, we further tested the acute light effects on PVT response speed during the AEE light interventions, applying the same model structure. No significant effect was detectable during the moderate light intervention (β=-0.10, *p*=0.163) or bright light intervention (β=-0.09, *p*=0.176) compared to during dim light. No significant effect for the factor “time” (median centred) was detected (β=0.02, *p*=0.622). Neither the interaction “moderate light intervention * time” (β=0.01, p=0.907) nor the interaction “bright light intervention * time” (β=-0.05, p=0.622) showed a significant effect compared to the interaction “dim light * time”.

There was no significant effect of the bright light history covariate on response speed during the AEE light administration (β=-0.04, *p*=0.201).

A full summary of the LMM estimates for PVT mean response speed is shown in Suppl. Table S6 (sparse models in Suppl. Table S7). Time courses of PVT measures other than mean response speed (see Figure 3D) are shown in Suppl. Fig. S1.

### Skin temperature (DPG)

Contrary to our hypothesis, there was a significant negative effect on later evening DPG following the moderate light intervention, showing a lower DPG compared to after dim light (β=-0.41, *p*<0.001). In contrast to this finding, there was a significant positive effect after the bright light intervention, associated with a higher DPG compared to after dim light (β=0.49, *p*<0.001). Factor “time” (median centred) did not yield a significant effect (β=0.07, *p*=0.388). Neither the interaction “moderate light intervention * time” (β=0.17, *p*=0.136) nor the interaction “bright light intervention * time” (β=0.05, *p*=0.665) showed a significant effect compared to the interaction “dim light * time”.

There was no significant effect of the bright light history covariate on later evening DPG (β=0.00, *p*=0.994).

As a post-hoc analysis, we further tested the acute light effects on DPG *during* the AEE light intervention, applying the same LMM structure. A significant negative effect was observed during the moderate AEE light intervention compared to dim light (β=-0.27, *p*=0.008). Contrary to those results we found a significant positive effect during the bright light intervention, showing a higher DPG compared to during dim light (β=0.69, *p*<0.001). Factor “time” (median centred) showed a significant negative effect, with later time (1-hour increments) being associated with a lower DPG during the light interventions (β=-0.26, *p*<0.001). Neither the interaction “moderate light intervention * time” (β=-0.09, *p*=0.357), nor the interaction “bright light intervention * time” was significant (β=0.09, *p*=0.374) compared to the interaction “dim light * time”.

There was no significant effect of the bright light history covariate on DPG during the light interventions (β=-0.02, *p*=0.652).

A full summary of the LMM estimates for DPG is given in Suppl. Table S8 (sparse models in Suppl. Table S9). More detailed skin temperature data showing the distal-proximal temperature bands and the distal temperatures in hands and feet are shown in Suppl. Figure S2, panels A, B and C, while results for room temperature are described in the Supplementary Information and presented in Suppl. Fig. S2 D, E and F. The visual comfort and well-being ratings are shown in Suppl. Fig. S3 and S4).

## Discussion

In the present study, we set out to investigate whether increasing after-school light exposure could be a potential strategy for increasing melatonin levels and mitigating the alerting effects of artificial light around bedtime in adolescents.

Contrary to our hypothesis, we found evidence that later evening melatonin levels were reduced when preceded by the bright light intervention (7.5 and 3 hours before HBT). We found no effect on evening sleepiness or PVT response speed, but, contrary to our hypothesis, a lower DPG after the moderate light intervention. The bright light intervention resulted in a higher DPG (in line with our hypothesis), but this was confounded by an increased room temperature. Interestingly, the wrist-recorded bright light history of the previous day to the scheduled laboratory session was associated with an earlier melatonin onset, and increased evening melatonin and sleepiness levels.

The most striking finding of this study is that later evening melatonin levels were reduced when preceded by the bright light intervention (7.5 and 3 hours before HBT). These results contrast with previous studies that reported increased melatonin levels and decreased alertness following bright light adaptation in adults.

One study based on a similar experimental design in young adult women (n=12) found a significantly greater increase (pre-post difference) in melatonin during late evening light exposure (750 lx illuminance, 22:30-23:30) when preceded by bright (1200 lx, 18:30-21:00) than when preceded by dim light. However, when comparing melatonin AUC, DPG or subjective sleepiness, the authors only report a “trend” and no statistically significant differences for these conditions^79^. The discrepancies with our findings may be due to the different samples, light characteristics, their inclusion of dim light adaptation, or the use of the same clock schedule for all participants as opposed to the individualised times related to HBT in the present study.

Other studies in adults that have found decreased melatonin suppression after light adaptation have used a markedly different timing of prior light exposure. Two reports showed reduced nighttime salivary melatonin suppression after morning exposure to bright light (900 or 2700 lx illuminance)^74^ or blue light (79 lx)^76^ compared to dim (<10 lx) or white (100 lx) light, respectively. Another study found reduced nocturnal plasma melatonin suppression under full-field monochromatic blue light exposure at 460 nm (3.1 and 7 μW/cm²) when preceded by dim light (18 lx, midnight to 2 am) compared to dark adaptation^67^. Additionally, several studies in adults used earlier and longer-term light exposure protocols over multiple days before testing for differences in melatonin levels^33,64–66,68,71,73,83^, compared with the 4.5 h AEE light intervention in the present study. They found that a day to a week of bright light exposure led to increased nighttime urinary melatonin secretion^68,83^, blunted melatonin suppression^33^, increased melatonin amplitude^66^ and AUC^64,66^, as well as decreased nocturnal CBT^83^. Others found a decrease in melatonin suppression after days of moderate light exposure^65,71^, and increased subjective sleepiness and PVT lapses^71^.

The discrepancies between these studies and the unexpected later evening melatonin reduction in the present trial may be explained by the timing of our AEE light intervention. It is plausible, that the AEE bright light until 3 hours before HBT had an acute phase-delaying effect on melatonin release, overriding any potential increase of melatonin secretion amplitude or adaptation of circadian photosensitivity. Consistent with that notion, our post-hoc analysis showed that the preceding bright light intervention was associated with a 21-min later melatonin onset compared to the preceding dim light condition. The moderate light intervention, curiously, did not yield significant results, but indicated the same effect direction on melatonin as the bright light intervention.

Interestingly, Crowley and Eastman^25^ found that exposing adolescents to 2 h of bright light in the late afternoon for 3 consecutive days advanced their dim light melatonin onset by about 25 min. In our study, the late afternoon light exposure occurred around similar clock times on average, but later relative to our participants’ habitual sleep schedule. These differences in relative timing and their repetition across 3 days may explain why, contrary to their findings, we found an acute reduction in melatonin and some indications for an acute phase delay after the AEE bright light intervention. Although the largest proportion of the phase-delaying and melatonin-suppressing effects of light appear to occur within the first hour of exposure^84^, evidence from young adults^29,85^ suggests that bright light in the late afternoon may already cause phase delays, which is consistent with our findings. Curiously, our data showed that the adolescents’ wrist-recorded bright light history from the morning of the previous day to the scheduled laboratory session (∼32 h period) was positively associated with increased evening melatonin levels, an earlier melatonin onset, and increased subjective sleepiness on the experimental evenings, but not during the afternoon. Thus, in contrast to the very recent prior light history manipulated in the experiment, the bright light history from the day before may have been related to phase advances, paralleled by higher melatonin levels and evening sleepiness. It is also possible that the increased exposure to daylight, which acts as a strong ‘zeitgeber’, synchronised individuals more closely with the natural light-dark cycle and increased the amplitude of their circadian oscillator, resulting in greater evening melatonin production and sleepiness ^60,66^. These findings may also suggest that the adaptation of circadian photosensitivity in adolescents with regards to melatonin suppression and alertness may not manifest within a couple of hours but rather a couple of days.

Our wrist-recorded bright light history results are consistent with those observed in other studies using ambulatory light history data. This includes a report indicating an association between increased wrist-recorded light exposure in the previous 24 h and earlier sleep onset in adolescents^81^, a study in elderly people showing that time spent in daylight above 1000 lx illuminance within the previous 48 h was positively related to nocturnal urinary melatonin excretion^86^, and a study in adult indoor and outdoor workers showing that higher average 24-h wrist-recorded light exposure was associated with a lower percentage of melatonin suppression on experimental evenings^78^.

We did not observe any effect of the AEE light interventions on later evening sleepiness or PVT response speed. As expected, evening sleepiness increased over time, most likely due to decreasing circadian wake promotion and increasing sleep pressure^87–89^. Even though the adolescents remained awake until 90 min after their HBT and reported high levels of sleepiness, we found no significant effect of time on evening vigilance response speed (mean 1/RT), a measure shown to be sensitive to increasing sleep pressure^90^. Similarly, Agostini et al.^20^ only found an effect of time on morning PVT lapses compared to the rest of the day in late- and post-pubertal adolescents. These findings seem plausible in view of previous reports, suggesting that the accumulation of sleep pressure slows down during puberty such that pubertal and post pubertal adolescents have a higher tolerance for staying awake after their HBT^16–18^.

Our findings for evening skin temperature were mixed. We found a reduced DPG after the moderate light compared to dim light, contrary to our hypothesis. These results fall in line with Takasu et al.^66^ and may suggest that the moderate light intervention sustained sympathetic activation until later in the evening rather than reducing it. In contrast, we found an elevated DPG after the bright light intervention but this effect may have been influenced by a higher ambient room temperature, which has been shown to enhance heat loss through distal regions^91,92^. This raises uncertainty about whether the higher DPG was due to the light itself or the room temperature and warrants caution in interpreting this result.

In our study, we observed acute alerting responses of moderate and bright AEE light. During the bright light intervention, subjective sleepiness increased less over time compared with dim light, and both the moderate and bright light intervention acutely produced lower subjective sleepiness but did not affect PVT response speed. The light interventions also acutely affected DPG, with lower DPG during the moderate compared to dim light and higher DPG during the bright light intervention.

Several studies, meta-analyses, and reviews have examined the daytime alerting effects of light^93–97^. A recent mini-review^98^ summarizes these findings and concludes that different intensities and qualities of light can increase both objective and subjective alertness, increase activity levels, modulate brain responses, and potentially improve cognitive performance during the day^98^. Siraji et al^93^ reported in their review that of the studies investigating bright light exposure, particularly in the afternoon, ∼57% (8 out of 14) showed a reduction of subjective sleepiness and only ∼17% (1 out of 6) demonstrated a decrease in reaction times. Our study accords with the majority of these studies in showing that the AEE light interventions were effective in acutely inducing subjective alertness but not in terms of PVT reaction time. It remains unclear to what extent the reported sleepiness was psychologically influenced by the perception of brightness as participants also clearly perceived the light interventions as being of different brightness (see Suppl. Figure S3 A, B). In addition, previous research has indicated a close relationship between thermoregulation and sleepiness regulation^99–103^ where DPG reflects the body’s readiness for sleep through core cooling via peripheral vasodilation^101^. Consistent with this literature, we found reduced sleepiness together with lower DPG during the moderate light intervention. In contrast, the reduction in sleepiness during the bright light intervention was linked to an increased DPG, again, likely masked by the higher ambient room temperature^91,92^. In general, increasing light levels have been shown to decrease DPG^37,104^ and increase CBT^31,38,105^ via sympathetic activation^106^, although these effects may be time-dependent and more pronounced when the circadian timing system promotes sleep^37,104,105^. Other studies showed contradictory effects of bright light on DPG and CBT^91,107–110^, rendering the effects of daytime light exposure on thermoregulation inconclusive.

Consensus lighting recommendations for circadian health in adults suggest that light levels should be dimmed at least 3 h before bedtime^111^. Conversely, the present findings suggest that reducing light 3 h before HBT may not be early enough for healthy adolescents. Taken together, our data indicate that afternoon exposure to bright light up to 3 h before habitual bedtime has acute alerting and possibly persistent thermoregulatory effects and reduces melatonin levels in adolescents during subsequent light exposure later in the evening, possibly via a circadian phase delay. It may therefore be inappropriate for mitigating the non-visual effects of light around bedtime on the same evening. Notably, while behavioural interventions^112,113^ and morning light exposure^49–52^ have shown some potential, a growing body of evidence indicates that delaying school start times for middle and high school students effectively improves sleep and circadian health^114–120^, suggesting that policy changes are needed to address the widespread problem of inadequate sleep in adolescents^121,122^.

Despite the insights that can be drawn from this study, we acknowledge several limitations. First, the lack of an evening dim light condition prevented us from determining a dim light melatonin onset (DLMO) and calculating the relative melatonin suppression under later evening moderate light. In addition, although we estimated melatonin onset under moderate light conditions, we did not record salivary melatonin levels over two consecutive evenings, which prevented us from reliably investigating circadian phase shifts in response to the different light conditions. The visible difference in brightness between the lighting interventions made it impossible to blind participants, despite counterbalancing the order of application. However, this should not have affected objective measures such as our primary outcome melatonin, skin temperature or PVT responses. Furthermore, the bright light intervention was confounded by differences in room temperature, which specifically affected the skin temperature for this condition and limited the interpretation of these results. Finally, our results were based on a rather small sample of healthy adolescents (aged 14-17 years), which limited our ability to test for sex differences and other demographic variations. The sample size and cohort studied, although sufficient to detect significant effects in the present study, may limit the generalisability of our findings to a wider population.

This study also has several strengths. To our knowledge, it is the first controlled experiment to examine the effects of afternoon light exposure on alertness and melatonin levels during subsequent evening light exposure in adolescents. The study was conducted in a well-controlled laboratory setting that allowed precise manipulation and measurement of light exposure, while multimodal monitoring allowed inclusion of relevant covariates such as pre-experimental bright light history. In addition, precise measurement of salivary melatonin with high temporal resolution using a sensitive radioimmunoassay provided accurate estimates of our primary outcome. The within-subject design minimised the confounding effects of individual variability, thereby increasing the accuracy of the results. In addition, the comprehensive multi-stage screening process, which included various health assessments, ensured the inclusion of a healthy sex-balanced sample. Realistic evening and afternoon light levels and after-school hours were used to ensure transferability to real-world contexts while maintaining a high level of experimental control.

Our study highlights the complex effects of differently timed light and emphasises the importance of considering light exposure history when assessing a particular light stimulus. The present data raise the possibility that light history-driven adaptations of circadian photosensitivity may take longer than a few hours to manifest in adolescents. The physiological mechanisms underlying these adaptations are not well understood and there is no consensus on light exposure history in terms of ‘how long ago’ is meaningful^111,123^. Further research is needed to clarify how circadian photosensitivity adapts to different patterns of light exposure, including differences in timing, duration and intensity, and how these effects may differ in different populations, such as age groups. Building on current models for predicting non-visual effects of light^40,124–129^, we propose that future work can then aim to develop complex models that consider the integration of light exposure beyond instantaneous light measurements to more accurately reflect the complex nature of non-visual responses to light.

## Methods

### Ethics information

This study was carried out in the Centre for Chronobiology in Basel (Switzerland), between September 2022 and July 2023. The experimental protocol, screening questionnaires and consent form were approved by the Ethics Committee northwest/central Switzerland (2022-00432) and pre-registered as a clinical trial (https://clinicaltrials.gov/study/NCT05483296). The study execution strictly adhered to the Declaration of Helsinki, and all the recruited participants and their legal representatives were fully informed with study details and consented in a written form.

### Participants

A total of 82 students initially expressed interest in participating in the study, of whom a sex-balanced sample (n=22, 11 female) of healthy German-speaking teenagers (14-17 years) completed the study protocol (see demographics in Table 1). The selection criteria, screening instruments used, and corresponding cut-off points are shown in Suppl. Table S1, while the number of exclusions and withdrawals with their reasons are shown and described in Suppl. Fig. S5. Full participation in the study was compensated with 550 (CHF), or a partial amount in case of dropout. All questionnaires and instructions during the screening and study procedures were administered in German language. If an individual was excluded for non-adherence during the sleep-wake rhythm stabilisation period but had already completed one or more study visits, none of their data were used.

### Study protocol

The study procedure including study sampling and timing is illustrated in Figure 1. Before each scheduled experimental session, the ambulatory part of the protocol consisted of ≥5 days of sleep-wake rhythm stabilisation, during which participants were instructed to adhere to agreed target habitual bedtimes (HBT, lights off) and habitual wake times (HWT) within 1 h (±30 min, ≥8 h sleep opportunity window), and to abstain from alcohol, caffeine, and other drugs/medications. The target times were agreed-with the adolescent participants and their legal representatives to ensure compatibility with daily life, school, and their habitual sleep-wake rhythm, while requiring 8-10 h sleep opportunity, corresponding to the National Sleep Foundation’s recommendations at that age^130^, to prevent large sleep debt during the experiment. Adherence was monitored starting ≥5 days before each in-lab session, using wrist actimetry (ActTrust 2; Condor Instruments, Brazil) and self-report sleep diaries^131^. Non-compliance (HBT or HWT deviation of ≥1 h) in more than one night per study week led to exclusion from study participation (see Suppl. Table S1). During each of the 3 laboratory sessions, participants spent 18.5 h (including sleep) without external time cues such as watches, smartphones and daylight in the light-, temperature- and sound-controlled laboratory facilities of the Centre for Chronobiology, University Psychiatric Clinics in Basel, Switzerland. Baseline room temperature for the laboratories, corridor and toilets were 22°C with ventilation provided by an automated integrated air conditioning system in the facility. The laboratory sessions were individually scheduled to begin 8 h before HBT and last until 10.5 h after HBT. On entering the laboratory, participants were asked to take a urine-based multi-panel drug test (Nal von Minden; Moers, Germany) and a breathalyser-based alcohol test (ACE X, ACE Instruments; Freilassing, Germany) under bathroom lighting (see Suppl. Table S10), while their previous sleep-wake pattern and caffeine consumption were checked. An in-lab 8-h sleep opportunity window in darkness was scheduled between 1.5 and 9.5 h after the agreed-upon at-home bedtime (HBT). 4 approximately equicaloric meals were provided in 2-h intervals (first meal: 6 h before HBT) with similar food content across participants. In task-free periods, participants were allowed to read and study, play games, or engage in other activities that do not require self-luminous displays (e.g., listening to music and podcasts on provided speakers), but were instructed to remain seated and monitored to keep their eyes open with their sitting posture directed forward. Participants were asked to keep their activities consistent across all 3 experimental sessions. While awake, participants were instructed to take toilet breaks and 3-min walking activity in the corridor, scheduled in 1.5-h intervals.

After 8 h of sleep opportunity in darkness (1.5 to 9.5 h after HBT), participants were awakened in the morning and provided 2 more morning saliva samples, subjective ratings of sleepiness and completed a self-reported sleep quality questionnaire^132^ within a 30 min time window before showering, eating breakfast and leaving the laboratory.

### Light conditions

During the experiment, the participants received overhead room light exposure from a large (119 cm by 119 cm) fluorescent ceiling lamp panel (Philips Lighting, Eindhoven, The Netherlands), which comprised 2700 K fluorescent tubes (Master TL5 HO 54 W/827) and 17,000 K fluorescent tubes (Master TL5 HO Activiva Active 54W 1sl). These lamps have been successfully applied in previous studies with young adults in the Centre for Chronobiology^133^. In the dim light condition, we used a neutral density filter (211, 0.9 ND, LEE Filters Worldwide, Andover, UK) covering the light source to achieve homogenous dim light from the same lamp. The study room was furnished with a white curtain and a white tablecloth directly in front of the participant’s seat to ensure homogeneous light reflection. All walls had a uniform, highly reflective white paint finish, and the floor adjacent to the participant was covered with white fleece (see Fig. 2, Panel A, C). The experimental light conditions had an approximately equivalent spectral composition, with a colour temperature between 4000K and 4100K (see Fig. 2, Panel D). Light characteristics were measured vertically with a spectroradiometer placed at the eye level of the seated participants (115 cm from the floor, 85 cm from the overhead light source, 80 cm from the white curtain, Spectraval 1501, JETI Technische Instrumente GmbH, Jena, Germany, last calibration: 07.03.2023). Over the course of an experimental lighting condition, the light level varied by approximately 5-9%. Subjective sleepiness, visual comfort and well-being scales, as well as instructions for the auditory psychomotor vigilance test and the Karolinska drowsiness test, were displayed on a 22-inch LCD screen (Samsung SyncMaster 2243EW) with a black background and red lettering (red letters. luminance=3.20 Cd/m², melanopic EDL=1.87 Cd/m²) in order not to interfere with the lighting conditions. The screen was hidden behind a white curtain (see Fig. 2, Panel A), and only visible when the screen was in use (see Fig. 2, Panel C). Whenever participants left the study room after the experiment started (for bathroom breaks and pupil measurements), they were instructed to wear red-orange tinted filter goggles (see Suppl. Fig. S6), while all lights along their path were dimmed to avoid aberrant light exposure (illuminance≤6 lx, mEDI≤0.2 lx at eye level, see Suppl. Table S10).

#### AEE light intervention

The light exposure intervention started in the afternoon after conducting compliance checks, baseline sleepiness, vigilant attention, and pupil measurements (∼20 min incl. 3 min dark adaptation, properties shown Suppl. Table 11). Light exposure was consistent in timing and duration across all 3 experimental sessions of a participant, ranging from 7.5 h to 3 h before each participant’s HBT (∼3.75-h net light exposure over a 4.5-h period). Light levels differed by ∼1.3 log_10_-unit increments of melanopic equivalent daylight illuminance (mEDI) between the 3 experimental AEE light conditions (dim ∼4 lx; moderate ∼80 lx; bright ∼1400 lx, vertical plane with curtains closed, see Figure 2). The full tabulated spectral power distributions in CSV format for the conditions, displayed in Figure 2 b and in Suppl. Fig S7 in logarithmic scaling, are provided along with the datasets for analysis (see *Code and data availability*). Different light units derived from spectral irradiance and radiance values, measured at different angles, directions, and reflection scenarios, are given in Table 3 (irradiance), Suppl. Figure S8 (radiance), as well as Suppl. Table S12 (radiance). 45 min before the end of the light intervention, a second pupil measurement session was conducted. A summary report about the light conditions using the ENLIGHT checklist^134^ is provided in Suppl. Table S13.

**Table 3:** Detailed stimulus specification of the light interventions. Summary of the spectral irradiance derived values for the 3 AEE light interventions (between 7.5 and 3 h before HBT) at different angles and reflectance scenarios. Measurements were taken vertically facing forward, 45° angled downwards with an open book on the table, and vertically, with the sensor 90° angled to the left and right. The subsequent evening light condition (from 3 h before HBT to 1.5 h after HBT) had the same characteristics as the ‘moderate’ light intervention. Participants were instructed and monitored to remain seated and keep their eyes open with their sitting posture directed forward. The spectral irradiance measurements were taken from the observer’s point of view with a research-grade spectroradiometer (Spectraval 1501, JETI Technische Instrumente GmbH, Jena, Germany). The sensor was placed at the eye level of the seated participants (115 cm from the floor, 85 cm from the overhead light source, 80 cm from the white curtain in front of them). Values used for stimulus specification include illuminance, alpha-opic equivalent daylight illuminances (alpha-opic EDIs), alpha-opic irradiances (given in mW ⋅ m⁻²), CIE 1931 xy chromaticity coordinates, correlated colour temperature (CCT), and colour rendering index (CRI). These values were calculated from spectral irradiance measurements using the luox application^163^.

|  | Light intervention condition |  |  |  |  |  |  |  |  |  |  |  |
| --- | --- | --- | --- | --- | --- | --- | --- | --- | --- | --- | --- | --- |
|  | Dim |  |  |  | Moderate |  |  |  | Bright |  |  |  |
| Sensor orientation | forward | forward | 90°left | 90° right | forward | forward | 90°left | 90° right | forward | forward | 90°left | 90° right |
| Measurement plane | vertical | 45° down (book) | vertical | vertical | vertical | 45° down (book) | vertical | vertical | vertical | 45° down (book) | vertical | vertical |
| Illuminance (lx) | 6.48 | 3.86 | 3.53 | 3.86 | 132.36 | 83.12 | 73.90 | 77.11 | 2545.94 | 1488.91 | 1389.34 | 1413.00 |
| Melanopic EDI (lx) | 3.69 | 2.19 | 2.00 | 2.21 | 84.04 | 52.00 | 45.65 | 47.71 | 1410.50 | 813.98 | 753.19 | 765.66 |
| S-cone-opic EDI (lx) | 4.40 | 2.59 | 2.30 | 2.54 | 97.44 | 59.16 | 51.24 | 53.85 | 1666.84 | 945.18 | 873.76 | 889.11 |
| M-cone-opic EDI (lx) | 5.61 | 3.34 | 3.06 | 3.35 | 115.41 | 72.29 | 64.08 | 66.96 | 2205.91 | 1287.58 | 1200.69 | 1220.11 |
| L-cone-opic EDI (lx) | 6.45 | 3.84 | 3.51 | 3.83 | 132.95 | 83.46 | 74.20 | 77.38 | 2524.59 | 1475.82 | 1376.36 | 1400.34 |
| Rhodopic EDI (lx) | 4.19 | 2.49 | 2.28 | 2.50 | 91.85 | 57.10 | 50.30 | 52.58 | 1623.60 | 941.45 | 873.86 | 887.92 |
| Melanopic irradiance (mW · m<sup>-2</sup>) | 4.90 | 2.91 | 2.66 | 2.93 | 111.46 | 68.96 | 60.54 | 63.27 | 1870.62 | 1079.51 | 998.89 | 1015.42 |
| S-cone-opic irradiance (mW · m<sup>-2</sup>) | 3.60 | 2.11 | 1.88 | 2.08 | 79.64 | 48.35 | 41.88 | 44.01 | 1362.29 | 772.48 | 714.12 | 726.66 |
| M-cone-opic irradiance (mW · m<sup>-2</sup>) | 8.17 | 4.86 | 4.45 | 4.88 | 168.02 | 105.24 | 93.30 | 97.48 | 3211.43 | 1874.49 | 1748.00 | 1776.26 |
| L-cone-opic irradiance (mW · m<sup>-2</sup>) | 10.51 | 6.26 | 5.72 | 6.24 | 216.57 | 135.95 | 120.87 | 126.04 | 4112.33 | 2403.98 | 2241.96 | 2281.02 |
| Rhodopic irradiance (mW · m<sup>-2</sup>) | 6.07 | 3.61 | 3.30 | 3.63 | 133.16 | 82.78 | 72.92 | 76.23 | 2353.73 | 1364.83 | 1266.84 | 1287.22 |
| CIE 1964 x<sub>10</sub>y<sub>10</sub> chromaticity [x<sub>10</sub>] | 0.38 | 0.38 | 0.38 | 0.38 | 0.38 | 0.38 | 0.38 | 0.38 | 0.38 | 0.38 | 0.38 | 0.38 |
| CIE 1964 x<sub>10</sub>y<sub>10</sub> chromaticity [y<sub>10</sub>] | 0.35 | 0.36 | 0.36 | 0.36 | 0.35 | 0.35 | 0.35 | 0.35 | 0.36 | 0.36 | 0.36 | 0.36 |
| CCT (K) - Ohno, 2013 | 4040 | 4018 | 4022 | 4061 | 4010 | 3956 | 3904 | 3933 | 4058 | 4019 | 4014 | 4000 |
| Colour Rendering Index [Ra] | 80 | 80 | 80 | 80 | 87 | 87 | 87 | 87 | 80 | 79 | 79 | 79 |

#### Later evening light condition

The later evening light exposure ranged from 3 h before HBT to 1.5 h after HBT (∼4 h net light exposure over a 4.5-h period) and had the same spectrum and light level as the “moderate” AEE light condition (∼130 lx illuminance, ∼80 lx mEDI, ∼4010K). This light level is within the range of common evening domestic lighting conditions^58^ and has been shown to reduce melatonin levels in adolescents^23^.

### Salivary melatonin

Saliva samples (>1ml) were collected every 30 min from 6 h before HBT until 1 h after HBT, plus a final evening sample taken 1 h and 20 min after HBT (10 min before the experimental bedtime). Two samples were collected in the morning, 5 and 35 min after the experimental wake-up time. In total, 18 samples were collected per session, 7 during the AEE intervention, 9 during the later evening light condition that followed the AEE light intervention, and 2 in the morning. Melatonin was measured using a direct double antibody radioimmunoassay (RIA, RK-DSM261) with a limit of quantification of 0.9 pg/mL, an analytical sensitivity of 0.2 pg/mL, an intra-assay precision of 7.9% and an inter-assay precision of 9.8%. Measurements were performed by an external service laboratory (NovoLytiX GmbH, Witterswil, Switzerland). The area under the curve (AUC) of salivary melatonin was calculated using the trapezoidal method (AUC function from the “DescTools” package^135^) on samples taken during the evening light exposure, starting 3 h before HBT. Additionally, we visualised the time course of melatonin (t1, t2, …tn). As a post-hoc analysis, the melatonin onset (MO) was determined based on the 16 samples from the afternoon to experiment bedtime and calculated with a piecewise linear-parabolic function using the hockey stick method (v2.5)^82^ with a default area of interest upper border of 5 pg/ml. When necessary, default area of interest upper border was adapted based on an individual’s baseline melatonin values, following the procedure of Blume et al^136^. R.L. and C.C. independently performed the MO analyses while blinded to the experimental conditions. Divergences were then resolved in a final discussion with F.F.

### Subjective sleepiness

Subjective sleepiness was assessed every 30 min using the German version of the single-item 9-point Karolinska Sleepiness Scale (KSS) sleepiness^137^ plus a baseline value immediately after arrival.

### Vigilant attention

Reaction time performance of vigilant attention was measured using a modified auditory version of the Psychomotor Vigilance Test (PVT)^138^, which was terminated with the last response after a total elapsed time of 5 min. After a response, the next tone was played randomly after 2-10 s. False starts were coded for RTs <100 ms^139^ and lapses were coded for RTs≥2 times the median. After removing false starts and lapses, response speed (mean 1/RT) was calculated as the outcome of interest, as it was shown to be most sensitive for slight deviations in sleep pressure^90^. The first test was taken on arrival as a baseline, then 5 times during the protocol at 90-min intervals and the last at a 60-min interval, 30 min before scheduled laboratory bedtime (1 h after HBT).

### Skin temperature

Starting 7 h before HBT, we continuously monitored skin temperature with 6 surface skin temperature thermocouples^140^ (BS 1922L Thermochron iButton®, Maxim, US) with 60-s intervals and a 0.0625°C sensitivity. Sensors were placed with air-permeable surgical tape (Fixomull®; Beiersdorf, Hamburg, Germany) on proximal (2 probes, subclavicular area) and distal (4 probes, Skin temperatures (distal & proximal) and the distal-proximal skin temperature gradient (DPG) were calculated accordingly by the author M.D. All raw recordings were visually inspected for each subject and segments where the sensors were either removed or malfunctioning were cleaned out manually. DPG computation during the cleaned periods thus relies on the remaining working probes. Room temperature was monitored with 1 surface temperature thermocouple, also with 60-s intervals at a 0.0625°C sensitivity, placed in a small cotton sachet 10 cm below the right armrest of the participant’s chair. As for skin temperatures, these recordings were individually inspected to warrant any recording issues.

### Ambulatory light history

Light levels at the wrist, expressed in photopic illuminance, were measured at 1-min intervals during the ambulatory monitoring part of the protocol, ≥5 days before each experimental session, using wrist-worn actimeters (ActTrust 2, Condor Instruments, Brazil). Participants were instructed to wear the device continuously and on their non-dominant wrist without sleeves covering it, except when it might get wet or damaged. The recorded illuminance data were pre-processed using the Light Log R package^141^ in R and corrected by a factor of 1.22 to account for the average difference between the ActTrust 2 devices and a Jeti spectraval 1501 spectroradiometer (JETI Technische Instrumente GmbH, last calibration: 07.03.2023) measured under the same illuminant as in the experiment at different intensities. Based on the procedure in^142^, samples below 1 lx, likely due to sleeve covering as well as samples taken during the scheduled bedtime, were excluded from this analysis. Bright light history was used as a covariate in the mixed linear models to account for participants’ light exposure prior to laboratory entry and was quantified as the time above the 1000 lx illuminance threshold (TAT 1000 lx) from the previous day’s HWT to the scheduled laboratory session (∼32 h period). Similar periods (24-48 h) for prior light exposure have been used and found to be relevant in other studies^78,81,86^. The average light history during the day prior to each experiment, expressed as time over threshold at different illuminance thresholds, is shown in Figure 4. However, it is important to note that there are likely to be differences between light exposure at eye level and that measured at the wrist^143,144^.

### Pubertal stage and chronotype

Self-reported pubertal stage^145^ and chronotype^146,147^ (mid-sleep time corrected for oversleeping) were assessed as part of the screening procedure and considered as passive stratification covariates in the LMM analyses, as previous reports have shown associations with light-induced melatonin suppression^23^, melatonin onset^148^, and melatonin amplitude^149^.

### Additional measures

Furthermore, a visual comfort and well-being scale (VCS) was employed. An adapted German version of the first 6 items of a visual comfort scale^150^ was used to assess the participant’s visual comfort under the different lighting conditions. Additionally, participants rated the subjective room temperature and temperature preference with 2 custom questions, and their momentary affect and well-being in relation to mood, hunger, relaxation, and motivation with 4 questions based on the questionnaire used by Reichert et al.^151^. These self-reports were administered every 90 min together with the PVT and used to check on the well-being of the participants. Moreover, participants were asked to fill out a daily sleep and daytime diary^131^. At the end of the last experimental session, participants also filled in a questionnaire assessing their light exposure-related behaviour retrospectively^152^. Further measured outcomes that were not included in the analysis of this particular study were pupillary light responses^153^, polysomnography, Karolinska Drowsiness Tests^137^, self-reported sleep quality^132^ and dream recall^154^.

### Design, reproducibility, and statistics

In this within-subject design, all participants received the same 3 experimental conditions but in counterbalanced order. There was a washout period of ≥1 week (1 exception of 5 days) between the 3 in-lab sessions to minimise any carry-over effects of the previous sleep-wake behaviour and light condition. Experiments were typically scheduled on the same weekday for consistency. The counter-balanced random assignment to the condition sequence was conducted in separate strata for female and male participants with the “blockrand” package^155^ in *R*. In case of dropouts, the respective “new” participant was assigned to the same condition sequence as the participant who dropped out.

A total of 22 healthy adolescents (50% female, 50% male, target n=18 complete data sets) were assigned to one of 6 possible condition sequences. The 4 additionally recruited participants had the same experimental condition order as the first 4, which had missing baseline data and only a third of the samples for subjective sleepiness and missing baseline data for the PVT. Consequently, these first 4 participants were excluded from the subjective sleepiness analysis (n=18 complete datasets) but not the vigilant attention analysis (n=22 complete datasets), because the baseline samples were not processed in the LMM analyses. We excluded n=2 participants from the melatonin analyses because their maximum evening melatonin levels were lower than their diurnal melatonin fluctuations and no clear onset could be detected (see melatonin profiles in Suppl. Figure S9). The 9 samples taken during the late evening light condition formed the basis for the primary outcome analysis (melatonin AUC). Participants with single missing evening melatonin samples (n=2) were included and for AUC calculation, these missing values were estimated by linear interpolation, resulting in a sample of n=20 for evening melatonin analyses. In the melatonin onset (MO) analysis using the hockey-stick method^82^ the default “area of interest upper border” of 5 pg/ml had to be adjusted in n=6 individuals, and in 3 cases, single outlier values before MO (diurnal fluctuations) had to be excluded from the hockey-stick fitting. These deviations from the default setting were discussed between R.L. and C.C. and agreed upon in discussion with F.F.

For skin and room temperature, 1 participant had erroneous thermocouple data (resolution=0.5 instead of 0.0625 °C) and was hence excluded from the analysis. Otherwise, data from 12 single thermocouples (2 in dim light conditions, 1 in bright light condition, 8 in moderate light condition, and 1 room temperature) were compromised and therefore excluded from the analysis. For one participant, room temperature data was not recorded. The missing thermocouples were compensated by using the data from the corresponding sensor on the other side of the body, resulting in a total of n=21 participants’ DPG datasets for the LMM analysis. The first hour recorded at the start and after the offset of the light intervention (transitional periods) was not included in the LMM analyses. Thus, each LMM, including data recorded during and after the light intervention, respectively, contained a total of 7 30-min DPG bins. As described in the supplementary information, we found that the room temperature was considerably higher during and after the bright light intervention (see Suppl. Figure S2 D) compared to the moderate and dim light conditions and was also subjectively rated as slightly but noticeably higher (see Suppl. Figure S2 E, F). Thus, the DPG and skin temperature results of the bright light intervention need to be interpreted with caution, as the internal validity of the effect of the light level vs. temperature differences is not given.

#### Statistics

We conducted all statistical analyses in R (4.3.1, R Core Team, 2023)^156^, using the packages “lme4”^157^ and “lmerTest”^158^ to test our estimate-specific hypotheses with LMMs. Degrees of freedom for the fixed effects were estimated using Satterthwaite’s approximation and the model was fit using Restricted Maximum Likelihood (REML). As hypothesised a priori, we first tested whether our outcome measures during the later evening light exposure were significantly different (two-sided, p<0.05) when preceded by the “bright” or “moderate” light interventions, compared to “dim” light. Post-hoc hypotheses were analysed with the same model structure and procedure but – except for melatonin – were based on the data recorded *during* the AEE light interventions. LMM assumptions were visually inspected with the help of the “performance” package^159^ (see Supplementary Figures S10-S18), which indicated that they were largely met, with some slight violations of the ‘normality of residuals’ assumption. LMM analyses were nonetheless performed due to the robustness of LMMs to deviations in the distribution assumptions^160^. Two models were calculated for each of the different outcome variables: a “covariate” model including the theoretically relevant covariates (see full models below), and a “sparse” model including only the main effect(s) (see model variables in bold below). The LMMs had the following structure:

1) ***Melatonin AUC ∼ AEE light intervention condition** + bright light history + pubertal stage + chronotype +* (1| *participant*)
2) ***Secondary Outcome ∼ AEE light intervention condition * time (median − centred)** + bright light history + pubertal stage + chronotype +* (1| *participant*)

We report here, the estimates of the linear mixed model (LMM) analyses from the covariate model. Additional estimates for a sparse model without covariates are reported in Suppl. Tables S2, S3, S5, S7, and S9. The covariate of primary interest was “bright light history” (TAT 1000 lx) used as a secondary intervention variable, while the other covariates were used as passive stratification factors to control for additional variance. These estimates were not interpreted but are reported in the LMM result tables for completeness. The following variables were modelled as fixed effects: The light intervention conditions (categories “moderate” and “bright”, relative to “dim”), bright light history (numeric, “TAT 1000 lx”), self-reported pubertal stage (categories, relative to “post-pubertal stage”) and self-reported chronotype (numeric, midpoint of sleep corrected for oversleep, “MSF_SC_”). Repeated measures per participant were included as a random factor. For subjective sleepiness, vigilant attention and DPG, a (median centred) “time” variable was additionally included in the linear models together with the “time * AEE light intervention condition” interaction terms to account for the effects of the factor time during the protocol. Median centring for the “time” variable was performed for each tested period separately (during the AEE light condition and late evening light condition, respectively), so that “time=0” refers to the middle of each tested period. This prevented high variance inflation factors (VIFs) between the factor “time” and the “time * “AEE light intervention condition” interaction term.

## Code and data availability

The anonymised and de-identified datasets analysed in the current study, as well as a laboratory log file, will be made publicly available on Figshare under the CC BY-ND 4.0 licence after peer-reviewed publication, while the statistical analysis code using these datasets will be made publicly available on GitHub under the MIT licence.

## Acknowledgements

We would like to thank Johannes Zauner for his assistance in computing the wrist-worn bright light history using the LightLogR package^141^, and Christine Blume for her support and providing code for the PVT analysis. Furthermore, we are grateful to Kurt Kräuchi for his valuable comments on skin temperature regulation. We also extend our gratitude to all the study helpers and interns for their contributions to data collection.

## Author contributions

The authors made the following contributions. Rafael Lazar: Conceptualization, Data curation, Formal Analysis, Investigation, Methodology, Project administration, Supervision, Validation, Visualization, Writing –original draft, Writing – review & editing; Fatemeh Fazlali: Conceptualization, Formal Analysis, Methodology, Writing – review & editing; Marine Dourte: Formal Analysis, Writing – review & editing; Christian Epple: Investigation, Writing – review & editing; Oliver Stefani: Conceptualization, Methodology, Supervision, Writing – review & editing; Manuel Spitschan: Conceptualization, Formal Analysis, Methodology, Supervision, Writing – review & editing; Christian Cajochen: Conceptualization, Formal Analysis, Funding acquisition, Methodology, Project administration, Resources, Supervision, Visualization, Writing – review & editing.

