## Supplementary information for "Afternoon to early evening bright light exposure reduces later melatonin production in adolescents"

#### Supplementary Methods

##### Detailed recruitment information

The selection criteria, screening instruments used, and corresponding cut-off points are shown in Suppl. Table S1, while number of exclusions and withdrawals with their reasons are shown and described in Suppl. Fig. S5. All questionnaires and instructions during the screening and study procedures were administered in German language. If an individual was excluded for non-adherence during the sleep-wake rhythm stabilisation period but had already completed one or more study visits, none of their data were used. A total of 82 students initially expressed interest in participating in the study, of whom 52 attended the mandatory telephone information session. 41 students and their parents then provided written informed consent. 37 of them completed the online screening questionnaire implemented on the online tool REDCap[1], of which 28 were invited to participate in the face-to-face screening at our facilities, which included a physical examination by the study physician (author C.E.). A total of 27 adolescents participated in the protocol, of whom 5 did not complete the study. 2 were excluded because of non-compliance with the agreed sleep-wake times, 1 was excluded because of inability to follow instructions and 2 dropped out after completing the first session because the protocol was too much of a burden alongside school. We recruited an additional 4 participants due to incomplete subjective sleepiness data from the first 4 participants to reach the target sample size of n=18 complete data sets. A total of 22 participants completed the study. Full participation in the study was compensated with 550 (CHF), or a partial amount in case of dropout.

**Morning light condition**
Participants were awakened by switching on a dimmed warm white, fluorescent light source (~2800K, ~1.5 lx illuminance, 0.6 lx mEDI at eye level while lying in bed) and 5 min later were seated at a table where they remained for 30 min illuminated by the same light source (~3.9 lx illuminance, 1.6 lx mEDI at eye level while seated).

### Supplementary results

**Morning melatonin results**
As an exploratory post-hoc analysis, we additionally tested the AUC of the 2 morning samples recorded within 35 min of waking with the same LMM structure. Here, only n=59 observations were included because of a missing morning sample. Neither of the predictors in the LMM yielded significant effects (see Suppl. Table S3).

**Phase angle between melatonin onset and habitual bedtime**
The phase angle between melatonin onset times (MO) and habitual bedtime (HBT) showed considerable variance (mean= 81 min, SD=69 min; median=96 min, Q25=8, Q75=124) ranging from a maximum of 230 min to a minimum of -25 min. In total, 5 MOs from 3 individuals (out of 60 included MOs) yielded a phase angle >180 min, i.e. their MO was already detected during the light intervention period. For these individuals, the moderate and bright light intervention may not only have acutely delayed their phase but also may have acutely suppressed their melatonin release. However, our primary outcome (evening melatonin area under the curve, AUC) only included the data in the later evening light condition, collected *after* the afternoon to ealy evening (AEE) light intervention. On the other hand, 6 MOs from 4 individuals yielded a phase angle<0, i.e. their MO was detected after HBT. Thus, despite the individualised timing of the study with respect to HBT, which the participants were required to keep consistent for 5 days prior to each laboratory session, the light interventions were still applied at different "biological times" with respect to MO.

**Room temperature results**
We further examined the recorded room temperature across the different conditions, given the contrasting results during and after the bright and moderate light interventions. Suppl. Fig. 2 panel D shows a clear difference in room temperature during and after the bright light intervention (mean=23.18 °C, *SD*=0.40 °C) compared to during and after the moderate light intervention (mean=22.29 °C, *SD*=0.83 °C) and dim light (mean=22.45 °C, *SD*=0.74 °C). In all conditions, there is a peak in room temperature just after 3 h before the HBT, which is related to the presence of several people in the laboratory to attach the EEG cap to the participant's head. The higher room temperature during and after the bright light intervention was also reflected in the subjective room temperature ratings shown in Suppl. Fig. S2, panels E and F, where subjects rated the room as warmer on average and showed a slightly higher preference for a 'cooler' room than during and after the dim and moderate light interventions. The ratings for the dim and moderate light interventions largely overlapped, again reflecting the objectively measured room temperatures for these conditions.

#### Supplementary Figures

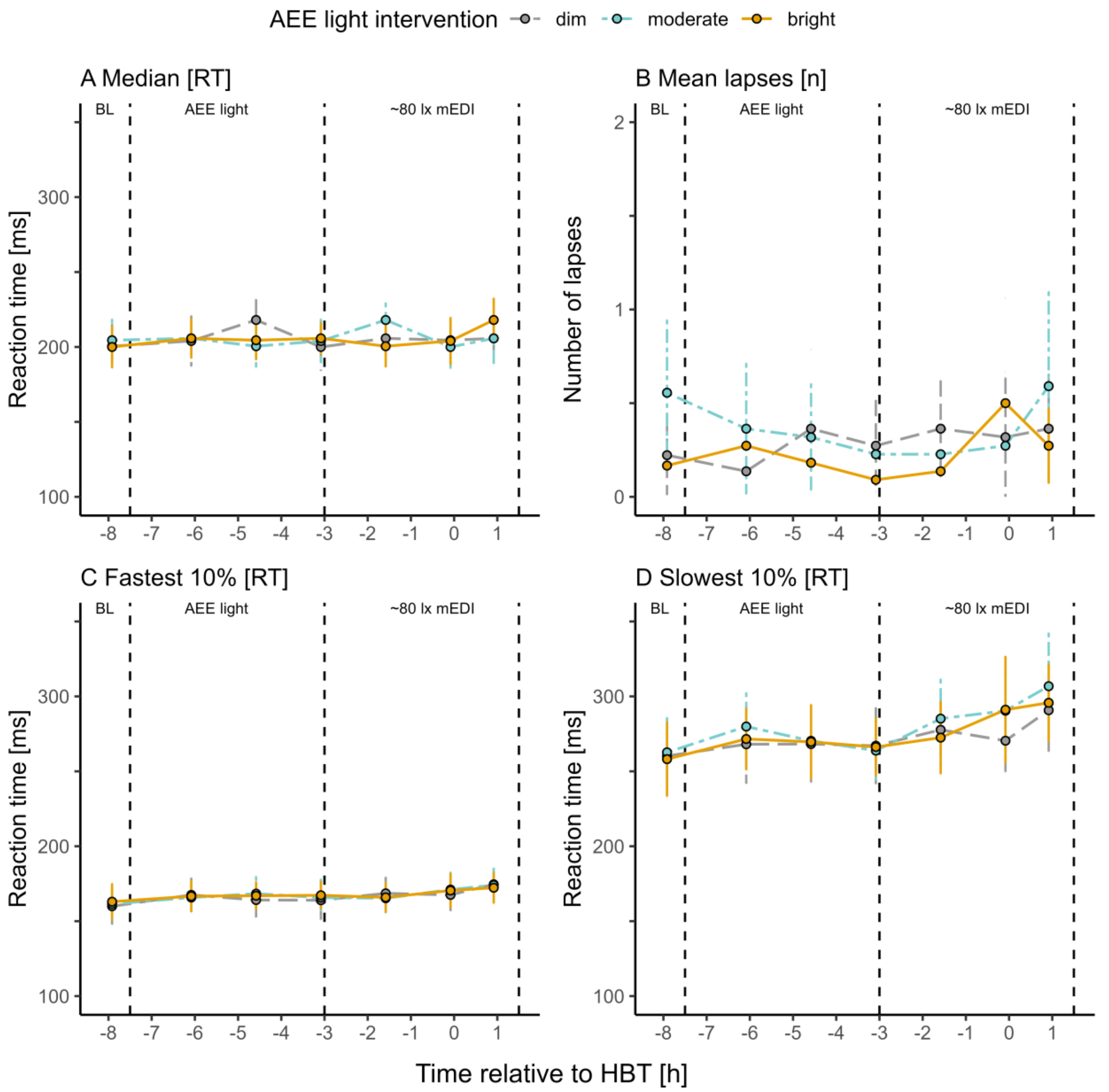


**Figure S1. Time course of other PVT outcomes across the protocol (relative to HBT).** The graphs are based on PVT reaction time data from 22 participants (0 excluded). Values at baseline (BL) are only based on 18 participants. HBT (0 on the x-axis) on average corresponded to 22:15 (SD=31 min). **A**, Coloured dots show the median reaction time per condition, with error bars indicating the 95% confidence interval. **B**, Coloured dots show the mean lapses per condition, with error bars indicating the 95% confidence interval. **C**, Coloured dots show the mean fast 10% per condition, with error bars indicating the 95% confidence interval. **D**, Coloured dots show the mean slowest 10% per condition, with error bars indicating the 95% confidence interval. Abbreviations: AEE = afternoon to early evening, BL = Baseline measurement, mEDI = melanopic Equivalent Daylight Illuminance, HBT = Habitual bedtime (for home sleep), n = number of lapses, PVT = Psychomotor Vigilance Task, RT = Reaction time.

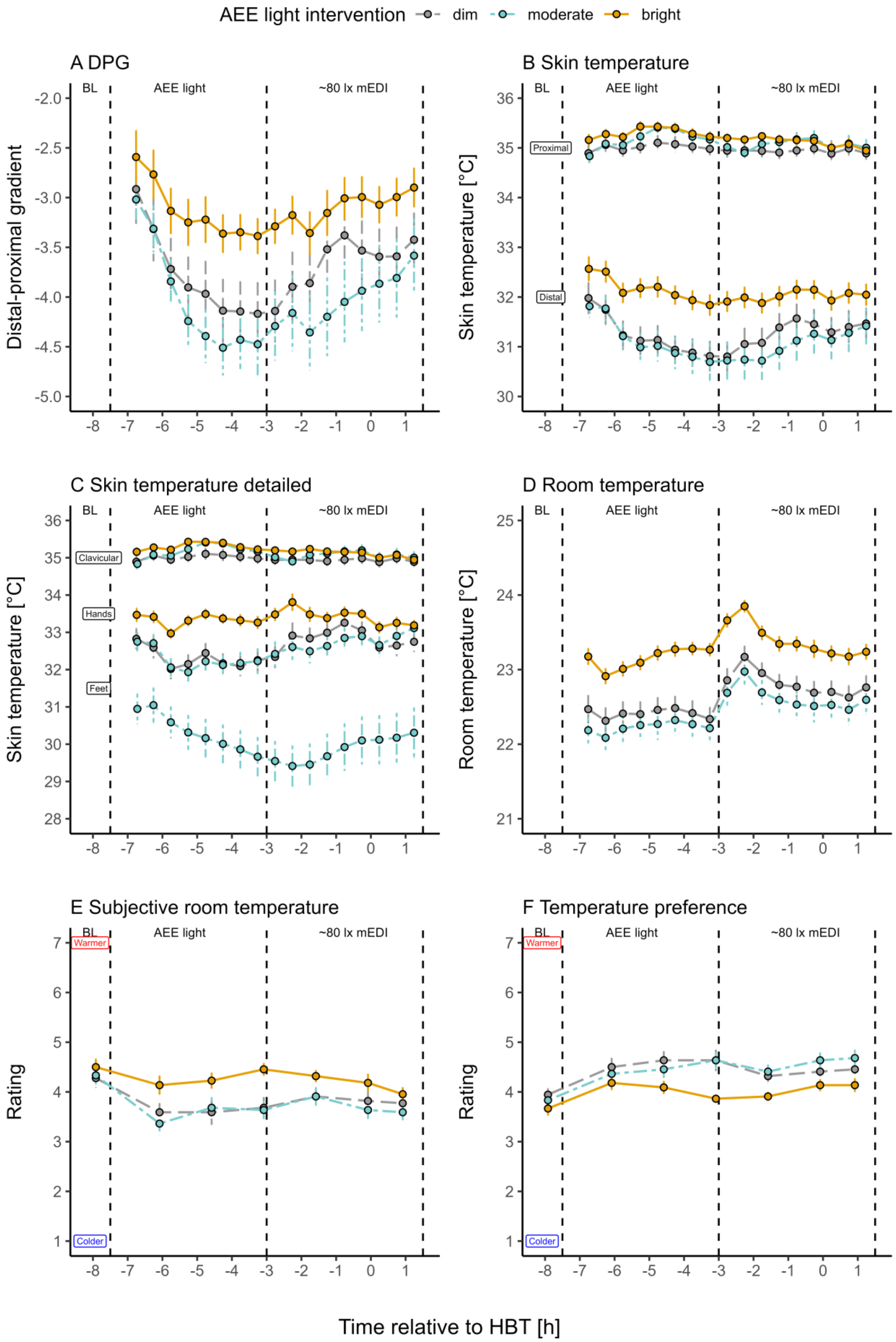
AEE light intervention dim moderate bright
A DPG
B Skin temperature
C Skin temperature detailed
D Room temperature
E Subjective room temperature
F Temperature preference
BL
AEE light
~80 lx mEDI
Distal-proximal gradient
Skin temperature [°C]
Room temperature [°C]
Rating
Proximal
Distal
Clavicular
Hands
Feet
Warmer
Colder
Time relative to HBT [h]

**Figure S2. Time course of outcomes related to temperature regulation across the protocol (relative to HBT).** HBT (0 on the x-axis) on average corresponded to 22:15 (SD=31 min). **A**, DPG throughout the experiment based on data from 21 participants (1 excluded). Coloured dots show the mean DPG in 30-min bins per condition, with error bars indicating the standard error. The 1st time bin (transition sample only used for plotting) is based on 19 single datapoints during the moderate light intervention,18 single datapoints for the "bright" light intervention and 17 single datapoints for dim light. **B**, Distal and proximal skin temperature data throughout the experiment based on data from 21 participants (1 excluded). Coloured dots show the mean temperature in 30-min bins per condition, with error bars indicating the standard error. The first mean time bins are based on 19 single datapoints during the moderate light intervention,18 single datapoints for the "bright" light intervention and 17 single datapoints for dim light. **C**, Skin temperature data for the different locations in the proximal (clavicular) and distal (hands, feet) regions of the body throughout the experiment based on data from 21 participants (1 excluded). Coloured dots show the mean temperature in 30-min bins per condition, with error bars indicating the standard error. The first mean time bins are based on 19 single datapoints during the moderate light intervention,18 single datapoints for the "bright" light intervention and 17 single datapoints for dim light. **D**, Coloured dots show the mean room temperature in 30-min bins per condition, with error bars indicating the standard error. Room temperature values are based on data from 20 participants (1 excluded, 1 missing) for the moderate and bright light intervention and 19 participants for the dim light. The first afternoon time bins are based on 18 single datapoints during the moderate light intervention,17 single datapoints for the "bright" light intervention and 15 single datapoints for dim light. **E**, Subjective ratings of room temperature based on 22 participants' replies to the question "How do you find the room temperature?" from "1 - very cold" to "7 - very warm". Values at baseline (BL) were based on 18 participants. Coloured dots show the mean values per condition, error bars show the standard error. **F**, Subjective ratings of room temperature preference based on 22 participants' replies to the item "I would prefer it to be..." with responses ranging from "1 - much colder" to "7 - much warmer". Values at baseline (BL) were based on 18 participants. Coloured dots show the mean per condition, error bars show the standard error. Abbreviations: AEE = afternoon to early evening, BL = Baseline measurement, DPG = Distal-proximal skin temperature gradient, mEDI = melanopic Equivalent Daylight Illuminance, HBT = Habitual bedtime (for home sleep).

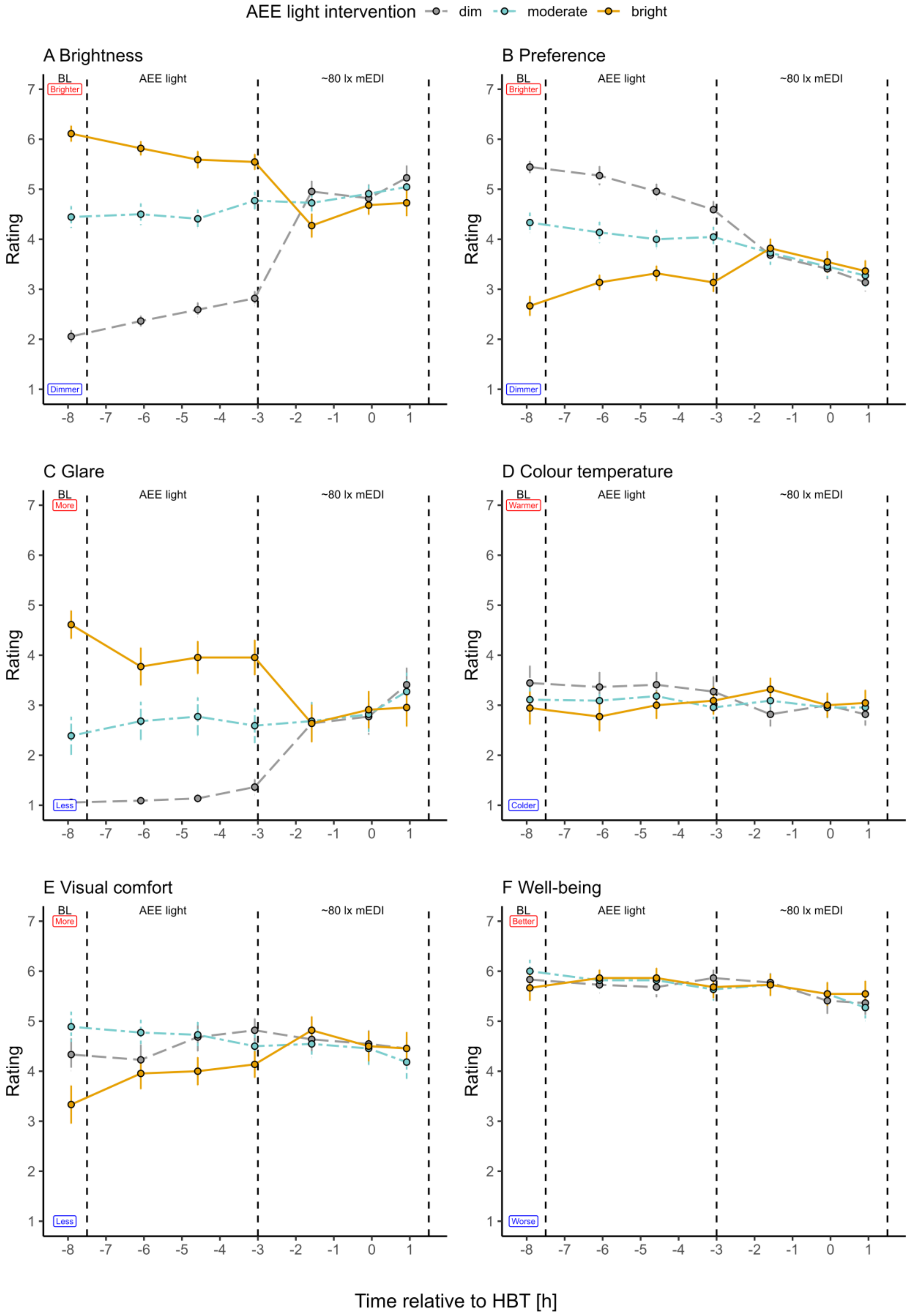
AEE light intervention
dim
moderate
bright
A Brightness
B Preference
C Glare
D Colour temperature
E Visual comfort
F Well-being
BL
AEE light
~80 lx mEDI
Brighter
Dimmer
More
Less
Warmer
Colder
Better
Worse
Rating
Time relative to HBT [h]

**Figure S3. Time course of the visual comfort scale and well-being ratings across the protocol (relative to HBT).** Subjective ratings as part of the visual comfort and well-being scale (VCS), based on 22 participants. Values at baseline (BL) were only based on 18 participants. HBT (0 on the x-axis) on average corresponded to 22:15 (SD=31 min). **A**, Subjective ratings of brightness of the lighting based on replies to the item "How do you find the brightness of the light?" with responses ranging from "1 – very dark" to "7 – very bright". Coloured dots show the mean per condition, error bars show the standard error. **B**, Subjective ratings of brightness preference of the lighting based on replies to the item "I would prefer it to be..." with responses ranging from "1 – much darker" to "7 – much brighter". Coloured dots show the mean per condition, error bars show the standard error. **C**, Subjective ratings of glare of the lighting based on replies to the item "This light is glaring." with responses ranging from "1 – not at all" to "7 – very much". Coloured dots show the mean per condition, error bars show the standard error. **D**, Subjective ratings of the colour temperature of the lighting based on replies to the item "How do you feel the light colour is?" with responses ranging from "1 – very cold" to "7 – very warm". Coloured dots show the mean per condition, error bars show the standard error. **E**, Subjective ratings of the visual comfort of the lighting based on replies to the item "In general, the light is pleasant." with responses ranging from "1 – not at all" to "7 – very much". Coloured dots show the mean per condition, error bars show the standard error. **F**, Subjective ratings of well-being based on replies to the item "How do you feel at the moment?" with responses ranging from "1 – unwell" to "7 – well". Coloured dots show the mean per condition, error bars show the standard error. Abbreviations: AEE = afternoon to early evening, BL = Baseline measurement, mEDI = melanopic Equivalent Daylight Illuminance, HBT = Habitual bedtime (for home sleep).

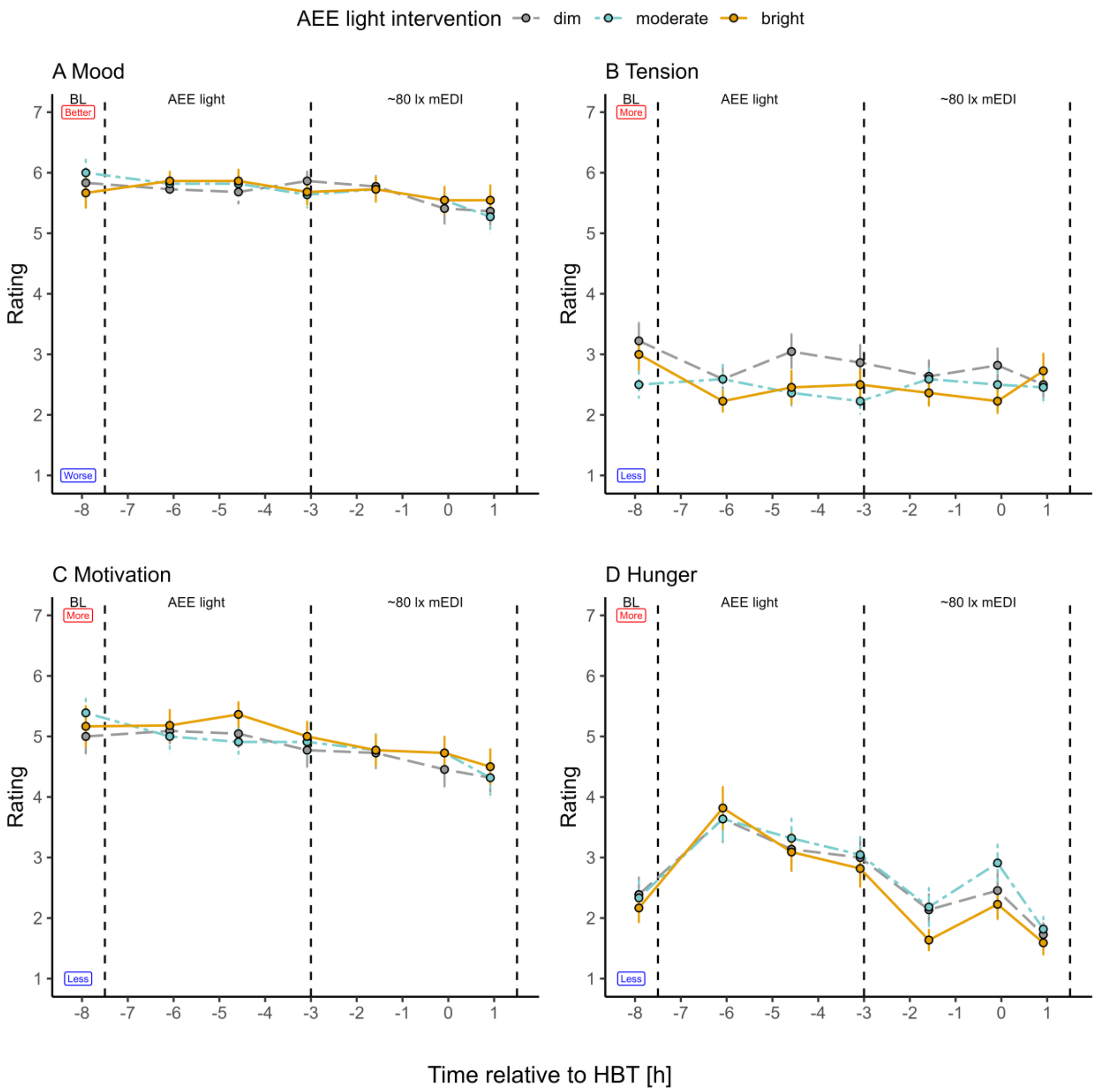


**Figure S4. Time course of subjective ratings of mood, relaxation, motivation and hunger across the protocol (relative to HBT).** Subjective ratings as part of the VCS, based on 22 participants. Values at baseline (BL) were only based on 18 participants. HBT (0 on the x-axis) on average corresponded to 22:15 (SD=31 min). **A**, Subjective ratings of mood based on replies to the item “How do you feel at the moment?” with responses ranging from "1 –very bad mood" to "7 – very good mood". Coloured dots show the mean per condition, error bars show the standard error. **B**, Subjective ratings of tension based on replies to the item “How do you feel at the moment?” with responses ranging from "1 – very relaxed" to "7 – very tense". Coloured dots show the mean per condition, error bars show the standard error. **C**, Subjective ratings of motivation based on replies to the item “How do you feel at the moment?” with responses ranging from "1 – very unmotivated" to "7 – very motivated ". Coloured dots show the mean per condition, error bars show the standard error. **D**, Subjective ratings of hunger based on replies to the item “How do you feel at the moment?” with responses ranging from "1 – very sated" to "7 – very hungry ". Coloured dots show the mean per condition, error bars show the standard error. Abbreviations: AEE = afternoon to early evening, BL = Baseline measurement, mEDI = melanopic Equivalent Daylight Illuminance, HBT = Habitual bedtime (for home sleep), VCS = visual comfort and well-being scale.

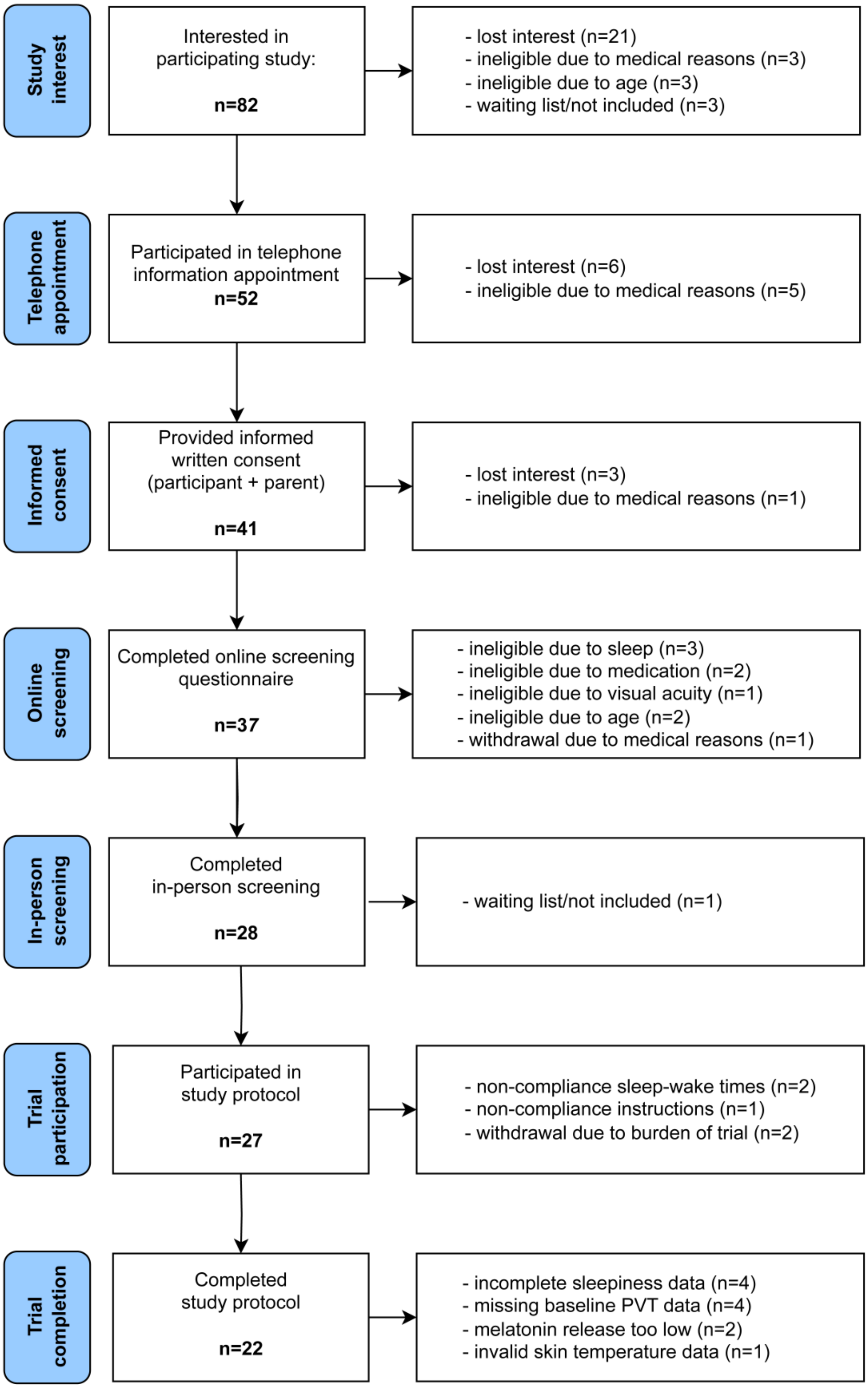


**Figure S5. Study flow diagram from study interest to trial completion.** A total of 82 students initially expressed interest in participating in the study, of whom 52 attended the mandatory telephone information session. 41 students and their parents then provided written informed consent. 37 of them completed the online screening questionnaire implemented on the online tool REDCap[1], of which 28 were invited to participate in the face-to-face screening at our facilities, which included a physical examination by the study physician (author C.E.). A total of 27 adolescents participated in the protocol, of whom 5 did not complete the study. 2 were excluded because of non-compliance with the agreed sleep-wake times, 1 was excluded because of inability to follow instructions and 2 dropped out after completing the first session because the protocol was too much of a burden alongside school. We recruited an additional 4 participants due to incomplete subjective sleepiness data from the first 4 participants to reach the target sample size of n=18 complete data sets. A total of 22 participants completed the study.

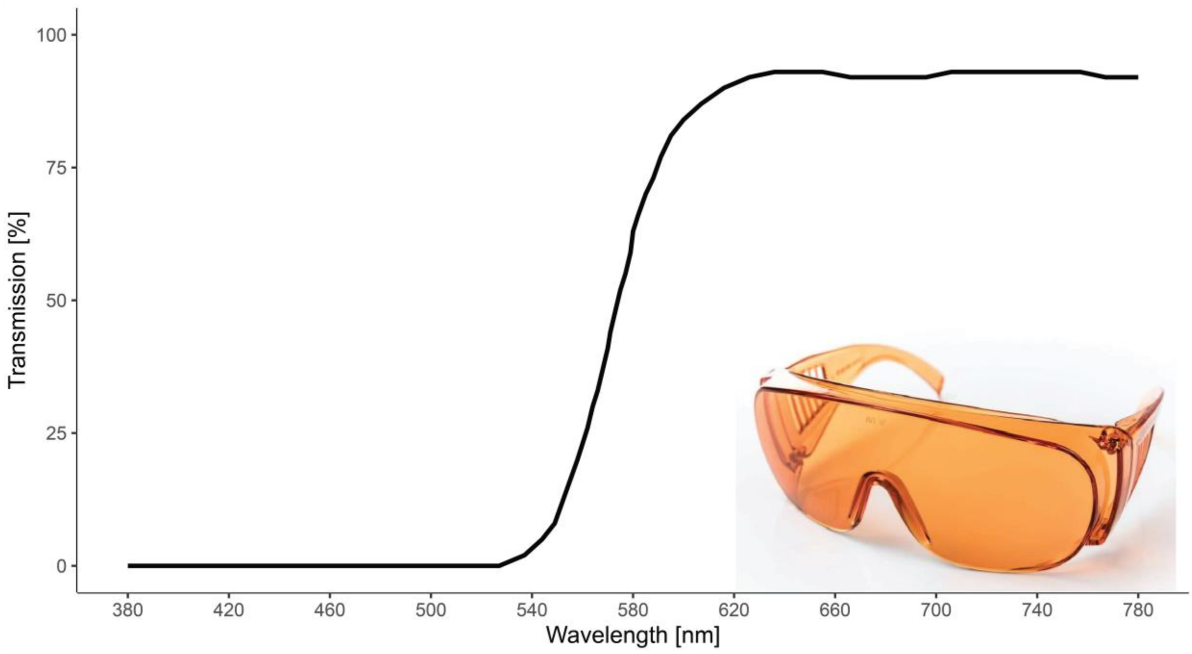


**Figure S6. Transmission of short-wavelength filter glasses applied out of the laboratory.** Participants wore red-orange tinted short-wavelength filter goggles when exposed to light outside the laboratory during toilet/walking breaks and on the way to the pupillograph in the neighbouring laboratory during the experiment. The respective light conditions in the corridor and bathroom, filtered and unfiltered, are specified in Suppl. Table S10. The graph shows the transmittance data in % per wavelength in the visual light spectrum (380-780 nm).

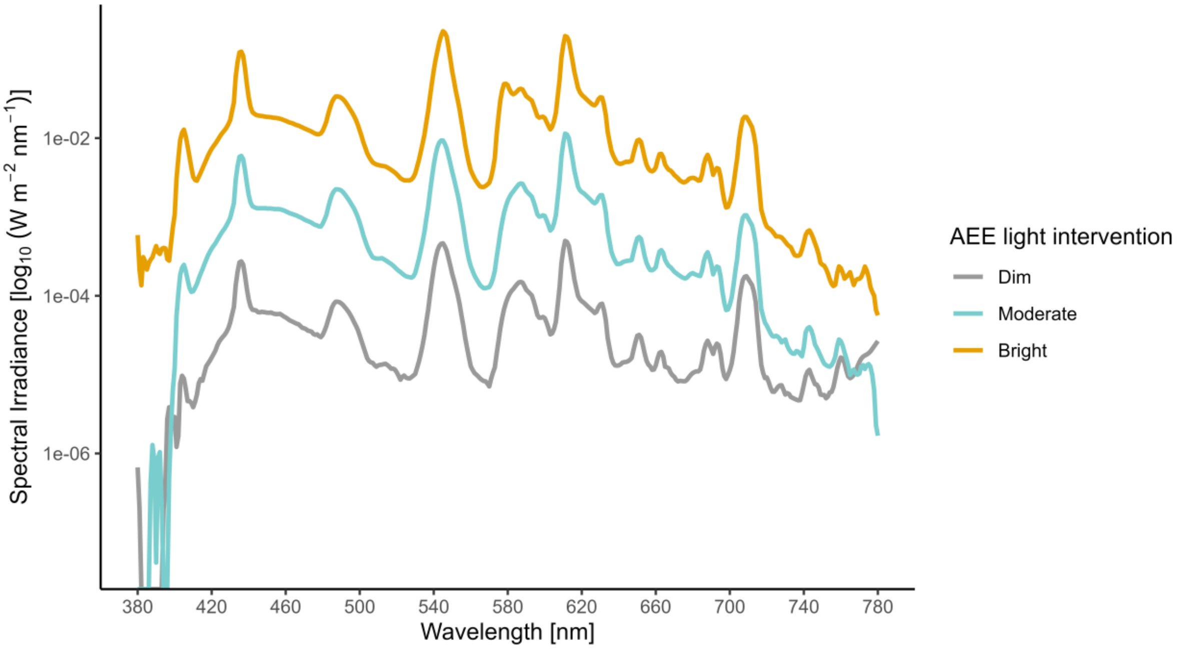


**Figure S7. Spectral power distribution of the light conditions in $log_{10}$ scale.** Distribution of spectral irradiances of the 3 AEE light exposure interventions during task-free periods (curtains closed). The vertical light measurements were taken with a spectroradiometer placed at the eye level of the seated participants (115 cm from the floor, 85 cm from the overhead light source, 80 cm from the white curtain, Spectraval 1501, JETI Technische Instrumente GmbH, Jena, Germany, last calibration: 07.03.2023).

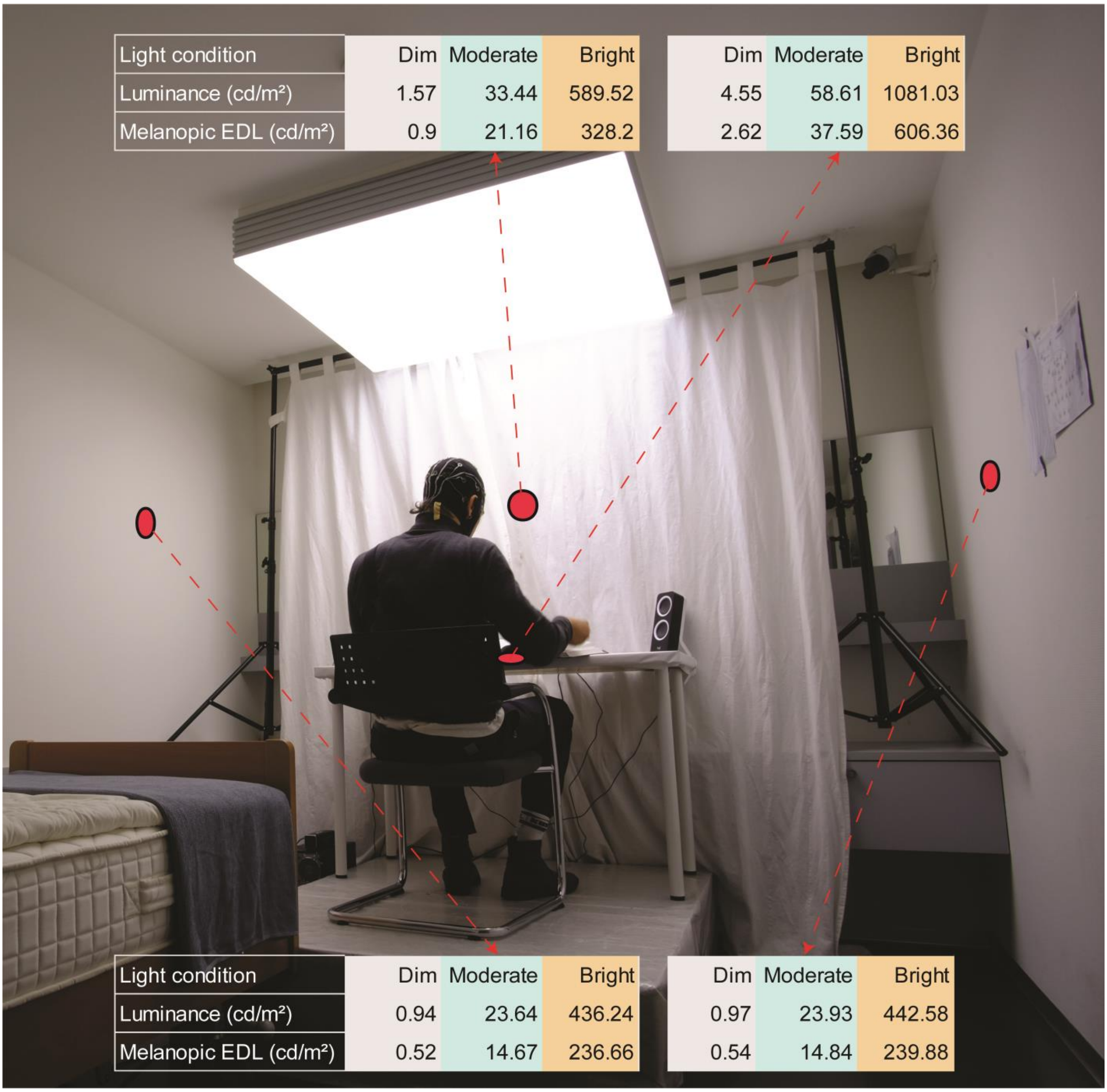


**Figure S8. Laboratory setup with stimulus radiances.** Summary of luminance and melanopic Equivalent Daylight Luminance (EDL), calculated with the luox application[2] from spectral radiance measurements of different object surface locations in the laboratory setup. Measurements correspond to task-free periods (curtains closed) under the 3 different AEE light intervention conditions. Further details of these stimulus radiances can be found in Suppl. Table 13. The subsequent evening light condition had the same characteristics as the “moderate” light intervention. Participants were instructed and monitored to remain seated with their eyes open and their sitting posture facing forward. The individual depicted is author R.L. Spectral radiance was measured from the observer's point of view using a research-grade spectroradiometer (Spectraval 1501, JETI Technische Instrumente GmbH, Jena, Germany). The sensor was placed at the eye level of the seated participants (115 cm from the floor, 85 cm from the overhead light source, 80 cm from the white curtain in front of them).

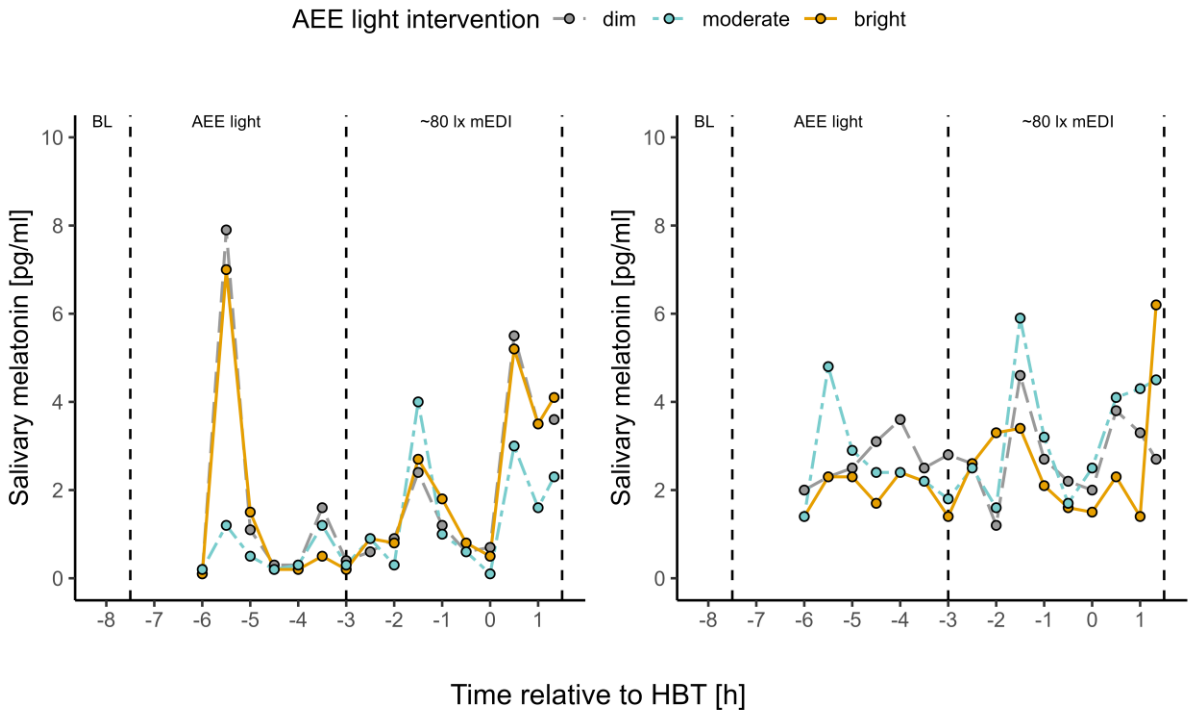


**Figure S9. Time courses of salivary melatonin concentrations (pg/ml) of participants excluded from the analysis.** Coloured dots show the measured melatonin levels per condition (AEE intervention) as a function of time (hours relative to HBT) with one panel per individual. Due to low melatonin release and high diurnal fluctuations, no clear salivary melatonin onset was detectable for these 2 participants. HBT (0 on the x-axis) corresponded to 22:15 (left panel) and 22.00 (right panel), respectively. Abbreviations: AEE = afternoon to early evening, BL = Baseline measurement, mEDI = melanopic Equivalent Daylight Illuminance, HBT = Habitual bedtime (for home sleep).

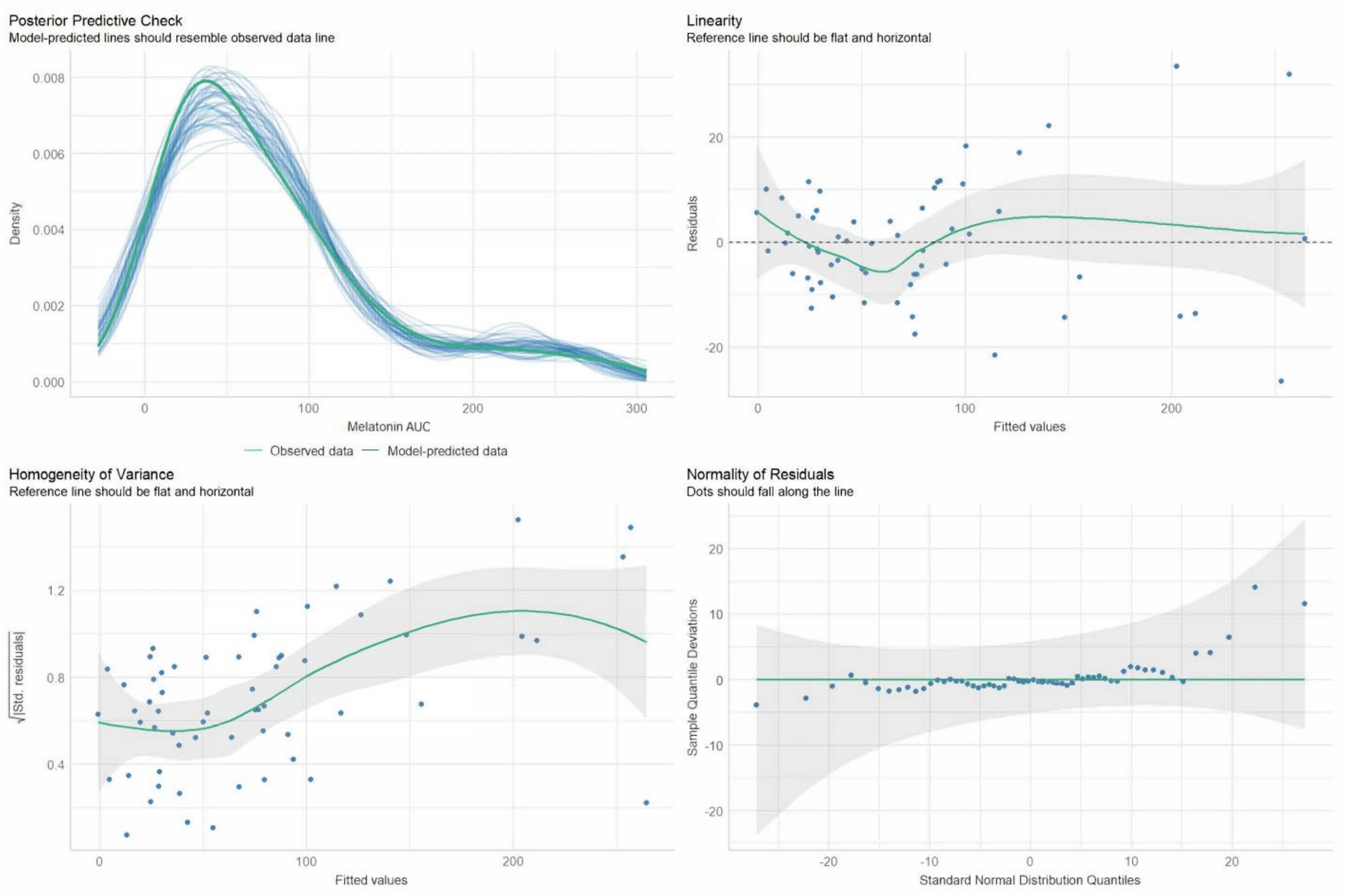


**Figure S10. Linear mixed model assumption checks for evening melatonin AUC.** Four different assumption checks were visualised for inspection: 1) Posterior predictive check comparing the density of predicted evening melatonin AUC data with observed data. 2) Linearity check, plotting residuals as a function of fitted values. 3) Homogeneity of variance (homoscedasticity) check, plotting the square root of standardized residuals as a function of fitted values. 4) Normality of Residuals check. In case of slight violations of these assumptions, the analyses were still performed due to the robustness of LMMs to deviations in distribution assumptions[3]. Abbreviations: AUC = Area under the curve, Std. residuals = Standardized residuals.

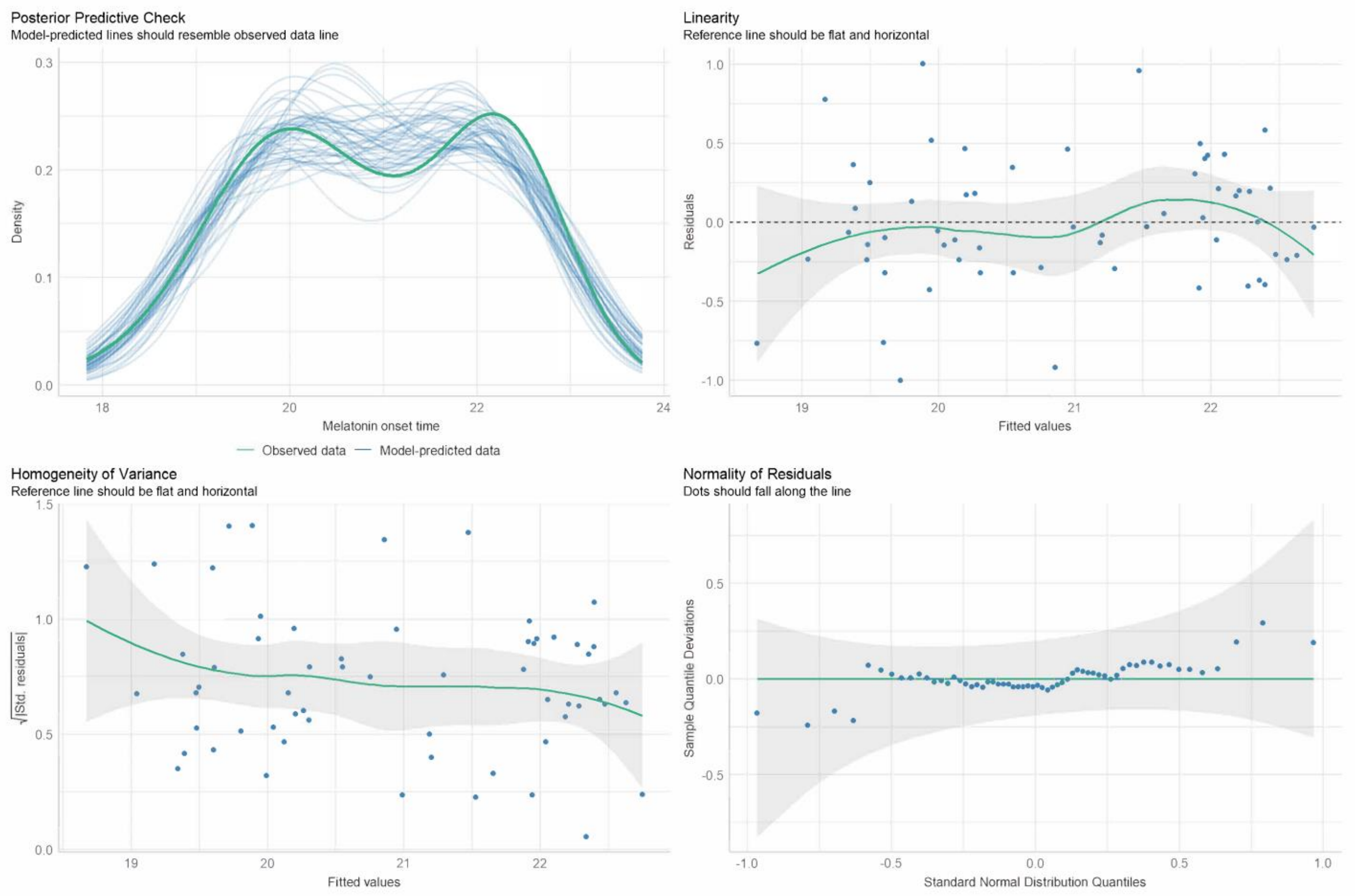


**Figure S11. Linear mixed model assumption checks for melatonin onset time.** Four different assumption checks were visualised for inspection: 1) Posterior predictive check comparing the density of predicted melatonin onset time data with observed data. 2) Linearity check, plotting residuals as a function of fitted values. 3) Homogeneity of variance (homoscedasticity) check, plotting the square root of standardized residuals as a function of fitted values. 4) Normality of Residuals check. In case of slight violations of these assumptions, the analyses were still performed due to the robustness of LMMs to deviations in distribution assumptions[3]. Abbreviations: Std. residuals = Standardized residuals.

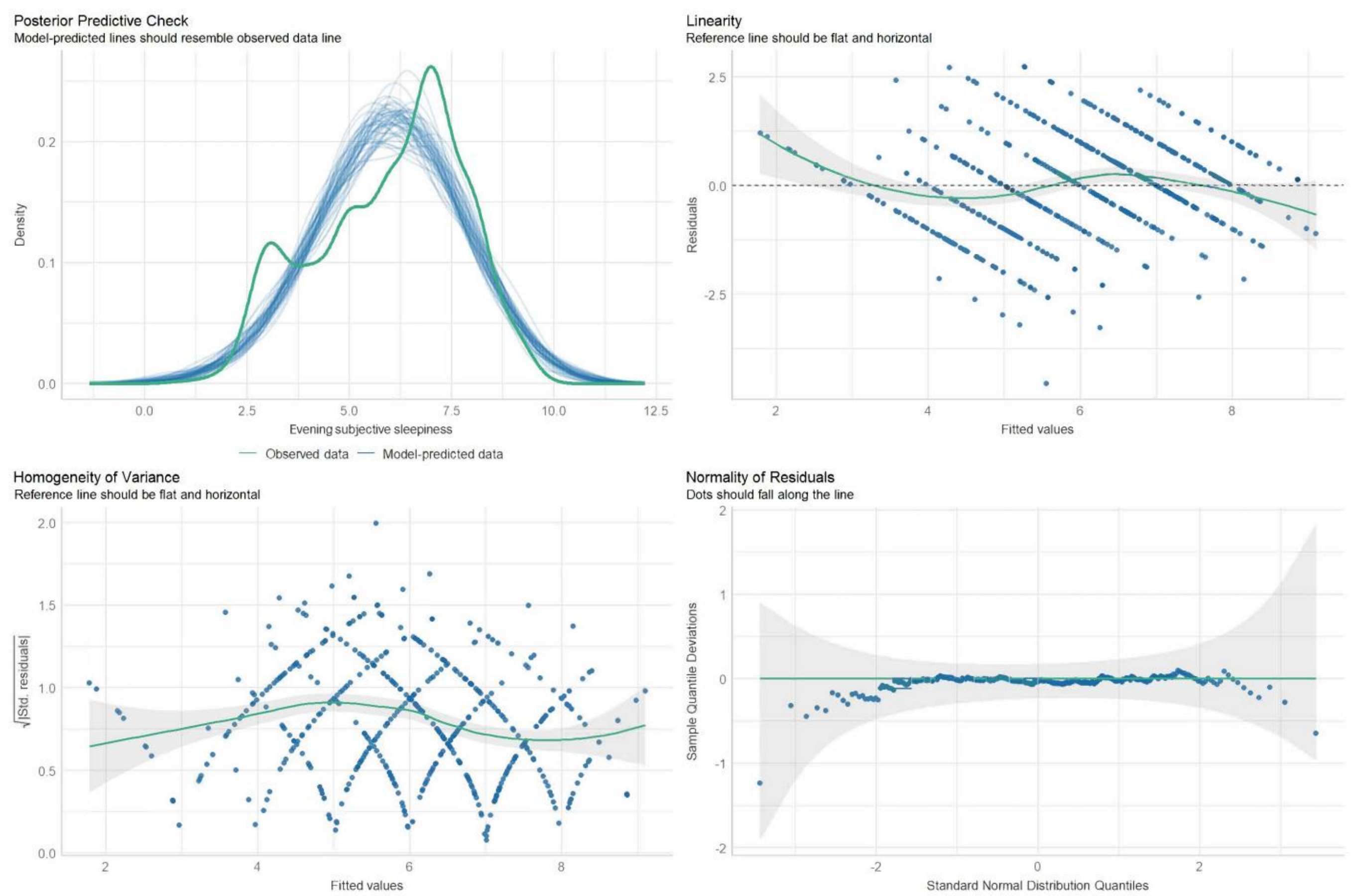


**Figure S12. Linear mixed model assumption checks for evening subjective sleepiness.** Four different assumption checks were visualised for inspection: 1) Posterior predictive check comparing the density of predicted later evening subjective sleepiness (KSS) data with observed data. 2) Linearity check, plotting residuals as a function of fitted values. 3) Homogeneity of variance (homoscedasticity) check, plotting the square root of standardized residuals as a function of fitted values. 4) Normality of Residuals check. In case of slight violations of these assumptions, the analyses were still performed due to the robustness of LMMs to deviations in distribution assumptions[3]. Abbreviations: KSS = Karolinska Sleepiness Scale, Std. residuals = Standardized residuals.

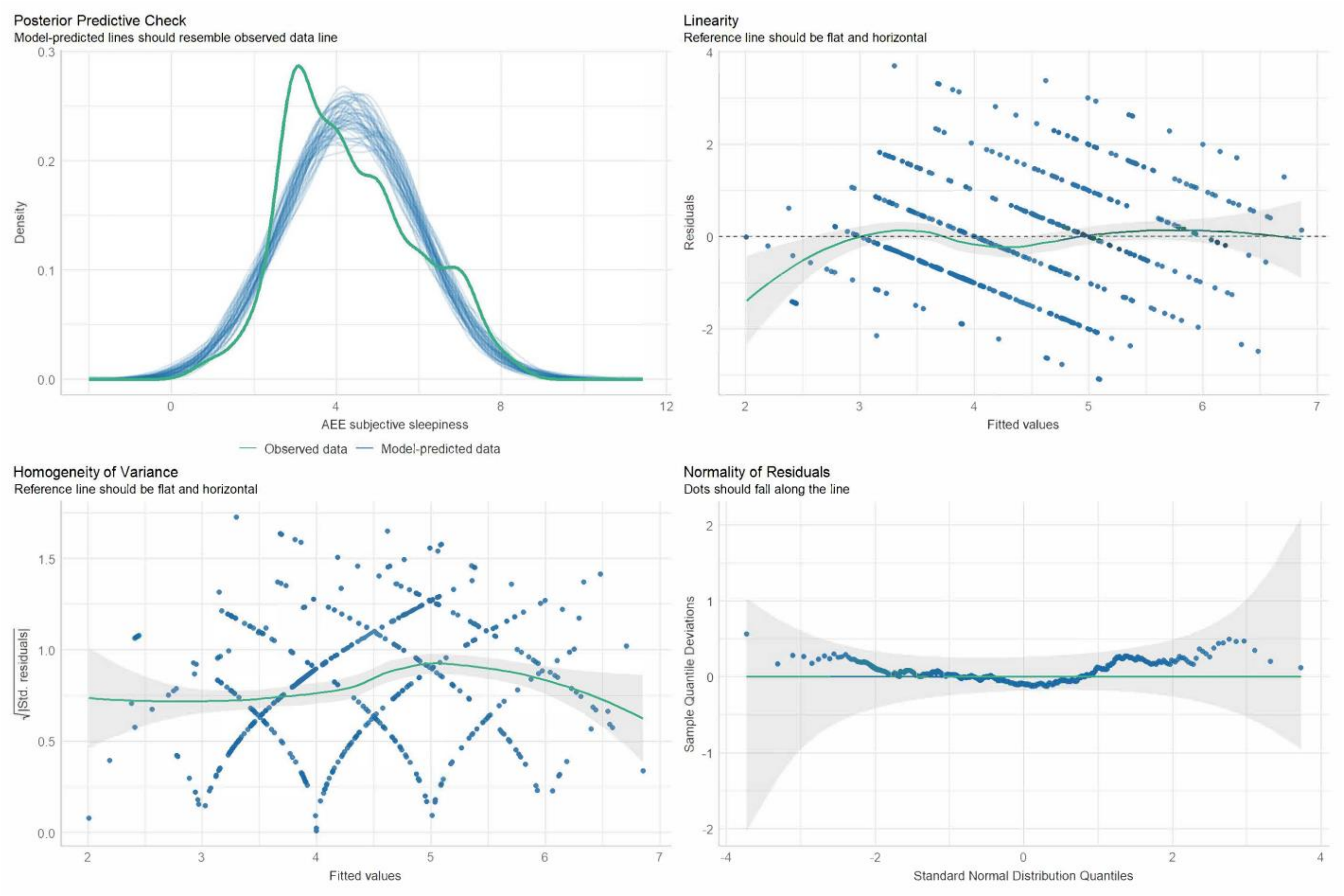


**Figure S13. Linear mixed model assumption checks for subjective sleepiness during the AEE interventions.** Four different assumption checks were visualised for inspection: 1) Posterior predictive check comparing the density of predicted subjective sleepiness data during the AEE light interventions with observed data. 2) Linearity check, plotting residuals as a function of fitted values. 3) Homogeneity of variance (homoscedasticity) check, plotting the square root of standardized residuals as a function of fitted values. 4) Normality of Residuals check. In case of slight violations of these assumptions, the analyses were still performed due to the robustness of LMMs to deviations in distribution assumptions[3]. Abbreviations: AEE = afternoon to early evening, KSS = Karolinska Sleepiness Scale, Std. residuals = Standardized residuals.

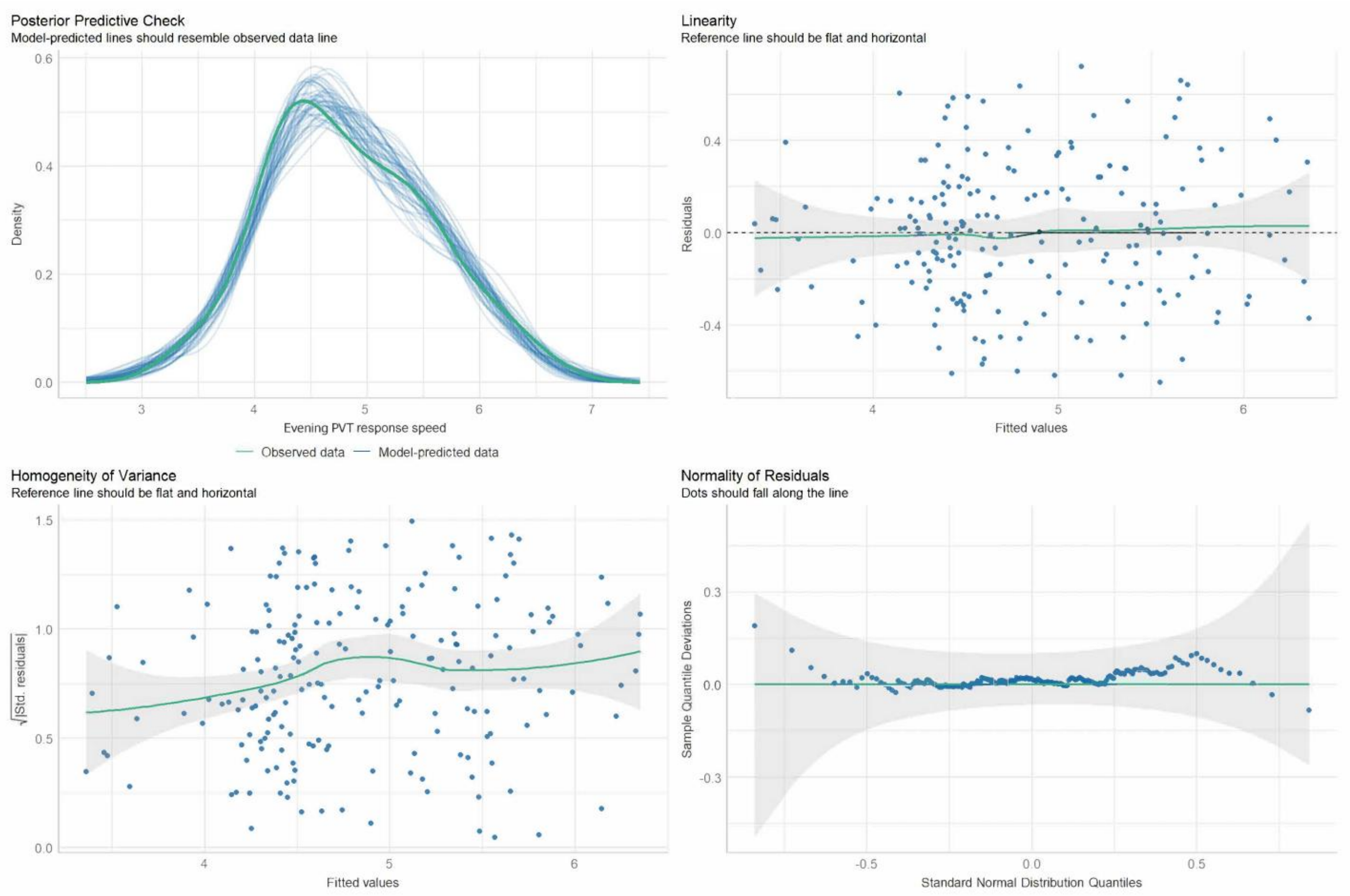


**Figure S14. Linear mixed model assumption checks for evening PVT response speed.** Four different assumption checks were visualised for inspection: 1) Posterior predictive check comparing the density of predicted later evening PVT response speed with observed data. 2) Linearity check, plotting residuals as a function of fitted values. 3) Homogeneity of variance (homoscedasticity) check, plotting the square root of standardized residuals as a function of fitted values. 4) Normality of Residuals check. In case of slight violations of these assumptions, the analyses were still performed due to the robustness of LMMs to deviations in distribution assumptions[3]. Abbreviations: PVT = Psychomotor Vigilance Task, Std. residuals = Standardized residuals.

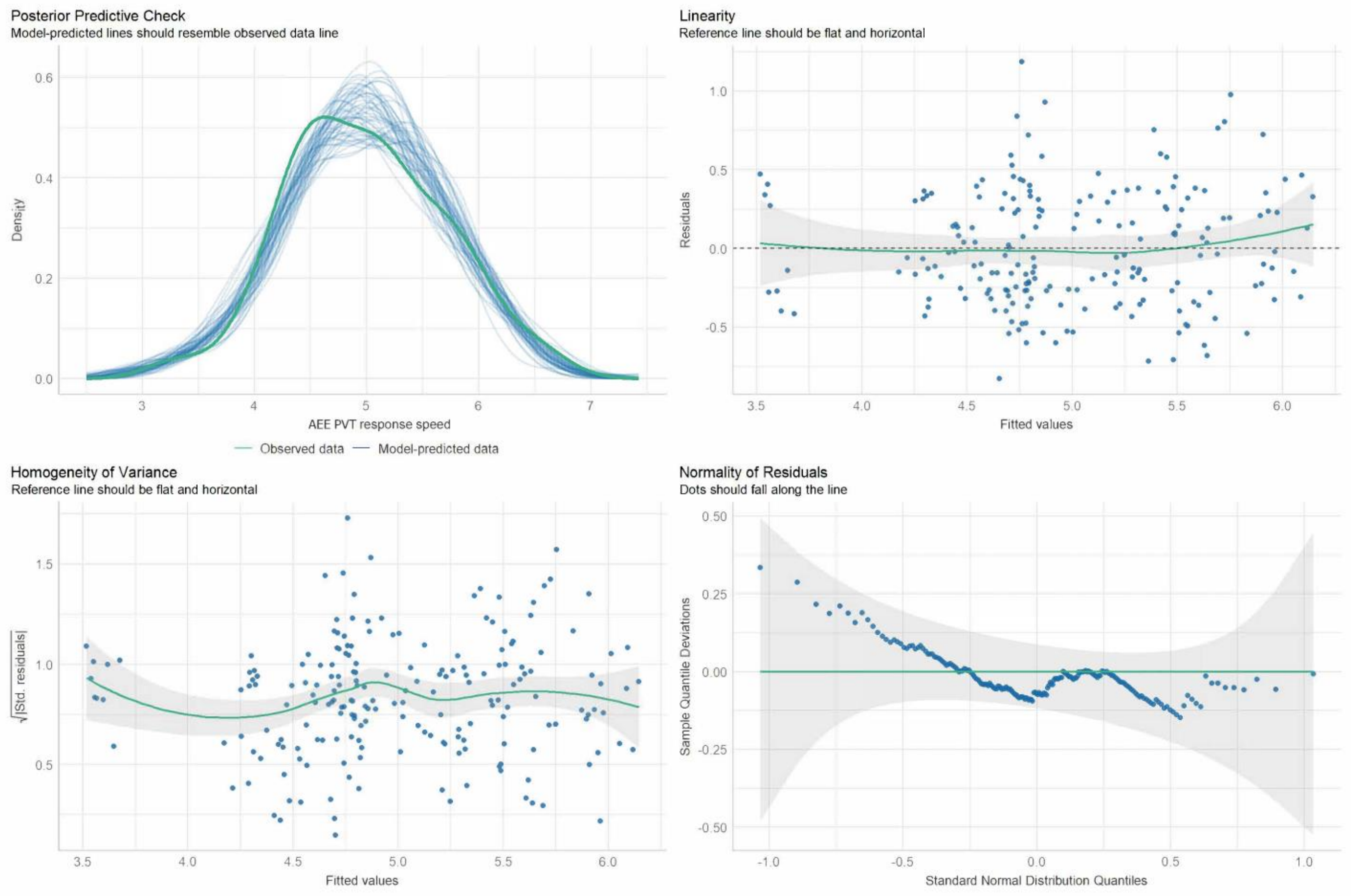


**Figure S15. Linear mixed model assumption checks for PVT response speed during the AEE interventions.** Four different assumption checks were visualised for inspection: 1) Posterior predictive check comparing the density of predicted PVT response speed during the AEE light interventions with observed data. 2) Linearity check, plotting residuals as a function of fitted values. 3) Homogeneity of variance (homoscedasticity) check, plotting the square root of standardized residuals as a function of fitted values. 4) Normality of Residuals check. In case of slight violations of these assumptions, the analyses were still performed due to the robustness of LMMs to deviations in distribution assumptions[3]. . Abbreviations: AEE = afternoon to early evening, PVT = Psychomotor Vigilance Task, Std. residuals = Standardized residuals.

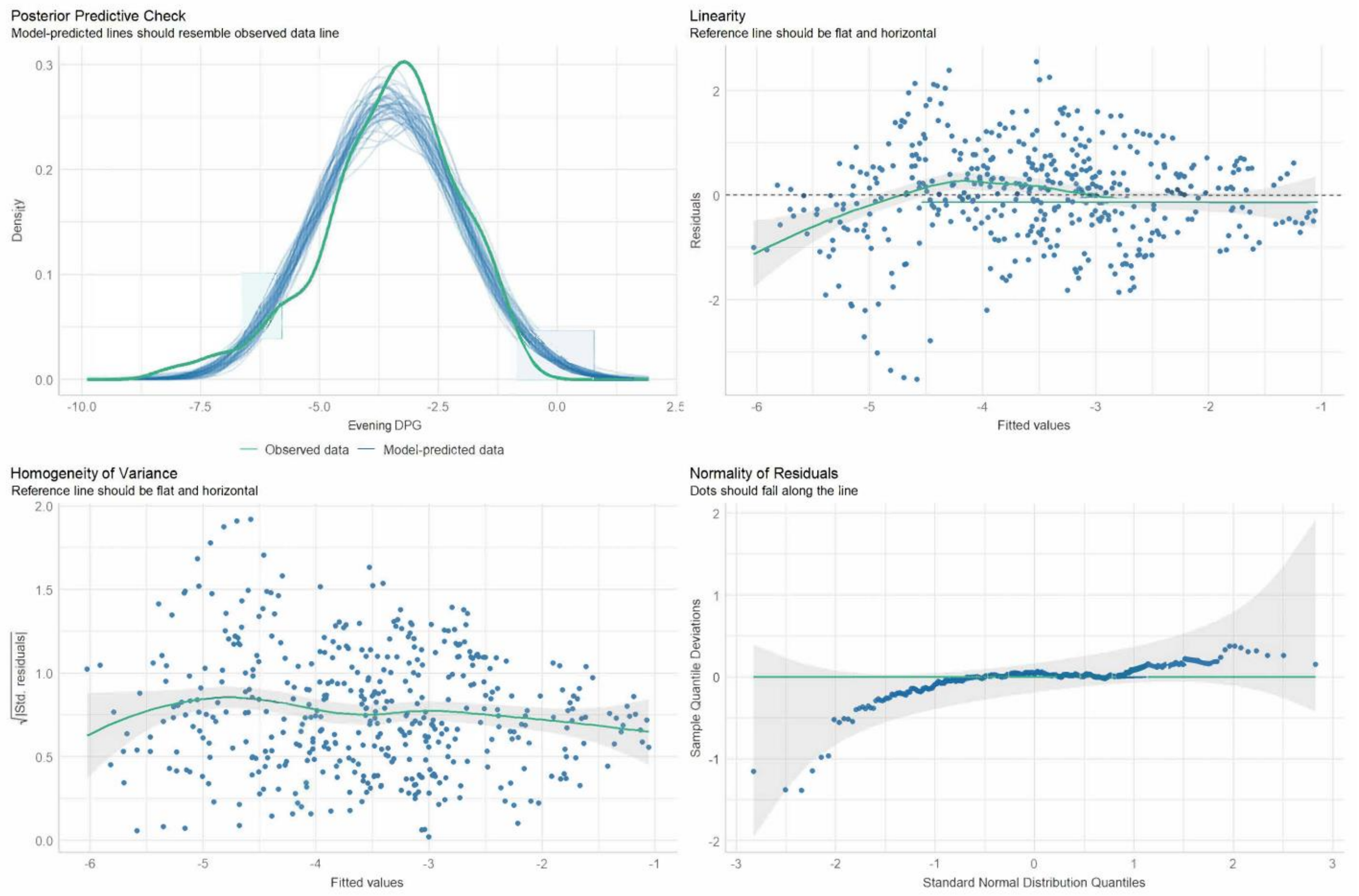


**Figure S16. Linear mixed model assumption checks for evening DPG.** Four different assumption checks were visualised for inspection: 1) Posterior predictive check comparing the density of predicted later evening DPG with observed data. 2) Linearity check, plotting residuals as a function of fitted values. 3) Homogeneity of variance (homoscedasticity) check, plotting the square root of standardized residuals as a function of fitted values. 4) Normality of Residuals check. In case of slight violations of these assumptions, the analyses were still performed due to the robustness of LMMs to deviations in distribution assumptions[3]. Abbreviations: DPG = distal-to-proximal skin temperature gradient, Std. residuals = Standardized residuals.

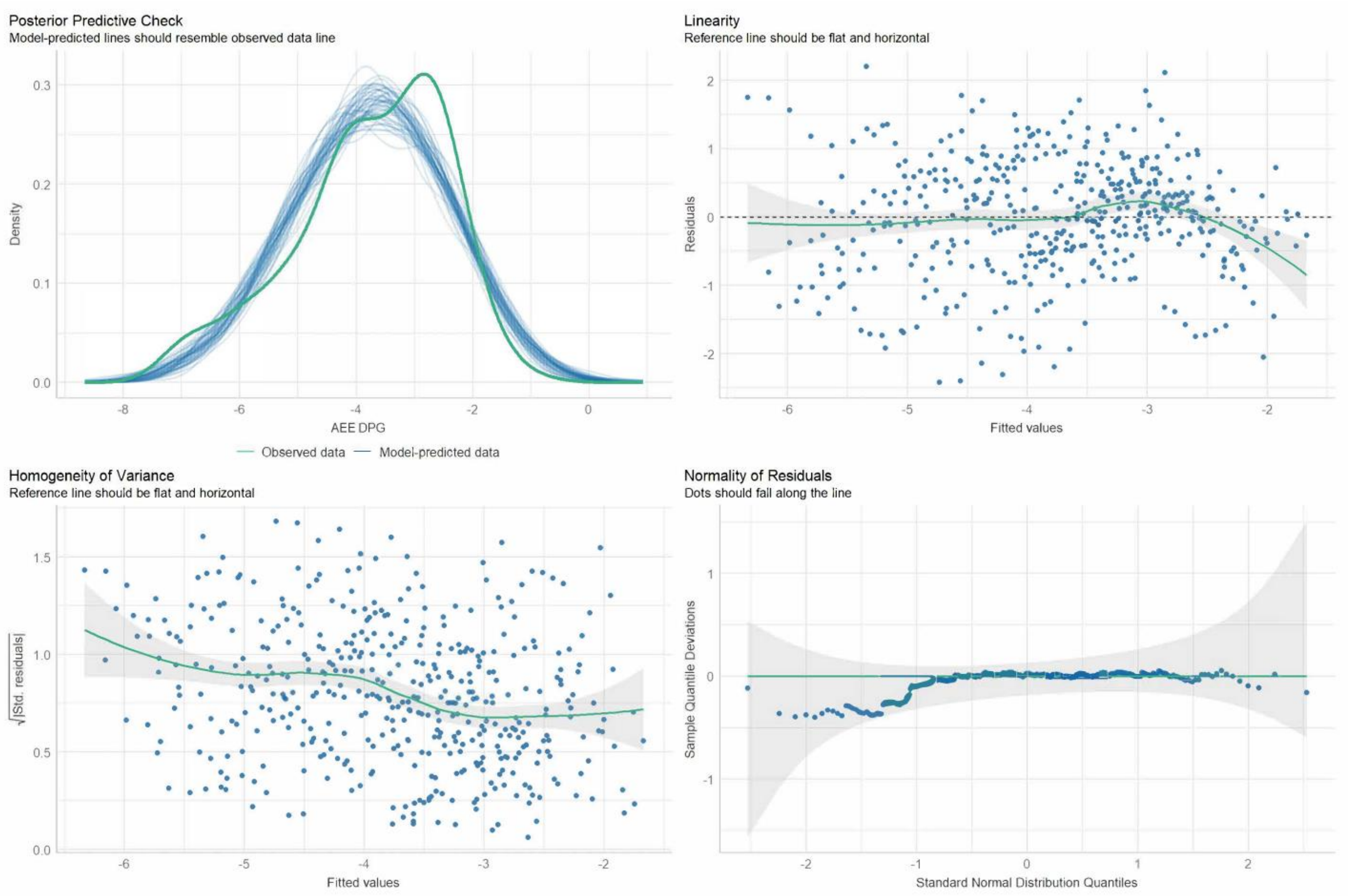


**Figure S17. Linear mixed model assumption checks for DPG during the AEE interventions.** Four different assumption checks were visualised for inspection: 1) Posterior predictive check comparing the density of predicted DPG during the AEE light interventions with observed data. 2) Linearity check, plotting residuals as a function of fitted values. 3) Homogeneity of variance (homoscedasticity) check, plotting the square root of standardized residuals as a function of fitted values. 4) Normality of Residuals check. In case of slight violations of these assumptions, the analyses were still performed due to the robustness of LMMs to deviations in distribution assumptions[3]. Abbreviations: AEE = afternoon to early evening, DPG = distal-to-proximal skin temperature gradient, Std. residuals = Standardized residuals.

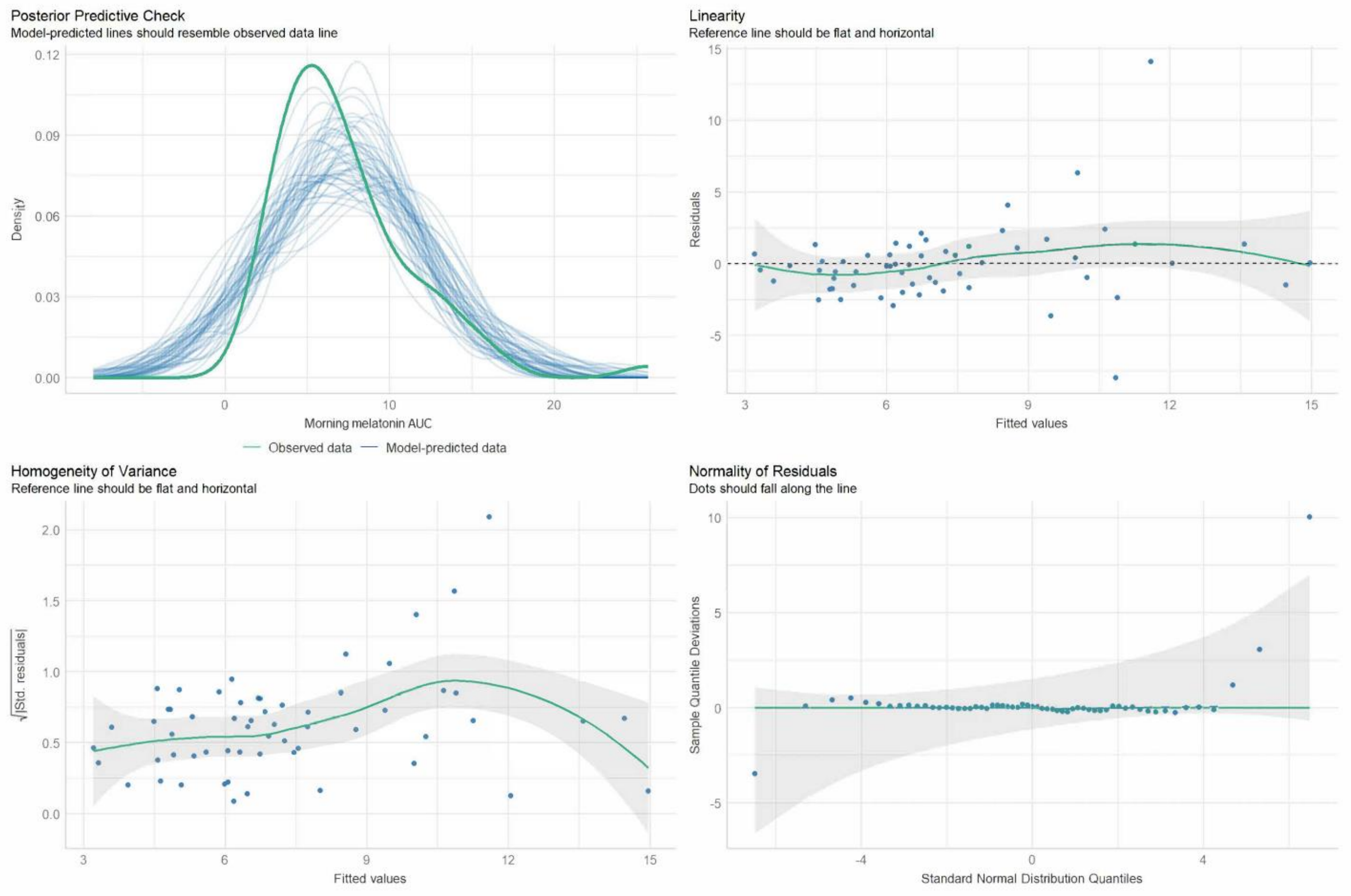


**Figure S18. Linear mixed model assumption checks for morning melatonin AUC.** Four different assumption checks were visualised for inspection: 1) Posterior predictive check comparing the density of predicted morning melatonin AUC data with observed data. 2) Linearity check, plotting residuals as a function of fitted values. 3) Homogeneity of variance (homoscedasticity) check, plotting the square root of standardized residuals as a function of fitted values. 4) Normality of Residuals check. In case of slight violations of these assumptions, the analyses were still performed due to the robustness of LMMs to deviations in distribution assumptions[3]. Abbreviations: AUC = Area under the curve, Std. residuals = Standardized residuals.

### Supplementary Tables

#### Table S1: Multi-stage screening and exclusion criteria.

| Aspect | Assessment modality | Exclusion criterion and cut-off |
|---|---|---|
| **Screening survey** | | |
| Previous enrolment into the current study | Self-report | Previous participation in the study |
| Age | Self-report | <14 years<br>>17 years |
| Pregnancy or breastfeeding (only female) | Participants are informed before study | 'Yes' response |
| Pubertal Stage | Self-administered rating scale for pubertal development[4] | Indication of pre-pubertal stage |
| BMI | Self-reported height and weight | BMI-PC <P3 or BMI-PC >P97 (Age-specific WHO curves) |
| Chronotype | Munich Chronotype Questionnaire (MCTQ)[5] | Extreme chronotype $MSF_{SC}$ < 1:00 / $MSF_{SC}$ > 7:00 |
| Sleep duration | Munich Chronotype Questionnaire (MCTQ)[5] | Extremely short or long sleep durations during schooldays < 6 hours<br>>11 hours |
| Sleep quality | Pittsburgh Sleep Quality Index, PSQI [6] | > 5 |
| Mental health & physical well-being | KIDSCREEN-27 index child & adolescent version[7] (Swiss Version)<br>Scales: Physical well-being & Psychological well-being | T values: T <35 (Swiss Norm, all genders, 12–18-year-olds) |
| Smoking | Self-reported frequency per day | >0 |
| High myopia | Self-report from prescription information | Worse than -6 diopters |
| High hyperopia | Self-report from prescription information | Worse than +6 diopters |
| Transmeridian travel (>2 zones) <1 month prior to the first session | Self-report | 'Yes' response |
| Shift work <3 months prior to study | Self-report | 'Yes' response |
| Current participation in other clinical trials | Self-report | 'Yes' response |
| Any ophthalmological or optometric conditions | Self-report in the general health questionnaire | Any 'Yes' response |
| Any general health concerns or disorders | Self-report in the general health questionnaire | Any 'Yes' response |
| Any debilitating allergies or allergies requiring medication | Self-report in the general health questionnaire | Any 'Yes' response |
| Any chronic medication | Self-report in the general health questionnaire | Any 'Yes' response |
| **In-person screening session** | | |
| Normal colour vision | Cambridge Colour Test, CCT[8,9] | |
| Normal best-corrected visual acuity (BCVA) | Landolt C test [10] | BCVA <0.5 (20/40) |
| Detectable Pupillary Light Response | Melanopsin-Sensitivity with Silent Substitution Pupil estimation[11] | Pupil not reliably detectable |
| General health & pubertal stage | Physical, and psychological examination | Study physician’s judgement |
| BMI | Measured height and weight | BMI-PC <P3 or BMI-PC >P97 (Age-specific WHO curves CH) |
| Ability to understand study language | In-person interaction with the experimenter | Experimenter judgment |

| Every study session | | |
|---|---|---|
| Drug use (AMP, BZD, COC, MOR/OPI, MTD and THC) | Drug-Screen-Multi 6; nal van Minden | Any positive test |
| Alcohol use | Alcohol test; (ACE X breathalyser) | Positive test (>0.1 ‰) |
| Sleep-wake times in 5 days prior to each experimental session | Actimetry record and sleep diary | >1 deviation from ±60-min window sleep and wake-up time |
| Caffeine abstinence in 5 days prior to each experimental session | Daily logs of caffeine intake | >1 times caffeine intake |
| Ability to follow study instructions | In-person interaction with the experimenter | Experimenter judgment |

The exclusion criteria were elaborated in detail on the informed consent sheet and discussed during an initial phone interview. Subsequently, a self-report screening survey (1st section in yellow), and an in-person screening session (2nd section in green) with the study physician (author C.E.) were conducted. Further criteria were monitored throughout the study (3rd section in blue), with tests conducted at every study session after arrival in the laboratory. In the table, each exclusion criterion is listed along with the measured aspect, the assessment modality, the timing during the screening process and (if applicable) the respective cut-off value.

**Table S2: LMM analysis for melatonin AUC and melatonin onset (sparse model).**

| Predictors | Estimates | | CI (95%) | | Std. error | | df | | t value | | *p* | |
|---|---|---|---|---|---|---|---|---|---|---|---|---|
| | Melatonin AUC | Melatonin onset time | Melatonin AUC | Melatonin onset time | Melatonin AUC | Melatonin onset time | Melatonin AUC | Melatonin onset time | Melatonin AUC | Melatonin onset time | Melatonin AUC | Melatonin onset time |
| Intercept | 83.17 | 20.68 | 53.07, 113.7 | 20.10 – 21.26 | 15.05 | 0.29 | 20 | 24 | - | - | - | - |
| Condition [moderate] | -7.81 | 0.31 | -17.69, 2.07 | -0.03 – 0.65 | 5.05 | 0.17 | 38 | 38 | -1.55 | 1.81 | 0.130 | 0.078 |
| Condition [bright] | -13.09 | 0.40 | -22.97, -3.21 | 0.06 – 0.74 | 5.05 | 0.17 | 38 | 38 | -2.59 | 2.31 | **0.013** | **0.026** |

| Random effects & Model fit criteria | $\sigma^2$ | $\tau_{00\ subjects}$ | ICC | $N_{subjects}$ | Observa-tions | Marginal $R^2$ / Conditional $R^2$ | AIC |
|---|---|---|---|---|---|---|---|
| Evening melatonin AUC | 254.88 | 4277.53 | 0.94 | 20 | 60 | 0.006 / 0.944 | 571.405 |
| Melatonin onset time | 0.30 | 1.40 | 0.82 | 20 | 60 | 0.017 / 0.827 | 163.646 |

Summary report of linear mixed model (LMM) analyses for melatonin AUC (primary outcome) and melatonin onset (post-hoc analysis), applying the sparse model. Estimates of the AEE light intervention conditions are given relative to the dim light condition. P-Values of ($p<0.05$) were considered as significant. Abbreviations: AIC = Akaike Information Criterion, AUC = Area under the curve, CI = Confidence interval (95%), ICC = Intraclass correlation coefficient, $N_{subjects}$ = Number of subjects in the analysis, $\sigma^2$ = Variance of the residuals of the model, $\tau_{00\ subjects}$ = Variance in the random intercepts at the subject level, Std. error = standard error of the estimates.

**Table S3: LMM analysis for morning melatonin AUC (covariate and sparse model).**

| Predictors | Estimates | | CI (95%) | | Std. error | | df | | t value | | *p* | |
|---|---|---|---|---|---|---|---|---|---|---|---|---|
| | Covariate model | Sparse model | Covariate model | Sparse model | Covariate model | Sparse model | Covariate model | Sparse model | Covariate model | Sparse model | Covariate model | Sparse model |
| Intercept | 10.46 | 7.41 | 3.31, 17.73 | 5.54, 9.28 | 4.14 | 0.96 | 15 | 40 | - | - | - | - |
| Condition [moderate] | -0.92 | -0.90 | -2.91, 1.07 | -2.87, 1.07 | 1.02 | 1.01 | 36 | 37 | 2.53 | -0.90 | 0.372 | 0.375 |
| Condition [bright] | 0.52 | 0.56 | -1.49, 2.56 | -1.43, 2.57 | 1.04 | 1.02 | 37 | 37 | -0.90 | 0.55 | 0.620 | 0.584 |
| Bright light history [TAT 1000 lx] | -0.22 | - | -0.74, 0.25 | - | 0.27 | - | 14 | - | -0.81 | - | 0.419 | - |
| *Chronotype [MSFsc] | -0.76 | - | -2.35, 0.83 | - | 0.92 | - | 15 | - | -0.83 | - | 0.641 | - |
| *Pubertal stage [late pubertal] | 0.82 | - | -2.24, 3.85 | - | 1.75 | - | 14 | - | 0.47 | - | 0.538 | - |
| *Pubertal Stage [midpubertal] | -1.22 | - | -4.63, 2.17 | - | 1.96 | - | 14 | - | -0.62 | - | 0.538 | - |
| *Pubertal Stage [early pubertal] | 6.57 | - | -0.44, 12.62 | - | 3.51 | - | 27 | - | 1.87 | - | 0.067 | - |
| **Random effects & Model fit criteria** | | | **$\sigma^2$** | **$\tau_{00}$ subjects** | **ICC** | **N subjects** | **Observa-tions** | **Marginal $R^2$ / Conditional $R^2$** | | **AIC** | | |
| Covariate model | | | 10.37 | 5.92 | 0.36 | 20 | 59 | 0.214 / 0.500 | | 321.833 | | |
| Sparse model | | | 10.11 | 8.22 | 0.45 | 20 | 59 | 0.020 / 0.459 | | 330.609 | | |

Summary report of linear mixed model (LMM) analyses for morning melatonin AUC (post-hoc analysis), applying the model with theoretically relevant covariates (covariate model) and the sparse model. Estimates of the AEE light intervention conditions are given relative to the dim light condition. Estimates of pubertal stage are given relative to “post pubertal stage”. P-Values of ($p<0.05$) were considered as significant. The covariates marked with an asterisk (*) were used as passive stratification factors to control for additional variance. Accordingly, these LMM estimates were not interpreted to avoid a "table 2 fallacy" [12]. Abbreviations: AEE = afternoon to early evening, AIC = Akaike Information Criterion, AUC = Area under the curve, CI = Confidence interval (95%), ICC = Intraclass correlation coefficient, $MSF_{SC}$ = Midpoint of sleep on free days corrected for oversleep, $N_{subjects}$ = Number of subjects in the analysis, $\sigma^2$ = Variance of the residuals of the model, $\tau_{00\ subjects}$ = Variance in the random intercepts at the subject level, Std. error = Standard error of the estimates, TAT 1000 lx = Time above threshold (>1000 lx illuminance, wrist-recorded).

**Table S4: LMM analysis for subjective sleepiness in the evening and during the AEE light intervention (covariate model).**

| Predictors | Estimates | | CI (95%) | | Std. error | | df | | t value | | *p* | |
|---|---|---|---|---|---|---|---|---|---|---|---|---|
| | Evening KSS | AEE KSS | Evening KSS | AEE KSS | Evening KSS | AEE KSS | Evening KSS | AEE KSS | Evening KSS | AEE KSS | Evening KSS | AEE KSS |
| Intercept | 6.50 | 5.90 | 3.44, 9.59 | 3.55, 8.21 | 1.77 | 1.35 | 12 | 13 | - | - | - | - |
| Condition [moderate] | 0.14 | -0.63 | -0.10, 0.39 | -0.90, -0.36 | 0.13 | 0.14 | 462 | 462 | 1.11 | -4.58 | 0.265 | **<0.001** |
| Condition [bright] | -0.13 | -0.92 | -0.38, 0.12 | -1.18, -0.64 | 0.13 | 0.14 | 463 | 465 | -1.00 | -6.63 | 0.320 | **<0.001** |
| Time (centred) | 0.73 | 0.29 | 0.59, 0.87 | 0.15, 0.44 | 0.07 | 0.08 | 461 | 462 | 10.24 | 3.90 | **<0.001** | **<0.001** |
| Condition [moderate] x Time (centred) | 0.01 | 0.08 | -0.19, 0.20 | -0.13, 0.28 | 0.10 | 0.11 | 461 | 462 | 0.05 | 0.71 | 0.958 | 0.477 |
| Condition [bright] x Time (centred) | -0.02 | -0.29 | -0.22, 0.17 | -0.49, -0.08 | 0.10 | 0.11 | 461 | 462 | -0.21 | -2.67 | 0.830 | **0.008** |
| Bright light history [TAT 1000 lx] | 0.24 | -0.04 | 0.13, 0.34 | -0.13, 0.07 | 0.05 | 0.05 | 245 | 113 | 4.57 | -0.70 | **<0.001** | 0.483 |
| *Chronotype [MSFsc] | -0.11 | -0.03 | -0.77, 0.55 | -0.52, 0.47 | 0.38 | 0.29 | 12 | 12 | -0.28 | -0.09 | 0.781 | 0.929 |
| *Pubertal stage [late pubertal] | -0.71 | -1.14 | -2.00, 0.56 | -2.10, -0.16 | 0.74 | 0.56 | 12 | 13 | -0.97 | -2.03 | 0.353 | 0.064 |
| *Pubertal Stage [midpubertal] | -1.03 | -0.87 | -2.44, 0.38 | -1.93, 0.20 | 0.82 | 0.62 | 12 | 12 | -1.26 | -1.40 | 0.232 | 0.185 |
| *Pubertal Stage [early pubertal] | -0.37 | -1.73 | -2.71, 1.94 | -3.48, 0.05 | 1.34 | 1.02 | 12 | 13 | -0.27 | -1.70 | 0.788 | 0.114 |
| **Random effects & Model fit criteria** | | | **$\sigma^2$** | **$\tau_{00\ SubjectID}$** | **ICC** | **$N_{SubjectID}$** | **Observa-tions** | | **Marginal $R^2$ / Conditional $R^2$** | | **AIC** | |

| | | | | | | | |
|---|---|---|---|---|---|---|---|
| Evening KSS | 1.30 | 1.24 | 0.49 | 18 | 486 | 0.346 / 0.665 | 1597.846 |
| AEE KSS | 1.54 | 0.68 | 0.31 | 18 | 486 | 0.182 / 0.432 | 1667.075 |

Summary report of linear mixed model (LMM) analyses of subjective sleepiness (KSS) in the later evening and during the AEE light intervention, applying the model with theoretically relevant covariates (covariate model). Estimates of the AEE light intervention conditions are given relative to the dim light condition. Estimates of pubertal stage are given relative to “post pubertal stage”. The “time” variable was median centred for each tested time period separately, so that “Time=0” refers to the middle of each tested time period. P-Values of ($p<0.05$) were considered as significant. The covariates marked with an asterisk (*) were used as passive stratification factors to control for additional variance. Accordingly, these LMM estimates were not interpreted to avoid a "table 2 fallacy" [12]. Abbreviations: AEE = afternoon to early evening, AIC = Akaike Information Criterion, CI = Confidence interval (95%), ICC = Intraclass correlation coefficient, KSS = Karolinska Sleepiness Scale, $MSF_{SC}$ = Midpoint of sleep on free days corrected for oversleep, $N_{subjects}$ = Number of subjects in the analysis, $\sigma^2$ = Variance of the residuals of the model, $\tau_{00\ subjects}$ = Variance in the random intercepts at the subject level, Std. error = standard error of the estimates, TAT 1000 lx = Time above threshold (>1000 lx illuminance, wrist-recorded).

**Table S5: LMM analysis for subjective sleepiness in the evening and during the AEE light intervention (sparse model).**

| Predictors | Estimates | | CI (95%) | | Std. error | | df | | t value | | *p* | |
|---|---|---|---|---|---|---|---|---|---|---|---|---|
| | Evening KSS | AEE KSS | Evening KSS | AEE KSS | Evening KSS | AEE KSS | Evening KSS | AEE KSS | Evening KSS | AEE KSS | Evening KSS | AEE KSS |
| Intercept | 6.00 | 4.86 | 5.51, 6.49 | 4.43, 5.29 | 0.22 | 0.24 | 21 | 23 | - | - | - | - |
| Condition [moderate] | 0.17 | -0.64 | -0.08, 0.43 | -0.90, -0.37 | 0.13 | 0.14 | 463 | 463 | 1.33 | -4.61 | 0.184 | **<0.001** |
| Condition [bright] | -0.19 | -0.91 | -0.45, 0.06 | -1.18, -0.64 | 0.13 | 0.14 | 463 | 463 | -1.48 | -6.59 | 0.140 | **<0.001** |
| Time (centred) | 0.73 | 0.29 | 0.59, 0.87 | 0.15, 0.44 | 0.07 | 0.08 | 463 | 463 | 10.00 | 3.90 | **<0.001** | **<0.001** |
| Condition [moderate] x Time (centred) | 0.01 | 0.08 | -0.20, 0.21 | -0.13, 0.28 | 0.10 | 0.11 | 463 | 463 | 0.05 | 0.71 | 0.959 | 0.477 |
| Condition [bright] x Time (centred) | -0.02 | -0.29 | -0.22, 0.18 | -0.49, -0.08 | 0.10 | 0.11 | 463 | 463 | -0.21 | -2.67 | 0.834 | **0.008** |

| Random effects & Model fit criteria | $\sigma^2$ | $\tau_{00\ SubjectID}$ | ICC | $N_{SubjectID}$ | Observations | Marginal $R^2$ / Conditional $R^2$ | AIC |
|---|---|---|---|---|---|---|---|
| Evening KSS | 1.37 | 0.92 | 0.40 | 18 | 486 | 0.272 / 0.566 | 1610.130 |
| AEE KSS | 1.54 | 0.69 | 0.31 | 18 | 486 | 0.108 / 0.385 | 1660.703 |

Summary report of linear mixed model (LMM) analyses of subjective sleepiness (KSS) in the later evening and during the AEE light intervention, applying the sparse model. Estimates of the AEE light intervention conditions are given relative to the dim light condition. The "Time" variable was median centred for each tested time period separately, so that "Time=0" refers to the middle of each tested time period. P-Values of ($p<0.05$) were considered as significant. Abbreviations: AEE = afternoon to early evening, AIC = Akaike Information Criterion, CI = Confidence interval (95%), ICC = Intraclass correlation coefficient, KSS = Karolinska Sleepiness Scale, $N_{subjects}$ = Number of Subjects in the analysis, $\sigma^2$ = Variance of the residuals of the model, $\tau_{00\ subjects}$ = Variance in the random intercepts at the subject level, std. Error = standard error of the estimates.

**Table S6: LMM analysis (covariate model) for PVT response speed in the evening and during the AEE light intervention.**

| Predictors | Estimates | | CI (95%) | | Std. error | | df | | t value | | *p* | |
|---|---|---|---|---|---|---|---|---|---|---|---|---|
| | Evening PVT | AEE PVT | Evening PVT | AEE PVT | Evening PVT | AEE PVT | Evening PVT | AEE PVT | Evening PVT | AEE PVT | Evening PVT | AEE PVT |
| Intercept | 4.88 | 5.11 | 3.49, 6.27 | 3.97, 6.25 | 0.77 | 0.64 | 17 | 17 | - | - | - | - |
| Condition [moderate] | -0.06 | -0.10 | -0.17, 0.05 | -0.23, 0.04 | 0.06 | 0.07 | 170 | 170 | -1.06 | -1.40 | 0.290 | 0.163 |
| Condition [bright] | -0.04 | -0.09 | -0.15, 0.07 | -0.23, 0.04 | 0.06 | 0.07 | 171 | 172 | -0.74 | -1.36 | 0.461 | 0.176 |
| Time (centred) | -0.07 | 0.02 | -0.15, 0.00 | -0.06, 0.10 | 0.04 | 0.04 | 170 | 170 | -1.82 | 0.49 | 0.070 | 0.622 |
| Condition [moderate] x Time (centred) | -0.01 | 0.01 | -0.12, 0.10 | -0.10, 0.12 | 0.05 | 0.06 | 170 | 170 | -0.19 | 0.12 | 0.846 | 0.907 |
| Condition [bright] x Time (centred) | -0.05 | -0.03 | -0.16, 0.05 | -0.14, 0.08 | 0.05 | 0.06 | 170 | 170 | -0.95 | -0.50 | 0.341 | 0.622 |
| Bright light history [TAT 1000 lx] | -0.04 | -0.04 | -0.09, 0.01 | -0.09, 0.02 | 0.03 | 0.03 | 180 | 133 | -1.54 | -1.29 | 0.126 | 0.201 |
| *Chronotype [MSFsc] | -0.02 | -0.03 | -0.33, 0.30 | -0.29, 0.22 | 0.18 | 0.14 | 17 | 17 | -0.09 | -0.24 | 0.926 | 0.816 |
| *Pubertal stage [late pubertal] | 0.31 | 0.39 | -0.32, 0.95 | -0.13, 0.91 | 0.35 | 0.29 | 17 | 17 | 0.89 | 1.33 | 0.376 | 0.199 |
| *Pubertal Stage [midpubertal] | 0.07 | 0.05 | - 0.66 – 0.80 | -0.55, 0.65 | 0.41 | 0.34 | 17 | 17 | 0.17 | 0.16 | 0.867 | 0.877 |
| *Pubertal Stage [early pubertal] | -1.28 | -1.34 | - 2.58 – 0.01 | -2.40, -0.27 | 0.72 | 0.59 | 17 | 17 | -1.78 | -2.25 | 0.077 | 0.038 |
| **Random effects & Model fit criteria** | | | **$\sigma^2$** | **$\tau_{00}$ SubjectID** | **ICC** | **N SubjectID** | **Observa-tions** | **Marginal $R^2$ / Conditional $R^2$** | | **AIC** | | - |

| | | | | | | | | | |
|---|---|---|---|---|---|---|---|---|---|
| Evening PVT response speed | 0.10 | 0.40 | 0.80 | 22 | 198 | 0.201 / 0.837 | 238.979 | | - |
| AEE PVT response speed | 0.16 | 0.26 | 0.63 | 22 | 198 | 0.256 / 0.722 | 303.077 | | |

Summary report of linear mixed model (LMM) analyses of the PVT response speed (mean 1/RT) in the later evening and during the AEE light intervention, applying the model with theoretically relevant covariates (covariate model). Estimates of the AEE light intervention conditions are given relative to the dim light condition. Estimates of pubertal stage are given relative to “post pubertal stage”. The “time” variable was median centred for each tested time period separately, so that “Time=0” refers to the middle of each tested time period. P-Values of ($p<0.05$) were considered as significant. The covariates marked with an asterisk (*) were used as passive stratification factors to control for additional variance. Accordingly, these LMM estimates were not interpreted to avoid a "table 2 fallacy" [12]. Abbreviations: AEE = afternoon to early evening, AIC = Akaike Information Criterion, CI = Confidence interval (95%), ICC = Intraclass correlation coefficient, $MSF_{SC}$ = Midpoint of sleep on free days corrected for oversleep, $N_{subjects}$ = Number of Subjects in the analysis, PVT = Psychomotor Vigilance Task, $\sigma^2$ = Variance of the residuals of the model, $\tau_{00\ subjects}$ = Variance in the random intercepts at the subject level, std. Error = standard error of the estimates, TAT 1000 lx = Time above threshold (>1000 lx illuminance, wrist-recorded).

**Table S7: LMM analysis (sparse model) for PVT response speed in the evening and during the AEE light intervention.**

| Predictors | Estimates | | CI (95%) | | Std. error | | df | | t value | | *p* | |
|---|---|---|---|---|---|---|---|---|---|---|---|---|
| | Evening PVT | AEE PVT | Evening PVT | AEE PVT | Evening PVT | AEE PVT | Evening PVT | AEE PVT | Evening PVT | AEE PVT | Evening PVT | AEE PVT |
| Intercept | 4.85 | 5.05 | 4.56, 5.15 | 4.78, 5.32 | 0.15 | 0.14 | 23 | 25 | - | - | - | - |
| Condition [moderate] | -0.06 | -0.10 | -0.17, 0.05 | -0.23, 0.04 | 0.06 | 0.07 | 171 | 171 | -1.03 | -1.38 | 0.303 | 0.169 |
| Condition [bright] | -0.03 | -0.08 | -0.14, 0.08 | -0.22, 0.05 | 0.06 | 0.07 | 171 | 171 | -0.54 | -1.21 | 0.589 | 0.226 |
| Time (centred) | -0.07 | 0.02 | -0.15, 0.00 | -0.06, 0.10 | 0.04 | 0.04 | 171 | 171 | -1.82 | 0.49 | 0.071 | 0.623 |
| Condition [moderate] x Time (centred) | -0.01 | 0.01 | -0.12, 0.10 | -0.10, 0.12 | 0.05 | 0.06 | 171 | 171 | -0.19 | 0.12 | 0.847 | 0.907 |
| Condition [bright] x Time (centred) | -0.05 | -0.03 | -0.16, 0.05 | -0.14, 0.08 | 0.05 | 0.06 | 171 | 171 | -0.95 | -0.50 | 0.343 | 0.617 |
| **Random effects & Model fit criteria** | | | **$\sigma^2$** | **$\tau_{00\ SubjectID}$** | **ICC** | **$N_{SubjectID}$** | **Observa-tions** | **Marginal $R^2$ / Conditional $R^2$** | | **AIC** | | - |
| Evening PVT response speed | | | 0.11 | 0.44 | 0.81 | 22 | 198 | 0.018 / 0.812 | | 230.391 | | - |
| AEE PVT response speed | | | 0.16 | 0.35 | 0.69 | 22 | 198 | 0.005 / 0.694 | | 296.263 | | |

Summary report of linear mixed model (LMM) analyses of the PVT response speed (mean 1/RT) in the later evening and during the AEE light intervention, applying the sparse model. Estimates of the AEE light intervention conditions are given relative to the dim light condition. The "Time" variable was median centred for each tested time period separately, so that "Time=0" refers to the middle of each tested time period. P-Values of ($p<0.05$) were considered as significant. Abbreviations: AEE = afternoon to early evening, AIC = Akaike Information Criterion, CI = Confidence interval (95%), ICC = Intraclass correlation coefficient, $N_{subjects}$ = Number of subjects in the analysis, PVT = Psychomotor Vigilance Task, $\sigma^2$ = Variance of the residuals of the model, $\tau_{00\ subjects}$ = Variance in the random intercepts at the subject level, std. Error = standard error of the estimates.

**Table S8: LMM analysis for the DPG in the evening and during the AEE light intervention (covariate model).**

| Predictors | Estimates | | CI (95%) | | Std. error | | df | | t value | | *p* | |
|---|---|---|---|---|---|---|---|---|---|---|---|---|
| | Evening DPG | AEE DPG | Evening DPG | AEE DPG | Evening DPG | AEE DPG | Evening DPG | AEE DPG | Evening DPG | AEE DPG | Evening DPG | AEE DPG |
| Intercept | -3.52 | -2.97 | -5.48, -1.56 | -5.06, -0.89 | 1.10 | 1.17 | 16 | 16 | - | - | - | - |
| Condition [moderate] | -0.41 | -0.27 | -0.63, -0.20 | -0.46, -0.07 | 0.11 | 0.10 | 415 | 414 | -3.71 | -2.67 | **<0.001** | **0.008** |
| Condition [bright] | 0.49 | 0.69 | 0.27, 0.71 | 0.50, 0.89 | 0.11 | 0.10 | 417 | 416 | 4.36 | 6.90 | **<0.001** | **<0.001** |
| Time (centred) | 0.07 | -0.26 | -0.09, 0.22 | -0.40, -0.12 | 0.08 | 0.07 | 414 | 414 | 0.86 | -3.69 | 0.388 | **<0.001** |
| Condition [moderate] x Time (centred) | 0.17 | -0.09 | -0.05, 0.38 | -0.29, 0.10 | 0.11 | 0.10 | 414 | 414 | 1.49 | -0.92 | 0.136 | 0.357 |
| Condition [bright] x Time (centred) | 0.05 | 0.09 | -0.17, 0.27 | -0.11, 0.28 | 0.11 | 0.10 | 414 | 414 | 0.43 | 0.89 | 0.665 | 0.374 |
| Bright light history [TAT 1000 lx] | 0.00 | -0.02 | -0.09, 0.10 | -0.10, 0.07 | 0.05 | 0.04 | 202 | 273 | 0.01 | -0.45 | 0.994 | 0.652 |
| *Chronotype [MSFsc] | -0.11 | -0.17 | -0.55, 0.33 | -0.64, 0.30 | 0.25 | 0.26 | 16 | 16 | -0.45 | -0.65 | 0.661 | 0.526 |
| *Pubertal stage [late pubertal] | 0.75 | -0.52 | -0.16, 1.67 | -1.49, 0.45 | 0.51 | 0.54 | 16 | 16 | 1.47 | -0.96 | 0.161 | 0.352 |
| *Pubertal Stage [midpubertal] | -0.31 | -0.08 | -1.34, 0.72 | -1.18, 1.02 | 0.58 | 0.62 | 16 | 16 | -0.54 | -0.13 | 0.598 | 0.895 |
| *Pubertal Stage [early pubertal] | 2.15 | 0.65 | 0.32, 3.98 | -1.29, 2.60 | 1.03 | 1.09 | 16 | 16 | 2.09 | 0.59 | 0.053 | 0.561 |
| **Random effects & Model fit criteria** | | | **$\sigma^2$** | **$\tau_{00\ SubjectID}$** | **ICC** | **$N_{SubjectID}$** | **Observa-tions** | **Marginal $R^2$ / Conditional $R^2$** | | **AIC** | | - |

| | | | | | | | | | |
|---|---|---|---|---|---|---|---|---|---|
| Evening DPG | 0.91 | 0.79 | 0.47 | 21 | 441 | 0.246 / 0.597 | 1306.459 | | - |
| AEE DPG | 0.73 | 0.92 | 0.56 | 21 | 441 | 0.176 / 0.635 | 1216.590 | | - |

Summary report of linear mixed model (LMM) analyses of the DPG in the later evening and during the AEE light intervention, applying the model with theoretically relevant covariates (covariate model). Estimates of the AEE light intervention conditions are given relative to the dim light condition. Estimates of pubertal stage are given relative to “post pubertal stage”. The “Time” variable was median centred for each tested time period separately, so that “Time=0” refers to the middle of each tested time period. P-Values of ($p<0.05$) were considered as significant. The covariates marked with an asterisk (*) were used as passive stratification factors to control for additional variance. Accordingly, these LMM estimates were not interpreted to avoid a "table 2 fallacy" [12]. Abbreviations: AEE = afternoon to early evening, AIC = Akaike Information Criterion, CI = Confidence interval (95%), DPG = distal-to-proximal skin temperature gradient, ICC = Intraclass correlation coefficient, $MSF_{SC}$ = Midpoint of sleep on free days corrected for oversleep, $N_{subjects}$ = Number of Subjects in the analysis, $\sigma^2$ = Variance of the residuals of the model, $\tau_{00\ subjects}$ = Variance in the random intercepts at the subject level, std. Error = standard error of the estimates, TAT 1000 lx = Time above threshold (>1000 lx illuminance, wrist-recorded).

**Table S9: LMM analysis (sparse model) for the DPG in the evening and during the AEE light intervention.**

| Predictors | Estimates | | CI (95%) | | Std. error | | df | | t value | | *p* | |
|---|---|---|---|---|---|---|---|---|---|---|---|---|
| | Evening DPG | AEE DPG | Evening DPG | AEE DPG | Evening DPG | AEE DPG | Evening DPG | AEE DPG | Evening DPG | AEE DPG | Evening DPG | AEE DPG |
| Intercept | -3.56 | -3.91 | -4.03, -3.09 | -4.33, -3.49 | 0.24 | 0.21 | 23 | 23 | - | - | - | - |
| Condition [moderate] | -0.41 | -0.26 | -0.63, -0.20 | -0.46, -0.07 | 0.11 | 0.10 | 415 | 415 | -3.72 | -2.65 | **<0.001** | **0.008** |
| Condition [bright] | 0.49 | 0.70 | 0.27, 0.71 | 0.50, 0.89 | 0.11 | 0.10 | 415 | 415 | 4.40 | 7.01 | **<0.001** | **<0.001** |
| Time (centred) | 0.07 | -0.26 | -0.09, 0.22 | -0.40, -0.12 | 0.08 | 0.07 | 415 | 415 | 0.87 | -3.70 | 0.387 | **<0.001** |
| Condition [moderate] x Time (centred) | 0.17 | -0.09 | -0.05, 0.38 | -0.29, 0.10 | 0.11 | 0.10 | 415 | 415 | 1.50 | -0.92 | 0.135 | 0.357 |
| Condition [bright] x Time (centred) | 0.05 | 0.09 | -0.17, 0.27 | -0.11, 0.28 | 0.11 | 0.10 | 415 | 415 | 0.43 | 0.89 | 0.665 | 0.374 |
| **Random effects & Model fit criteria** | | | **$\sigma^2$** | **$\tau_{00\ SubjectID}$** | **ICC** | **$N_{SubjectID}$** | **Observa-tions** | **Marginal $R^2$ / Conditional $R^2$** | | **AIC** | | - |
| Evening DPG | | | 0.91 | 1.03 | 0.53 | 21 | 441 | 0.077 / 0.568 | | 1303.002 | | - |
| AEE DPG | | | 0.73 | 0.82 | 0.53 | 21 | 441 | 0.134 / 0.592 | | 1206.811 | | - |

Summary report of linear mixed model (LMM) analyses of the DPG in the later evening and during the AEE light intervention, applying the sparse model. Estimates of the AEE light intervention conditions are given relative to the dim light condition. The “Time” variable was median centred for each tested time period separately, so that “Time=0” refers to the middle of each tested time period. P-Values of ($p<0.05$) were considered as significant. Abbreviations: AEE = afternoon to early evening, AIC = Akaike Information Criterion, CI = Confidence interval (95%), DPG = distal-to-proximal skin temperature gradient, ICC = Intraclass correlation coefficient, $N_{subjects}$ = Number of subjects in the analysis, $\sigma^2$ = Variance of the residuals of the model, $\tau_{00\ subjects}$ = Variance in the random intercepts at the subject level, std. Error = standard error of the estimates.

**Table S10: Specifications of light sources outside the laboratory without and with filter glasses.**

| Light Scene | Brightest corridor spot | | Brightest bathroom spot | |
|---|---|---|---|---|
| Filter status | no filter | filter lenses | no filter | filter lenses |
| Illuminance (lx) | 14.50 | 5.57 | 11.90 | 4.57 |
| Melanopic EDI (lx) | 6.65 | 0.20 | 5.33 | 0.16 |
| S-cone-opic EDI (lx) | 5.65 | 0.00 | 5.28 | 0.00 |
| M-cone-opic EDI (lx) | 12.12 | 2.75 | 9.87 | 2.22 |
| L-cone-opic EDI (lx) | 14.40 | 6.25 | 11.82 | 5.15 |
| Rhodopic EDI (lx) | 8.17 | 0.56 | 6.59 | 0.45 |
| Melanopic irradiance ($mW \cdot m^{-2}$) | 8.82 | 0.26 | 7.06 | 0.22 |
| S-cone-opic irradiance ($mW \cdot m^{-2}$) | 4.62 | 0.00 | 4.31 | 0.00 |
| M-cone-opic irradiance ($mW \cdot m^{-2}$) | 17.64 | 4.00 | 14.36 | 3.23 |
| L-cone-opic irradiance ($mW \cdot m^{-2}$) | 23.45 | 10.19 | 19.25 | 8.38 |
| Rhodopic irradiance ($mW \cdot m^{-2}$) | 11.84 | 0.81 | 9.55 | 0.66 |
| CIE 1964 $x_{10}y_{10}$ chromaticity [$x_{10}$] | 0.42 | 0.63 | 0.42 | 0.63 |
| CIE 1964 $x_{10}y_{10}$ chromaticity [$y_{10}$] | 0.40 | 0.37 | 0.39 | 0.37 |
| CCT (K) - Ohno, 2013 | 3450 | 1233 | 3429 | 1208 |
| Colour Rendering Index [Ra] | 81.13 | 40.75 | 80.63 | 43.25 |

During the experiment, participants were only exposed to these light conditions during their toilet/walking breaks and on their way to the pupillograph in the neighboring laboratory for the pupillary light response measurements. They wore red-orange tinted short-wavelength filter goggles at all times. Values used for stimulus specification include illuminance, alpha-opic equivalent daylight illuminances (alpha-opic EDIs), CIE 1931 xy chromaticity coordinates, correlated colour temperature (CCT), and colour rendering index (CRI). The unfiltered values were derived from spectral irradiance measurements taken in the vertical plane at the approximate eye level of a standing participant (160 cm height). Values with filter were derived from spectral irradiances calculated by multiplying the prior measurements with the filter transmittance data (see Suppl. Figure S6).

**Table S11: Stimulus specifications of the pupillary light response measurements.**

| Pupillograph setting | Light flux | | Melanopsin | |
|---|---|---|---|---|
| | Background | Modulation | Background | Modulation |
| Duration (sec) | 474 | 30 | 474 | 30 |
| Illuminance (lx) | 201.60 | 784.80 | 657.97 | 643.87 |
| Melanopic EDI (lx) | 75.00 | 296.78 | 7.84 | 38.81 |
| S-cone-opic EDI (lx) | 36.22 | 147.14 | 5.24 | 6.85 |
| M-cone-opic EDI (lx) | 116.28 | 455.67 | 265.48 | 260.84 |
| L-cone-opic EDI (lx) | 228.65 | 891.04 | 766.38 | 753.30 |
| Rhodopic EDI (lx) | 80.30 | 310.47 | 33.77 | 61.56 |
| Melanopic EDI (lx) | 75.00 | 296.78 | 7.84 | 38.81 |
| CIE 1964 $x_{10}y_{10}$ chromaticity [$x_{10}$] | 0.57 | 0.57 | 0.65 | 0.65 |
| CIE 1964 $x_{10}y_{10}$ chromaticity [$y_{10}$] | 0.36 | 0.36 | 0.35 | 0.35 |
| CCT (K) - Ohno, 2013 | 1324.01 | 1336.72 | N/A | N/A |
| Colour Rendering Index [Ra] | 53.00 | 57.88 | 23.25 | 48.50 |

Pupillary light response measurements were taken at baseline (7 h 50 min before HBT) and again 45 min before the end of the light interventions (3 h 45 min before HBT). Participants were dark adapted (<0.5 lx) for 3 min before the pupil measurements. Values used for stimulus specification include the total duration in seconds of each pupil measurements setting (“Lightflux Background”, “Lightflux Modulation”, “Melanopsin Background” and “Melanopsin Modulation”) and the respective illuminance, alpha-opic equivalent daylight illuminances (alpha-opic EDIs), CIE 1931 xy chromaticity coordinates, correlated colour temperature (CCT), and colour rendering index (CRI).

**Table S12: Detailed stimulus specification of the light interventions in radiance.**

| | Light intervention condition | | | | | | | | | | | |
|---|---|---|---|---|---|---|---|---|---|---|---|---|
| **Condition** | **Dim** | | | | **Moderate** | | | | **Bright** | | | |
| Object surface | curtain | wall | wall | table | curtain | wall | wall | table | curtain | wall | wall | table |
| Sensor orientation | forward | 90° left | 90° right | 60° down | forward | 90° left | 90° right | 60° down | forward | 90° left | 90° right | 60° down |
| Measurement plane | vertical | vertical | vertical | horizontal | vertical | vertical | vertical | horizontal | vertical | vertical | vertical | horizontal |
| Luminance (cd/m²) | 1.57 | 0.94 | 0.97 | 4.55 | 33.44 | 23.64 | 23.93 | 58.61 | 589.52 | 436.24 | 442.58 | 1081.03 |
| Melanopic EDL (cd/m²) | 0.90 | 0.52 | 0.54 | 2.62 | 21.16 | 14.67 | 14.84 | 37.59 | 328.20 | 236.66 | 239.88 | 606.36 |
| S-cone-opic EDL (cd/m²) | 1.08 | 0.60 | 0.62 | 3.00 | 24.24 | 16.42 | 16.63 | 43.10 | 389.06 | 271.87 | 275.46 | 717.86 |
| M-cone-opic EDL (cd/m²) | 1.36 | 0.81 | 0.84 | 3.96 | 29.15 | 20.57 | 20.83 | 51.37 | 512.22 | 377.45 | 383.04 | 941.99 |
| L-cone-opic EDL (cd/m²) | 1.56 | 0.93 | 0.96 | 4.52 | 33.58 | 23.71 | 24.00 | 58.78 | 583.99 | 431.92 | 438.10 | 1070.11 |
| Rhodopic EDL (cd/m²) | 1.02 | 0.60 | 0.62 | 2.97 | 23.16 | 16.17 | 16.37 | 41.06 | 377.70 | 274.74 | 278.67 | 697.08 |
| Melanopic radiance (mW · m⁻² · sr) | 1.20 | 0.69 | 0.71 | 3.48 | 28.06 | 19.45 | 19.68 | 49.85 | 435.26 | 313.86 | 318.14 | 804.16 |
| S-cone-opic radiance (mW · m⁻² · sr) | 0.88 | 0.49 | 0.51 | 2.45 | 19.81 | 13.42 | 13.59 | 35.22 | 317.97 | 222.19 | 225.13 | 586.70 |
| M-cone-opic radiance (mW · m⁻² · sr) | 1.98 | 1.18 | 1.22 | 5.76 | 42.44 | 29.95 | 30.32 | 74.78 | 745.70 | 549.50 | 557.64 | 1371.37 |
| L-cone-opic radiance (mW · m⁻² · sr) | 2.53 | 1.52 | 1.56 | 7.36 | 54.70 | 38.63 | 39.09 | 95.75 | 951.27 | 703.56 | 713.62 | 1743.11 |
| Rhodopic radiance (mW · m⁻² · sr) | 1.48 | 0.86 | 0.89 | 4.31 | 33.58 | 23.44 | 23.73 | 59.53 | 547.56 | 398.29 | 403.99 | 1010.57 |
| CIE 1964 $x_{10}y_{10}$ chromaticity [$x_{10}$] | 0.38 | 0.38 | 0.38 | 0.38 | 0.38 | 0.38 | 0.38 | 0.37 | 0.38 | 0.38 | 0.38 | 0.38 |
| CIE 1964 $x_{10}y_{10}$ chromaticity [$y_{10}$] | 0.35 | 0.36 | 0.36 | 0.36 | 0.35 | 0.35 | 0.35 | 0.35 | 0.36 | 0.36 | 0.36 | 0.36 |
| CCT (K) - Ohno, 2013 | 4119 | 4009 | 4015 | 4093 | 3997 | 3955 | 3960 | 4085 | 4108 | 4026 | 4031 | 4154 |
| Colour Rendering Index [Ra] | 80 | 79 | 79 | 81 | 87 | 87 | 87 | 87 | 79 | 79 | 79 | 79 |

Summary of the spectral radiance derived values for the 3 AEE light interventions on different object surfaces. The specific locations of these 4 reported radiance measurements are illustrated in Suppl. Fig. S8. The subsequent evening light condition had the same characteristics as the "moderate" light intervention. Participants were instructed and monitored to remain seated and keep their eyes open with their sitting posture directed forward. The spectral radiance measurements were taken from the observer's point of view with a research-grade spectroradiometer (Spectraval 1501, JETI Technische Instrumente GmbH, Jena, Germany). The sensor was placed at the eye level of the seated participants (115 cm from the floor, 85 cm from the overhead light source, 80 cm from the white curtain in front of them). Values used for stimulus specification were calculated using the luox application [2] and include luminance, alpha-opic equivalent daylight luminances (alpha-opic EDLs), alpha-opic radiances (given in mW · m⁻² · sr), CIE 1931 xy chromaticity coordinates, correlated colour temperature (CCT), and colour rendering index (CRI).

**Table S13: ENLIGHT Checklist.**

Note: This document is best viewed and completed with Adobe Acrobat Reader.

ENLIGHT

Release 1.0.2, 16 October 2023

### ENLIGHT Checklist

Below is the **ENLIGHT Checklist** for reporting ocular light exposures in human laboratory-based studies. We will strongly encourage that this checklist be used in conjunction with the **ENLIGHT Explanation & Elaboration (E&E) document**. This checklist is intended both to help authors, reviewers, and editors in evaluating the completeness of reporting in submitted studies, and for documentation of studies after publication. In the location column, please indicate the page, figure, or table number where the item or description can be found. If an item is not available, please select "**Not available**". If you consider an item not to be applicable in your specific study design after consulting the guidelines, please select "**Not applicable**". Items which do not have the option to select "Not applicable" were rated by experts as applicable for all studies, regardless of context. If you are unable to provide the information, please select "Not available".

The **ENLIGHT Checklist** (this document) and the **ENLIGHT E&E document** are released under the CC-BY-NC-ND License. For more information, please visit http://enlight-statement.org/.

#### General Information

**Author names:** R.Lazar, F. Fazlali, M. Dourte, C. Epple, O. Stefani, M.Spitschan, C. Cajochen

**Title of manuscript:** Afternoon to early evening bright light exposure reduces later melatonin production in adolescents

**Date:** 01 October 2024

#### A. Study Characteristics

##### A.1. Protocol-level characteristics

| | Location (page, figure, table number) | Not available | Not applicable |
|---|---|---|---|
| Description of experimental setting | Fig. 2, manuscript pages 11-13 | ☐ | |
| Timeline of experiment (including timing and duration of light) | Fig. 1, manuscript pages 11-13 | ☐ | |
| Pre-laboratory sleep-wake/rest-activity behaviour | Manuscript pages 11-13 | ☐ | ☐ |
| Pre-laboratory light exposure | Fig. 4, manuscript page 14 | ☐ | ☐ |
| Immediate prior light exposure (in laboratory) | Fig. 2, manuscript pages 11-13 | ☐ | ☐ |

##### A.2. Measurement-level characteristics

| | | | |
|---|---|---|---|
| Measurement plane (e.g., horizontal or vertical) | Fig. 2, Table 3, Suppl. Fig. S7, Suppl. Fig. S8, Suppl. Table S12 | ☐ | |
| Measurement viewpoint and location | Fig. 2, Suppl. Fig. S8 | ☐ | |
| Type, make and manufacturer of the measurement instrument | Manuscript pages 12-13, Fig. 2, Table 3, | ☐ | |
| Calibration status of the instrument | Manuscript pages 12-13, Fig. 2 | ☐ | ☐ |

##### A.3. Participant-level characteristics

| | | | |
|---|---|---|---|
| Ocular health and functioning | Suppl. Table S1 | ☐ | |
| Pupil size and/or dilation | | ☑ | ☐ |
| Relative time (e.g. to circadian phase or sleep) | Fig. 1, manuscript pages 12-13 | ☐ | ☐ |

#### B. Light characteristics

##### B.1. Light source type(s). Please select all that are relevant.

| Room illumination (overhead or other) | Emissive surfaces including displays (incl. light therapy devices) | Wearable light emitting glasses | Ganzfeld exposure | Other: |
|---|---|---|---|---|
| ☑ | ☐ | ☐ | ☐ | ☐ |

| Polychromatic light | Monochromatic or narrowband light |
|---|---|
| ☑ | ☐ |

| | Location (page, figure, table number) | Not available | Not applicable |
|---|---|---|---|
| Type, make and manufacturer of the light source | Manuscript page 12 | ☐ | |
| Use of wearable filtering apparatus (e.g., blue-blocking glasses) | Suppl. Fig. S6, Suppl. Table S10 | ☐ | ☐ |

##### B.2. Light level characteristics

| | | | |
|---|---|---|---|
| Illuminance (lux) and/or luminance (cd/m$^2$) | Fig. 2, Table 3, Suppl. Fig. S8, Suppl. Table S12 | ☐ | |
| Spectral irradiance and/or radiance distribution | Fig. 2, Suppl. Fig. S7 | ☐ | ☐ |
| α–opic irradiance and/or radiance (including melanopic) | Table 3, Suppl. Table S12 | ☐ | ☐ |
| α–opic equivalent daylight illuminance and/or luminance (EDI/EDL, including melanopic) | Fig. 2, Table 3, Table S12 | ☐ | ☐ |

**NOTE:** Luminance and radiance metrics (as opposed to illuminance and irradiance) are mainly relevant for emissive surfaces.

##### B.3. Colour characteristics

| | | | |
|---|---|---|---|
| Peak wavelength and bandwidth | Fig.2, Suppl. Fig. S7 | ☐ | ☐ |
| Colour appearance quantities (any) | Table 3, Table S12 | ☐ | ☐ |
| Colour rendering metrics (any) | Table 3, Table S12 | ☐ | ☐ |

**NOTE:** Peak wavelength and bandwidth are most relevant for monochromatic or narrowband light sources.

##### B.4. Temporal and spatial characteristics

| | | | |
|---|---|---|---|
| Location of stimulus and viewing distance | Manuscript page 12 | ☐ | |
| Temporal pattern (including flash frequency and waveform) | Fig. 1 | ☐ | ☐ |
| Relative or absolute size of the stimulus | Manuscript page 12 | ☐ | ☐ |
